# An efficient plasmid-based system for the recovery of recombinant vesicular stomatitis virus encoding foreign glycoproteins

**DOI:** 10.1101/2023.12.14.571669

**Authors:** María-Carmen Marqués, Iván Andreu-Moreno, Rafael Sanjuán, Santiago F. Elena, Ron Geller

## Abstract

Viral glycoproteins mediate entry into host cells, thereby dictating host range and pathogenesis. In addition, they constitute the principal target of neutralizing antibody responses, making them important antigens in vaccine development. Recombinant vesicular stomatitis virus (VSV) encoding foreign glycoproteins can provide a convenient and safe surrogate system to interrogate the function, evolution, and antigenicity of viral glycoproteins from viruses that are difficult to manipulate or those requiring high biosafety levels containment. However, the production of recombinant VSV can be technically challenging. In this work, we present an efficient and robust plasmid-based system for the production of recombinant VSV encoding foreign glycoproteins. We validate the system using glycoproteins from different viral families, including arenaviruses, coronaviruses, and hantaviruses, as well as highlight their utility for studying the effects of mutations on viral fitness. Overall, the methods described herein can facilitate the study of both native and recombinant VSV encoding foreign glycoproteins and can serve as the basis for the production of VSV-based vaccines.

## Introduction

The membranes of all enveloped viruses are decorated with one or more viral glycoproteins (GPs), whose principal task is to bind host receptors to facilitate viral entry into cells. As such, GPs dictate host and cellular tropism, providing a direct link to viral emergence and pathogenesis. Viral GPs are also highly visible to the immune system and comprise the principal targets of neutralizing antibodies ^1–3^. Consequently, these show high rates of evolution due to pressures of escaping host immunity ^4–6^. Vaccines inducing immunity to viral GPs have proven effective in reducing disease burden for multiple viruses, highlighting their potential as therapeutic targets ^1,3,7^. Hence, methods that facilitate the study of viral GP function can help illuminate basic processes of viral infection, evolution, or immunity and facilitate the development of novel therapies.

Studying the GPs of high biosafety level viruses or those that lack efficient infectious clone systems is challenging and costly. Pseudotyped viral systems, where a virus lacking its native GP is produced in cells expressing a GP of interest, can be used to generate viral particles decorated with foreign GPs. Such particles can perform a single round of infection utilizing the foreign GP for entry and have been shown to faithfully mimic viral entry processes for numerous viruses (*e*.*g*., arenaviruses, coronaviruses, filoviruses, or paramyxoviruses) ^8–12^. As such, these pseudotyped viral systems are useful for measuring neutralizing antibody titers and deciphering entry mechanisms. The most common viruses for pseudotyping have been lentiviruses and vesicular stomatitis virus (VSV) due to their ability to non-specifically acquire foreign GPs expressed on the plasma membrane during their budding process. However, as the viral GP is not encoded in the genome of these pseudotyped virus systems, they are not suitable for evaluating viral fitness. Similarly, they do not allow for harnessing the high mutation rates of the pseudotyped virus to interrogate mechanisms of adaptation, drug resistance, or escape from antibody neutralization. To overcome these limitations, experimental systems have been developed in which the GP of interest is encoded within the genome of the pseudotyped virus. As lentivirus genomes can integrate into the host genome, posing an unacceptable biosafety risk, replication-competent pseudotyped virus systems have focused on VSV ^13^, a biosafety level 2 virus that causes only mild infection in humans ^14,15^. Moreover, the VSV GP (VSV G) shows an extremely broad tropism in cell culture, making its replacement with a foreign GP unlikely to expand tropism. Indeed, recombinant VSV-based vaccines have been evaluated in clinical trials ^16–18^, leading to the approval of a VSV-based vaccine against Ebola virus, proving the safety of VSV-based vectors in humans. In addition, recombinant VSV encoding foreign proteins have been evaluated for oncolytic therapy with good safety profiles ^14^.

VSV is a bullet-shaped, negative-strand rhabdovirus with a genome of ∼11 kb. It encodes only five genes (in 3’ to 5’ order): the nucleocapsid (N), which packages the viral genome, protecting it from degradation and host antiviral effectors; a phosphoprotein (P) that is an essential cofactor of the replication complex; the matrix protein (M), which coordinates the packing of viral nucleoproteins into viral particles and blocks cellular antiviral mechanisms; the viral GP (G), which mediates receptor binding and the fusion of the viral membrane with that of the host cell to facilitate entry; and the large (L) polymerase protein that carries out viral genome transcription and replication. Each of these genes is flanked by defined gene start and end sequences that form individual transcription units. As transcription always initiates at the 3’ of the genome, a gradient of gene expression is generated in infected cells due to the reduced probability of successful reinitiation at each downstream transcription unit ^19^. Of significant advantage for the production of recombinant viruses, VSV can tolerate insertions that increase its genome size by up to 40% ^20^ as well as the introduction of additional transcription units. Moreover, as negative-strand RNA viruses do not recombine frequently, inserts are stably maintained during multiple rounds of replication ^21^.

The generation of recombinant VSV has been shown to require the co-expression of the viral antigenome together with helper plasmids encoding the minimal replication components: the N, P, and L genes ^22,23^. Early systems relied on the antigenomic RNA and helper genes being driven by a T7 promoter, which was supplied by coinfection with a vaccinia virus encoding the T7 polymerase (VV-T7) ^22,23^. Using this system, recombinant VSV in which the GP has been replaced by a foreign GP from multiple viruses have been recovered, including arenaviruses, bunyaviruses, coronaviruses, filoviruses, and paramyxoviruses ^8,13,24^. However, the additional biosafety risk posed by VV, the need to isolate the recombinant VSV away from VV, and potential interference between the two viruses constitute important disadvantages of this system. Additionally, the multiplicity of infection and the purity of VV can strongly affect recovery efficiency ^25^. Finally, despite extensive experience in recovery of recombinant VSV in our laboratory ^26^, we found this system to be inefficient for generating recombinant VSV encoding GPs which do not efficiently mediate infection in the context of VSV (*e*.*g*. SARS-CoV-2 spike proteins). Alternative methods employing a cell line that encodes the T7 polymerase together with either T7-driven helper plasmids or cytomegalovirus (CMV)-driven helper plasmids have been reported ^13,25^ but their efficiency relative to the VV-based system is unclear.

In this work, we developed and optimized a system for the recovery of recombinant VSV encoding foreign GPs. The system is efficient, resulting in up to 100% recovery for some recombinant viruses, does not rely on the use of VV, can be carried out in small-scale formats, and even enables the recovery of VSV pseudotyped with GPs that are inefficient in mediating infection. In addition, we highlight the utility of these recombinant VSVs to study the effects of GP on viral fitness using two in vitro assays. The plasmids developed herein have been deposited in the European Virus Archive for their distribution to the scientific community.

## Results and Discussion

Our initial goal was to recover recombinant VSV encoding foreign GPs from different viral families to study their evolution and adaptation to different cell types. For this, we employed a plasmid encoding the VSV antigenome of the Indiana serotype that has been previously modified to encode GFP from an additional transcription unit between the G and L genes ^27,28^. This plasmid is flanked by a T7 promoter and a T7 terminator to produce the viral antigenome in cells expressing the T7 polymerase (pVSV-GFP). We further modified this plasmid by replacing the coding region of the VSV G gene with a short linker flanked by two restriction enzymes to facilitate cloning of foreign GPs (pVSVΔG-GFP-linker) and also generated a similar plasmid encoding mCherry (pVSVΔG-mCherry-linker). To recover recombinant VSV, we utilized a previously published system that has been used to successfully recover recombinant VSV by multiple groups, including ours ^26^. In this system, the T7 polymerase is expressed in BHK21 cells by infection with a vaccinia virus encoding the T7 polymerase (VV-T7) under the immediate-early promoter. Subsequently, plasmids encoding the native N, P, and L genes of VSV flanked by T7 promoter and terminator sequences are cotransfected with the VSV antigenome plasmid. Finally, an inhibitor of DNA-dependent transcription is added to block additional replication of VV-T7 after sufficient T7 polymerase has been expressed (see Material and Methods). Rather than employing standard BHK21 cells, we used a BHK21 cell line that can be induced to express VSV G (BHK21-G43) ^29^ in order to increase the titers of VSV encoding a foreign GPs and to enable the recovery of VSV lacking any GP ^13,24,30,31^. Despite multiple attempts to optimize the recovery of recombinant VSV carrying different GPs, recovery rates remained low (success rate of 0.11±0.16), were inconsistent, and some recombinant VSVs were not successfully recovered (Table S1).

The inability to obtain recombinant VSV encoding different GPs led us to seek an alternative recovery method that is safe, efficient, and robust. For this, we chose to modify a plasmid-based approach used for the recovery of paramyxoviruses ^32^ and respiratory syncytial virus ^33^. Specifically, the need for VV-T7 was bypassed by using a plasmid encoding a codon-optimized version of the T7 polymerase (pT7-opt), which was shown to improve the recovery of negative strand viruses compared to the wildtype T7 sequence ^34^. Secondly, we designed new helper plasmids to increase the expression level of the N, P, and L proteins by codon-optimizing their sequence to more closely match that of mammalian cells, which has been shown to facilitate the recovery of an unrelated negative-strand RNA virus ^33^. The new sequences increased the codon-adaptation index ^35^ of the helper genes from < 0.69 to > 0.87 relative to BHK21 cells (Table S2). These codon-optimized sequences were then introduced into a mammalian expression plasmid between a CMV promoter and an SV40 polyadenylation signal (pCMV-Nopt, pCMV-Popt, and pCMV-Lopt) to enable VV-T7 independent expression. Finally, a synthetic intron sequence was maintained in the plasmids, which has been shown to increase protein expression in mammalian cells ^36^.

To evaluate this system, we chose to use an antigenomic VSV plasmid encoding the Spike (S) gene of the SARS-CoV-2 Omicron BA4.5. variant (pVSVΔG-GFP-S2_BA4.5._). The antigenomic and helper plasmids were transfected together with the pT7-opt plasmid into BHK-G43 cells. Experiments were performed in either 24-well or 12-well plate formats to enable the testing of multiple conditions in the same experiment (Figure 1) and BHK-G43 cells were induced to express VSV G at the time of transfection to improve recovery.

**Figure 1.**
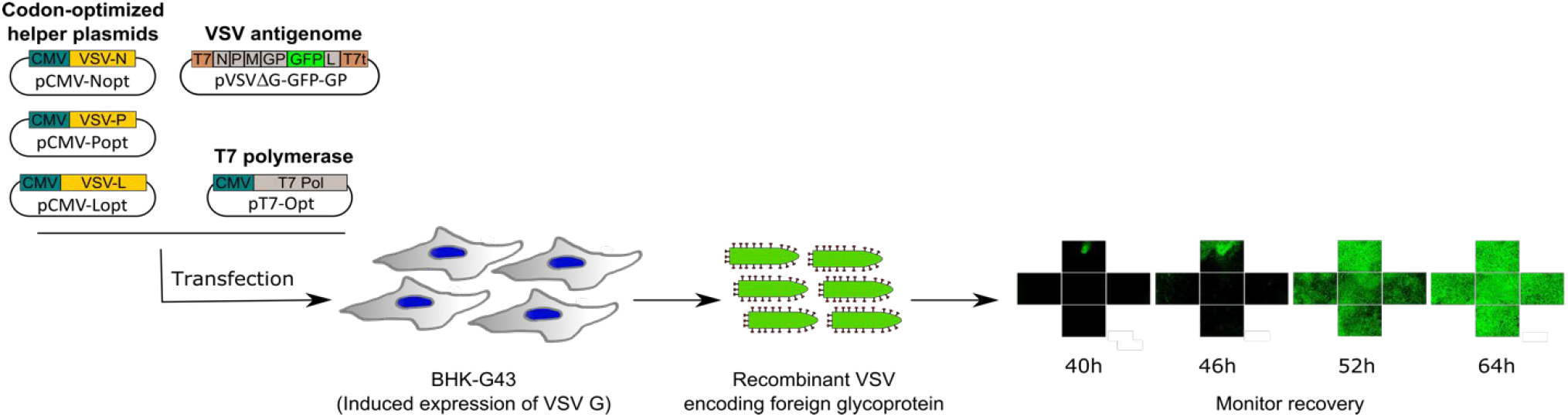
Overview of the protocol for recovery of recombinant VSV encoding foreign glycoproteins. BHK-G43 cells, which can be induced to express the VSV G protein, are utilized. To generate recombinant VSV, these cells are transfected with cytomegalovirus (CMV) immediate early promoter driven plasmids encoding codon-optimized sequences of the N, P, and L genes (helper plasmids) to provide the replication machinery, a plasmid encoding the antigenome of VSV in which the VSV glycoprotein (G) has been replaced by a foreign GP and which harbors a reporter gene (e.g. eGFP) from an additional transcription unit (pVSVΔG-GFP-GP), and a plasmid encoding a codon-optimized T7 polymerase (pT7-opt) to generate viral RNA. Virus recovery can be quantified via examination of GFP expression in transfected cells. T7, T7 promoter sequence; T7t, T7 terminator sequence.

Overall, two different transfection reagents (Lipofectamine 2000 and Lipofectamine 3000), four ratios of the helper plasmids, and three temperature conditions (33 °C, 33 °C for 40 h followed by incubation at 37 °C, or 37 °C) were evaluated to identify the best conditions for recovery. The number of successful events was monitored in a live cell microscope by examining GFP expression in each condition until ∼4 days post-infection. Recombinant virus was obtained in 11 of the 12 conditions evaluated (Figure 2). Overall, Lipofectamine 3000 resulted in improved recovery compared to Lipofectamine 2000 in all but one of the conditions (*p* = 0.002 by logistic regression), reaching 100% recovery in three of the conditions (range 17%–100%, mean 65.2%) versus a maximal success rate of 75% for Lipofectamine 2000 (range 0%–75%, mean 36.1%). Similarly, different plasmid ratios significantly altered recovery rates, with N:P:L:T7:VSVΔG-S2_BA4.5._ ratios of 3:1:1:2:1 and 2:1:1:2:1 yielding the best results (*p* < 0.0005 for both by logistic regression). No significant differences were observed between the different incubation temperatures overall (*p* > 0.05 by logistic regression), although the temperature showing the largest fraction of conditions with > 90% recovery (three of the four conditions) was observed when cells were first incubated at 33 °C for 40 h followed by incubation at 37 °C for 36 h, suggesting higher robustness for this condition.

**Figure 2.**
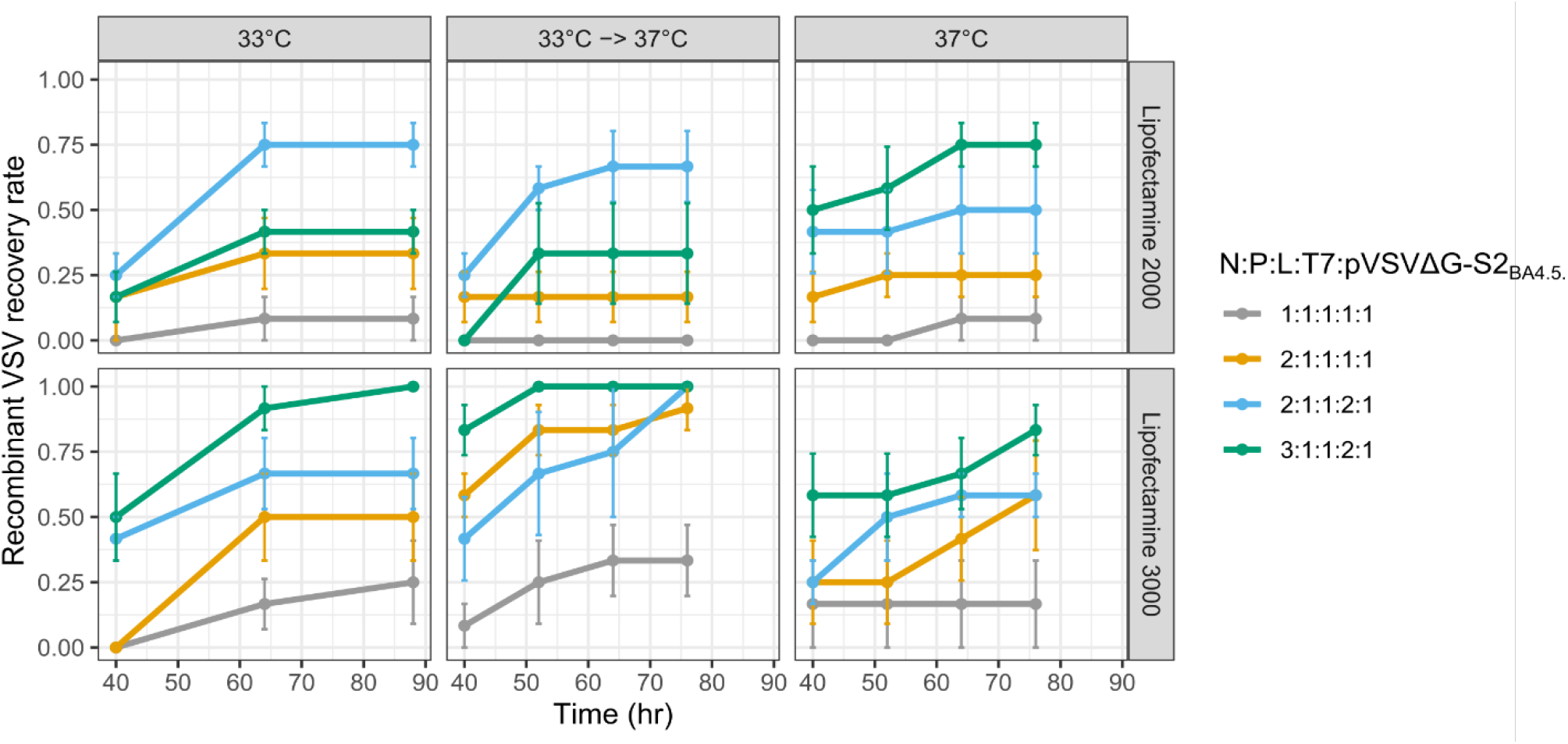
Optimization of recombinant VSV recovery. The success rate for the recovery of recombinant VSV for the indicated transfection reagent, plasmid ratios, and temperature incubation conditions. An antigenomic plasmid encoding the S gene of the SARS-CoV-2 BA4.5. variant (pVSVΔG-GFP-S2_BA4.5_) was used. GFP expression was monitored on a live-cell microscope and successful recovery was defined as those wells in which GFP fluorescence was observed at each time point. Data represents the means and SE of four biological replicates, each comprised of three technical replicates.

To validate the effect of codon optimization on the recovery of recombinant VSV, we compared the recovery rate of the antigenomic plasmid pVSVΔG-GFP-S2_BA4.5._ obtained using the codon-optimized helpers versus helpers encoding native VSV genes expressed from a standard mammalian expression plasmid (pCMV-N, pCMV-P, pCMV-L). Using the optimized conditions identified above, the recovery rate was 62.5% (5/8) using the codon-optimized helper plasmids versus 0% (0/8) for the helper plasmids encoding the native VSV sequences (*p* = 0.026 by Fisher’s exact test). Hence, codon optimization was associated with a significant increase in the recovery of recombinant VSV. Of note, a T7 promoter sequence that is commonly present in mammalian expression plasmids and could compete with the T7-mediated expression of the antigenome is present only in the helper plasmids encoding the native VSV genes and could reduce the efficiency of recovery.

Having established the conditions necessary to mediate efficient recovery of recombinant VSV, we next examined compatibility with different GPs. For this, we cloned the GPs of different arenaviruses, coronaviruses, and hantaviruses (*n* = 28), and evaluated the ability to recover recombinant VSV (Table S3). All viruses were successfully recovered, with several showing recovery rates > 75%, highlighting the robustness of this system to function in the context of divergent GPs. Of note, particles emerging from this first amplification in this system are decorated with VSV G and must be grown in cells that do not express VSV G to ensure entry via the foreign GP encoded in the viral genome.

As BHK-G43 cells are not available in all laboratories, we tested whether standard BHK21 cells can be utilized to recover recombinant VSV by supplying an additional plasmid encoding the VSV G gene. Using VSV-ΔG-GFP-S2_BA4.5._ as the antigenomic vector and a molar ratio of 2:1:1:2:1:2 for N:P:L:T7:VSVΔG-S2_BA4.5._:VSV-G plasmids, we performed four independent experiments, each comprised of four technical replicates (16 wells in total). Recombinant virus was successfully recovered in one or more wells of each replicate, with an average success rate of 37.5% (Figure 3). Similarly, when using the pVSVΔG-GFP-linker antigenomic plasmid, which does not encode a GP, recombinant virus was recovered in a single well in three of the four replicates, yielding an average recovery rate of 18.8% (Figure 3). Hence, the described protocol can be successfully implemented in standard BHK21 cells by the inclusion of a VSV G expression plasmid albeit at reduced efficiency. Additional optimization of plasmid ratios and/or the utilization of alternative cell lines that show higher transfection efficiency (*e*.*g*., HEK293T cells) could potentially increase recovery rates.

**Figure 3.**
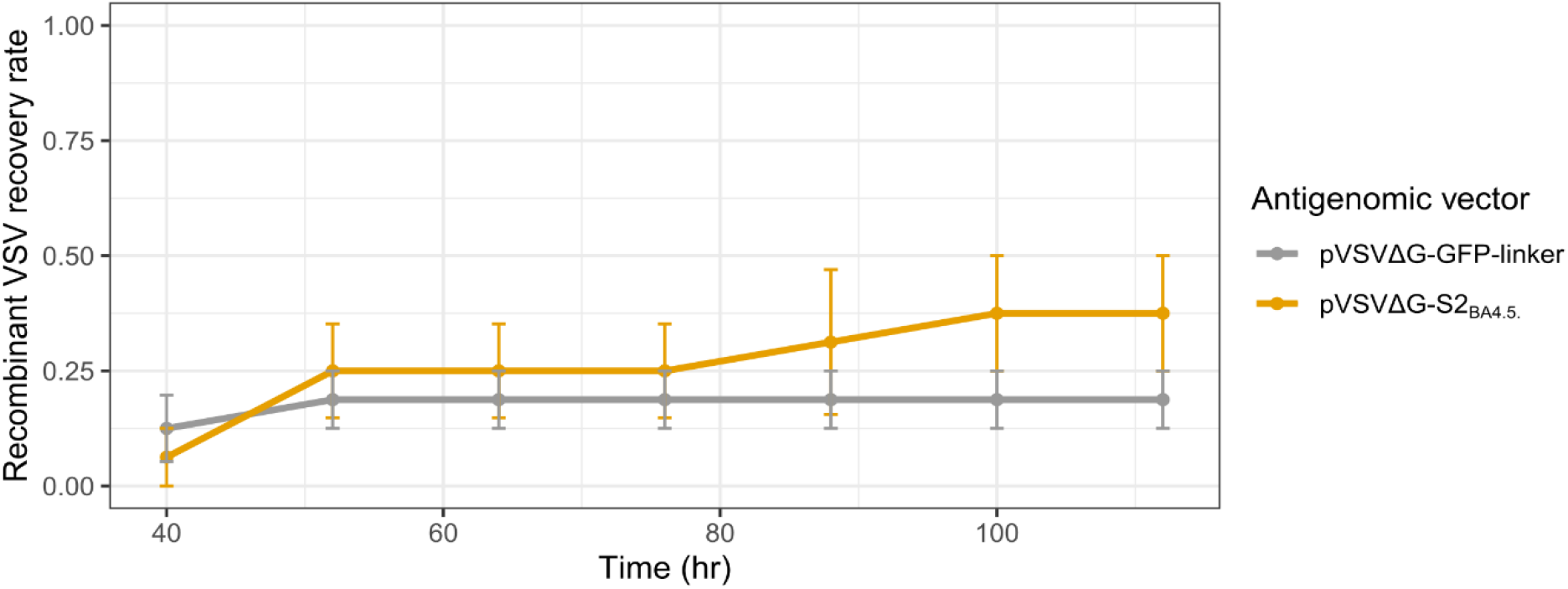
Recovery of recombinant VSV in standard BHK21 cells. The indicated antigenomic plasmid was transfected into BHK21 cells together with the plasmids N:P:L:T7:pVSVΔG.:VSV-G at a molar ratio of 2:1:1:2:1:2. Cells were incubated at 33 °C for 40 h and then transferred to 37 °C until the end of the experiment. GFP expression was monitored on a live-cell microscope and successful recovery was defined as those wells in which GFP fluorescence was observed at each time point. Data represent the mean and SE of four replicates.

Finally, to illustrate the utility of these recombinant VSVs for investigating the biology of viral GPs, we assessed the effect of VSV encoding the SARS-CoV-2 S sequences of Wuhan-Hu-1, Alpha, Delta, and Mu variants using two in vitro assays for measuring viral fitness. First, we directly competed each of these recombinant VSVs, which also encode eGFP, against a common reference VSV encoding the Wuhan-Hu-1 S gene and mCherry in lieu of eGFP. Each virus was competed against the reference virus for two sequential rounds of infection in VeroE6-TMPRSS2 cells and the amount of each virus produced at the end of the competition was compared to its relative frequency in the inoculum. Compared to VSV encoding the Wuhan-Hu-1 S, VSV encoding the Alpha S produced significantly higher viral titers (1.19 ± 0.08 fold; *p* = 0.008 by two-tailed *t*-test), VSV encoding the Delta S showed a moderate, non-significant increase (1.07 ± 0.05 fold; *p* = 0.11 by two-tailed *t*-test), while VSV encoding the Mu S produced significantly lower viral titers (0.43 ± 0.28 fold; *p* = 0.02 by two-tailed *t*-test; Figure 4A). As an alternative assay for viral fitness, we assessed the ability of recombinant VSV encoding the different S genes to spread over time. For this, VeroE6-TMPRSS2 cells were infected with a single recombinant VSV encoding the different S sequences, and viral spread was monitored by measuring virus-driven GFP expression over time using a live-cell microscope. The GFP signal was integrated using area under the curve analysis to quantify the spread of each virus over time and standardized to that of the VSV encoding the Wuhan-Hu-1 S gene in each experiment. Relative spread measurements showed good consistency between experimental replicates (*n* = 3, Pearson correlation coefficient of 0.88 – 0.97). As observed in the direct competition assay, Alpha S showed significantly increased viral spread compared to Wuhan-Hu-1 (1.66 ± 0.24 fold; *p* = 0.0009 by two-tailed *t*-test), the spread of Delta S was improved but did not reach statistical significance (1.59 ± 0.39 fold; *p* = 0.07 by two-tailed *t*-test), and Mu S showed significantly decreased spread (0.82 ± 0.58 fold; *p* = 0.008 by two-tailed *t*-test; Figure 4B). Hence, both assays provided similar results, revealing significantly increased fitness of the Alpha variant relative to Wuhan-Hu-1, a more modest improvement in fitness of the Delta variant, and reduced fitness of the Mu variant. These results are in line with findings obtained using the full infectious SARS-CoV-2 Alpha and Delta variants, where improved fitness was observed over the Wuhan-Hu-1 strain, although Delta was shown to have higher fitness than Alpha ^37,38^. Hence, the recombinant VSV system described herein provides a convenient method for assessing the effect of different GPs on viral fitness.

**Figure 4.**
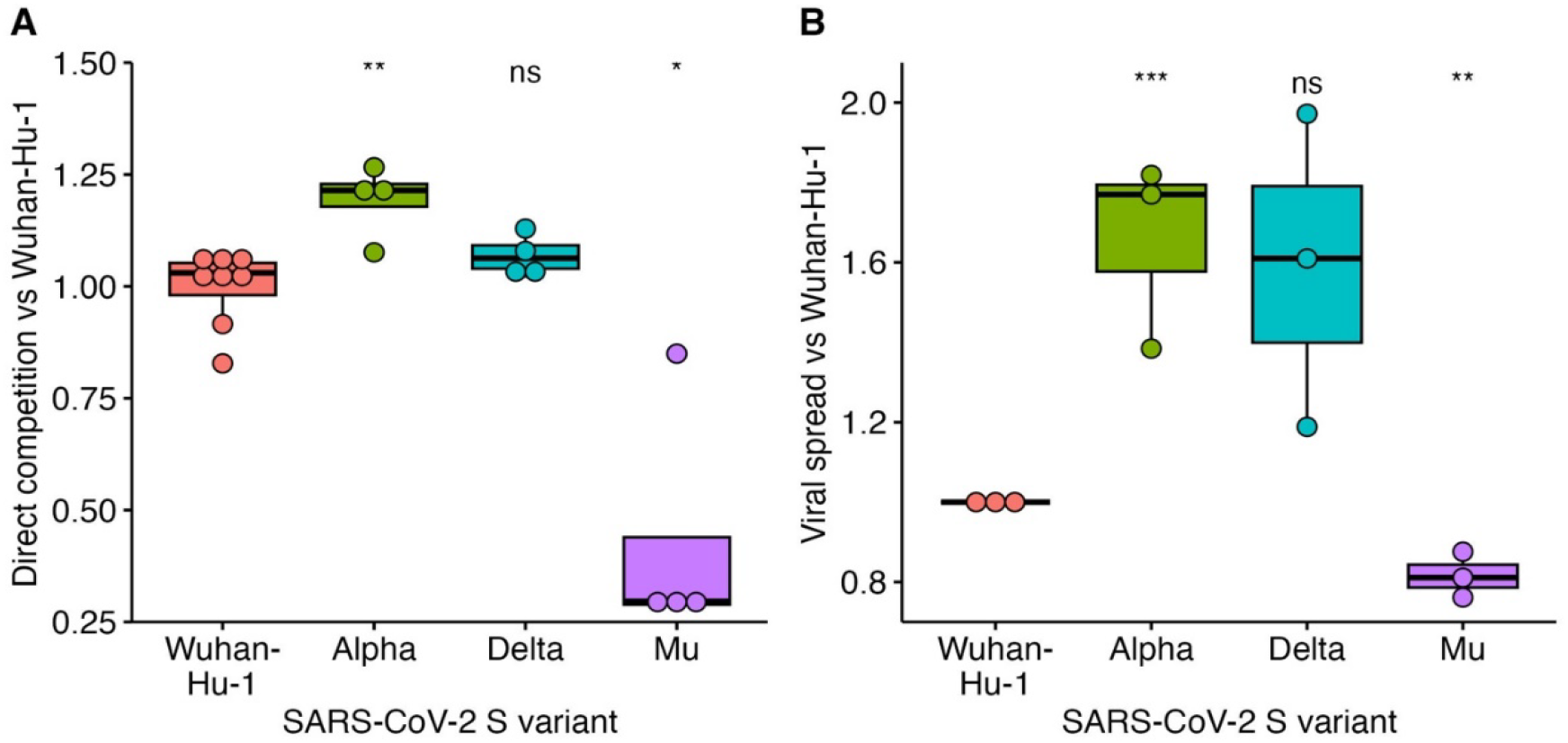
Evaluating the effect of SARS-CoV-2 S sequence on viral fitness using recombinant VSV. (**A**) Viral fitness assessment using direct competition. Recombinant VSVs encoding the indicated SARS-CoV-2 S sequence and eGFP were competed against a reference recombinant VSV encoding the Wuhan-Hu-1 S and mCherry for two sequential rounds, after which viral production of each variant and the reference was quantified. Data represents the relative fitness of each variant versus Wuhan-Hu-1 from at least four replicates. (**B**) Assessment of viral spread of recombinant VSV encoding the indicated SARS-CoV-2 S sequence. Recombinant VSVs were used to infect VeroE6-TMPRSS2 cells and viral-driven GFP expression was measured every 3 h starting at 9 h post-infection until signal saturation. The area under the curve was then calculated and standardized to that observed for recombinant VSV encoding the Wuhan-Hu-1 S gene. The data represent the results of three independent experiments, each comprising at least three technical replicates. ns, *p* > 0.05, *, *p* < 0.05; **, *p* < 0.01, ***, *p* < 0.01 by two-tailed *t*-test.

In sum, we describe an efficient, safe, and economical approach to generate recombinant VSV encoding foreign glycoproteins. We demonstrate the utility of the method by recovering a large number of recombinant VSVs encoding different GPs from multiple viral families, as well as VSV lacking a GP, which can be used for pseudotyping experiments where the GP is provided in *trans*. The recovered viruses encode a fluorescent reporter gene to facilitate the detection of successful recovery of recombinant viruses as well as their titration and can be used to interrogate adaptation to cellular environments, the identification of drug or neutralizing antibody escape mutations, fitness assessment, or the production of vaccine candidates.

## Supporting information

Data_S1_plasmid sequences

Data_S2_recovery protocol

## Acknowledgments

We would like to thank Gert Zimmer for sharing the BHK-G43 cells.

## Funding

Funding for this project was provided by grants SGL2021-03-009 and SGL2021-03-052 from European Union NextGenerationEU/PRTR through the CSIC Global Health Platform established by EU Council Regulation 2020/2094, to RG, SFE, and RS. RS and IA-M were also supported by Advanced Grant 101019724—EVADER from the European Research Council. Finally, RG was funded by grant CNS2022-135100 funded by MCIN/AEI/10.13039/501100011033 and by the European Union NextGenerationEU/PRTR.

## Author contributions

MCM, Conceptualization, Investigation, Methodology, Editing. IAM, Conceptualization, Investigation, Methodology, Editing. RS, Conceptualization, Investigation, Methodology, Editing, Resources. SFE, Conceptualization, Investigation, Methodology, Editing, Resources. RG, Conceptualization, Investigation, Methodology, Writing, Resources.

## Competing interests

The author(s) declare no competing interests.

## Data availability

All data is available upon request from RG.

## Materials and Methods

### Cell lines

All commercial cell lines were obtained from ATCC. BHK-G43 ^29^ were a kind gift of Dr. Gert Zimmer (Institute of Virology and Immunology, Switzerland) and were shared by Stefan Pohlmann (German Primate Center, Germany), VeroE6-TMPRSS2 were purchased from JCRB Cell Bank (JCRB1819). All cell lines were verified to be mycoplasma free. Cells were maintained in DMEM supplemented with 10% FBS with penicillin and streptomycin.

### Plasmids

The plasmid encoding the codon-optimized T7 polymerase was obtained from Addgene (plasmid 65974; ^34^) as was the VSV-G plasmid (pMD2.G, plasmid 12259). The plasmid encoding the VSV antigenome harboring an extra transcriptional unit between the G and L genes ^28^ was modified to encode eGFP was previously described ^27^. To generate the pVSVΔG-GFP-linker cloning plasmid, the vector was digested with restriction enzymes flanking the VSV G gene (*Mlu*I and *Xho*I) and phosphorylated, hybridized, complementary primers which generate the appropriate cohesive ends for relegation and encode necessary gene end and gene start sequences, as well as a *Pac*I restriction site, were cloned into this vector (see Table S2 for primer sequences). Foreign glycoproteins were then cloned into this vector by either digestion with *Mlu*I and *Pac*I or by seamless cloning using the HiFi NEBuilder (NEB) kit. To generate the pVSVΔG-mCherry-linker plasmid, the eGFP gene was replaced by the mCherry sequence. Of note, in some experiments, the pVSVΔG-GFP-linker harbored a deletion of amino acid 51 of the M protein, which renders VSV unable to counteract host interferon responses ^27^. This modification did not alter the efficiency of recovery (data not shown). The foreign GPs were synthesized by GeneScript and cloned into the pVSVΔG-GFP-linker plasmid in place of the linker sequence using standard molecular biology techniques. For the non-codon optimized helpers, the coding sequence of the N, P, and L gene were amplified by PCR from the pVSVΔG-GFP-linker plasmid and cloned into pCDNA3.1(+) between the *Kpn*I and *Xho*I restriction sites using standard molecular biology techniques. The correct sequence was verified by sequencing. For the codon-optimized helpers, the sequences of VSV Indiana strain P, N, and L genes were codon-optimized using the GenSmart™ Codon Optimization tool (www.genescript.com) using the human codon usage table as the reference. These sequences were then synthesized and cloned into the pIRES (Clonetech) vector, with the L gene synthesized as three fragments overlapping by 20 bp. The L gene was assembled into a pIRES vector digested *Nhe*I and *Xba*I using the NEBuilder HiFi DNA Assembly Master Mix (NEB #E2621) according to manufacturer instructions to generate pCMV-Lopt. The N and P genes were similarly cloned into the pIRES vector previously digested with *Nhe*I and *Not*I to generate pCMV-Nopt and pCMV-Popt. In all cases, the IRES sequence was eliminated in the cloning process. The sequences of all codon-optimized helper plasmids, the antigenomic, and VSV-mCherry can be found in supplementary Data 1 and are available from the European Virus Archive (https://www.european-virus-archive.com/; catalog numbers 007N-05469 – 007N-05474). All plasmids were transformed into *Escherichia coli* DH5α (NZYtech) and grown at 37 °C.

### Vaccinia virus-based recovery of recombinant VSV

BHK-G43 cells were seeded in a 24-well plate in DMEM with 10% FBS and no antibiotics until >90% confluence was achieved. Subsequently, the medium was removed and cells infected with 100 μL of VV encoding the T7 polymerase (∼10^7^ PFU/ml) at the experimentally determined optimal MOI ∼3 PFU/cell for our VV stock for 1 h at 37 °C. The virus was then removed, the cells were washed twice with PBS, and 0.2 mL of DMEM with 10% FBS and 10 nM mifepristone were added. Finally, cells were transfected with the T7-driven helper plasmids ^22^ using 2 μL Lipofectamine 2000 and the indicated DNA ratios. Following 5 h, 0.25 mL of DMEM containing 10% FBS, 20 nM mifepristone, and 100 μg/mL Ara-C (to block VV replication) were added and the cells were further incubated for 2-4 days at 37 °C.

### Plasmid-based recovery of recombinant VSV

For transfection, 1 μg total of DNA plasmids containing the helper and antigenomic plasmids at the indicated ratios were diluted into a final volume of 50 μL of Opti-MEM (GIBCO) together with 2 μL of the P3000 reagent when Lipofectamine 3000 was used. In a separate tube, 2 μL of the indicated lipofectamine reagent was diluted in Opti-MEM to a final volume of 50 μL per well. After a 5 min incubation at room temperature, both tubes were mixed gently and further incubated for 15 min at room temperature. BHK-G43 cells that were plated 24 h earlier at a density of 150,000 cells per well of a 12-well plate in DMEM supplemented with 5% FBS without antibiotics were washed with PBS and 200 μL of Opti-MEM added, followed by the addition of 100 μL of the transfection reagent. The cells were then incubated for 3 h at 37 °C, after which 1 mL of DMEM containing 10% FBS and 10 nM of mifepristone (Acros Organics) were added to induce VSV G expression. The cells were then incubated at the indicated temperature and recovery was monitored by examination of GFP expression on a live-cell microscope (Incucyte SX5; Sartorius). The optimized protocol is presented in Supplementary Data 2.

### Analysis of viral competition and spread

All viral stocks used in these experiments were amplified in VeroE6-TMPRSS2 cells and treated with a neutralizing antibody targeting G prior to infection to clear any VSV G from viral population resulting from their recovery in induced BHK-G43 cells which express VSV G. For direct competition, VeroE6-TMPRSS2 grown overnight in 24-well plates so as to reach 50% confluence were infected with 400 focus forming units (FFU) containing a 1:1 mixture of a reference VSV (encoding mCherry and the Wuhan-Hu-1 SARS-CoV-2 S sequence) and the test recombinant VSV encoding eGFP and the S sequence of interest. Following 2.5 h of infection at 37 ºC, the virus mixture was removed from the wells, the cells were washed with PBS, and 0.5 mL of DMEM containing 2% FBS was added. After 24 h, virus-containing supernatant was collected and the relative amount of each virus was quantified by plaque assay. Briefly, VeroE6-TMPRSS2 cells were infected with different dilutions of virus-containing supernatant, overlayed with DMEM containing 0.5% Agar (Sigma-Aldrich catalog A1296) to restrict viral spread, and imaged on a live-cell microscope (Incucyte SX5) after 24 h. Viral titers were then obtained by counting green or red foci. A second round of infection was similarly performed with 400 FFU/well of the viruses emerging from the first infection and quantified. The fitness, W, of each variant was calculated using the formula 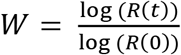 with R(0) and R(t) representing the ratio of the counts of virus production of the test virus versus the reference virus in the initial inoculation mixture or the supernatant recovered from the second round of infection. All values were standardized by the average fitness of the recombinant VSV encoding the Wuhan-Hu-1 S. For viral spread, VeroE6-TMPRSS2 cells in 12-well or 24-well well plates were infected at an MOI of 0.004 for 1.5 h. Subsequently, the media was replaced with DMEM containing 2% FBS and GFP fluorescence was monitored using a live-cell microscope starting at 9 h every 3 h for 48 h or until saturation was observed in the Wuhan-Hu-1 infected condition. The area under the curve of the log-transformed total integrated GFP fluorescence versus time was obtained using the DescTools package (version 0.99.50) in R (version 4.3.1). All values were standardized by dividing their value by the average AUC obtained for the Wuhan-Hu-1 S sequence in each experiment.

## Analysis of codon adaptation

The original and codon-optimized sequences were analyzed using the Codon Adaptation Index tool ^39^ (http://genomes.urv.es/CAIcal/) using the codon usage table of *Mesocricetus auratus* obtained from the Codon Usage Database (https://www.kazusa.or.jp/codon/cgi-bin/showcodon.cgi?species=10036).

## Notes

### Competing Interest Statement

The authors have declared no competing interest.

### Summary of Updates

Updated data on successful recovery of SARS-CoV-2 omicron BA4.5 pseudotype

## References

1. Schleiss, M. R. Viral Vaccines. in Encyclopedia of Infection and Immunity vol. 4 (2022).

2. Murin, C. D., Wilson, I. A. & Ward, A. B. Antibody responses to viral infections: a structural perspective across three different enveloped viruses. Nature Microbiology vol. 4 Preprint at 10.1038/s41564-019-0392-y (2019).

3. Burton, D. R. Antibodies, viruses and vaccines. Nature Reviews Immunology vol. 2 Preprint at 10.1038/nri891 (2002).

4. Corti, D. & Lanzavecchia, A. Broadly Neutralizing Antiviral Antibodies. Annu Rev Immunol 31, 705–742 (2013).

5. Cuevas, J. M., Geller, R., Garijo, R., López-Aldeguer, J. & Sanjuán, R. Extremely High Mutation Rate of HIV-1 In Vivo. PLoS Biol 13, e1002251 (2015).

6. Geller, R. et al.Highly heterogeneous mutation rates in the hepatitis C virus genome. Nat Microbiol 1, 16045 (2016).

7. Huang, D. B., Wu, J. J. & Tyring, S. K. A review of licensed viral vaccines, some of their safety concerns, and the advances in the development of investigational viral vaccines. Journal of Infection vol. 49 Preprint at 10.1016/j.jinf.2004.05.018 (2004).

8. Munis, A. M., Bentley, E. M. & Takeuchi, Y. A tool with many applications: vesicular stomatitis virus in research and medicine. Expert Opinion on Biological Therapy 1187–1201 Preprint at 10.1080/14712598.2020.1787981 (2020).

9. Salazar-García, M. et al.Pseudotyped Vesicular Stomatitis Virus-Severe Acute Respiratory Syndrome-Coronavirus-2 Spike for the Study of Variants, Vaccines, and Therapeutics Against Coronavirus Disease 2019. Frontiers in Microbiology vol. 12 Preprint at 10.3389/fmicb.2021.817200 (2022).

10. Cui, Q. & Huang, W. Application of Pseudotyped Viruses. Adv Exp Med Biol 1407, 45–60 (2023).

11. Duvergé, A.M &, M. Pseudotyping lentiviral vectors: When the clothes make the virus. Viruses vol. 12 Preprint at 10.3390/v12111311 (2020).

12. Whitt, M. A. Generation of VSV pseudotypes using recombinant ΔG-VSV for studies on virus entry, identification of entry inhibitors, and immune responses to vaccines. J Virol Methods 169, 365–374 (2010).

13. Yuan Fei and Zheng, A. Replicating-Competent VSV-Vectored Pseudotyped Viruses. in Pseudotyped Viruses (ed. Wang, Y.) 329–348 (Springer Nature Singapore, Singapore, 2023). doi:10.1007/978-981-99-0113-5_18.

14. Zhang, Y. & Nagalo, B. M. Immunovirotherapy Based on Recombinant Vesicular Stomatitis Virus: Where Are We? Frontiers in Immunology vol. 13 Preprint at 10.3389/fimmu.2022.898631 (2022).

15. Liu, G. et al.Vesicular stomatitis virus: From agricultural pathogen to vaccine vector. Pathogens vol. 10 Preprint at 10.3390/pathogens10091092 (2021).

16. Huttner, A. et al.The effect of dose on the safety and immunogenicity of the VSV Ebola candidate vaccine. Lancet Infect Dis 15, (2015).

17. Henao-Restrepo, A. M. et al.Efficacy and effectiveness of an rVSV-vectored vaccine in preventing Ebola virus disease: final results from the Guinea ring vaccination, open-label, cluster-randomised trial (Ebola Ça Suffit!). The Lancet 389, (2017).

18. Fuchs, J. et al.First-in-human phase I clinical trial of a recombinant vesicular stomatitis virus (rVSV)-based preventive HIV-1 vaccine. Retrovirology 9, P134 (2012).

19. Iverson, L. E. & Rose, J. K. Localized attenuation and discontinuous synthesis during vesicular stomatitis virus transcription. Cell 23, (1981).

20. Haglund, K., Forman, J., Kräusslich, H. G. & Rose, J. K. Expression of human immunodeficiency virus type 1 gag protein precursor and envelope proteins from a vesicular stomatitis virus recombinant: Highlevel production of virus-like particles containing HIV envelope. Virology 268, (2000).

21. Patiño-Galindo, J. A., Filip, I. & Rabadan, R. Global patterns of recombination across human viruses. Mol Biol Evol 38, (2021).

22. Whelan, S. P. J., Ball, L. A., Barr, J. N. & Wertz, G. T. W. Efficient recovery of infectious vesicular stomatitis virus entirely from cDNA clones. Proceedings of the National Academy of Sciences 92, 8388–8392 (1995).

23. Lawson, N. D., Stillman, E. A., Whitt, M. A. & Rose, J. K. Recombinant vesicular stomatitis viruses from DNA. Proceedings of the National Academy of Sciences 92, 4477–4481 (1995).

24. Rodriguez, S. E. et al.Vesicular Stomatitis Virus-Based Vaccine Protects Mice against Crimean-Congo Hemorrhagic Fever. Sci Rep 9, (2019).

25. Yang, F. et al.The Multiplicity of Infection of Recombinant Vaccinia Virus Expressing the T7 RNA Polymerase Determines the Rescue Efficiency of Vesicular Stomatitis Virus. Front Microbiol 13, (2022).

26. Sanjuán, R., Moya, A. & Elena, S. F. The distribution of fitness effects caused by singlenucleotide substitutions in an RNA virus. Proc Natl Acad Sci U S A 101, 8396–401 (2004).

27. Andreu-Moreno, I. & Sanjuán, R. Collective Infection of Cells by Viral Aggregates Promotes Early Viral Proliferation and Reveals a Cellular-Level Allee Effect. Curr Biol 28, 3212-3219.e4 (2018).

28. Schnell, M. J., Buonocore, L., Whitt, M. A. & Rose, J. K. The minimal conserved transcription stop-start signal promotes stable expression of a foreign gene in vesicular stomatitis virus. J Virol 70, 2318–2323 (1996).

29. Hanika, A. et al.Use of influenza C virus glycoprotein HEF for generation of vesicular stomatitis virus pseudotypes. Journal of General Virology 86, 1455–1465 (2005).

30. Dieterle, M. E. et al.A Replication-Competent Vesicular Stomatitis Virus for Studies of SARS-CoV-2 Spike-Mediated Cell Entry and Its Inhibition. Cell Host Microbe 1–11 (2020) doi:10.1016/j.chom.2020.06.020.

31. Case, J. B. et al.Neutralizing Antibody and Soluble ACE2 Inhibition of a Replication-Competent VSV-SARS-CoV-2 and a Clinical Isolate of SARS-CoV-2. Cell Host Microbe 1–11 (2020) doi:10.1016/j.chom.2020.06.021.

32. Beaty, S. M. et al.Efficient and Robust Paramyxoviridae Reverse Genetics Systems. mSphere 2, 1–14 (2017).

33. Hotard, A. L. et al.A stabilized respiratory syncytial virus reverse genetics system amenable to recombination-mediated mutagenesis. Virology 434, 129–136 (2012).

34. Yun, T. et al.Efficient reverse genetics reveals genetic determinants of budding and fusogenic differences between Nipah and Hendra viruses and enables real-time monitoring of viral spread in small animal models of henipavirus infection. J Virol 89, 1242–53 (2015).

35. Sharp, P. M. & Li, W. H. The codon adaptation index-a measure of directional synonymous codon usage bias, and its potential applications. Nucleic Acids Res 15, (1987).

36. Jo, B.-S. & Choi, S. S. Introns: The Functional Benefits of Introns in Genomes. Genomics Inform 13, 112–118 (2015).

37. Liu, Y. et al.Delta spike P681R mutation enhances SARS-CoV-2 fitness over Alpha variant. Cell Rep 39, (2022).

38. Ulrich, L. et al.Enhanced fitness of SARS-CoV-2 variant of concern Alpha but not Beta. Nature 602, (2022).

39. Puigbò, P., Bravo, I. G. & Garcia-Vallve, S. CAIcal: A combined set of tools to assess codon usage adaptation. Biol Direct 3, (2008).

