## Supplementary material for "An efficient plasmid-based system for the recovery of recombinant vesicular stomatitis virus encoding foreign glycoproteins": Data_S1_plasmid sequences

**Supplementary Data 1**

LOCUS	pCMV-Nopt	6691 bp DNA	circular SYN 04-NOV-2023

DEFINITION synthetic circular DNA

ACCESSION	.

VERSION	.

KEYWORDS	pCMV-Nopt

SOURCE	synthetic DNA construct

ORGANISM synthetic DNA construct

REFERENCE	1 (bases 1 to 6691)

AUTHORS	.

TITLE	Direct Submission

JOURNAL	Exported Nov 4, 2023 from SnapGene Viewer 7.0.3

https://www.snapgene.com

FEATURES	Location/Qualifiers

source	1..6691

/mol_type="other DNA"

/organism="synthetic DNA construct"

polyA_signal	47..168

/label=SV40 poly(A) signal

/note="SV40 polyadenylation signal"

rep_origin	354..809

/label=f1 ori

/note="f1 bacteriophage origin of replication; arrow

indicates direction of (+) strand synthesis"

/note="/snapgene_direction=RIGHT"

enhancer	926..1117

/label=SV40 enhancer

/note="enhancer for the SV40 early promoter (Herr, 1993)"

rep_origin	1134..1269

/label=SV40 ori

/note="SV40 origin of replication"

CDS	1334..2128

/codon_start=1

/gene="aph(3')-II (or nptII)"

/product="aminoglycoside phosphotransferase from Tn5"

/label=NeoR/KanR

/note="confers resistance to neomycin, kanamycin, and G418

(Geneticin(R))"

/translation="MIEQDGLHAGSPAAWVERLFGYDWAQQTIGCSDAAVFRLSAQGRP

VLFVKTDLSGALNELQDEAARLSWLATTGVPCAAVLDVVTEAGRDWLLLGEVPGQDLLS

SHLAPAEKVSIMADAMRRLHTLDPATCPFDHQAKHRIERARTRMEAGLVDQDDLDEEHQ

GLAPAELFARLKARMPDGEDLVVTHGDACLPNIMVENGRFSGFIDCGRLGVADRYQDIA

LATRDIAEELGGEWADRFLVLYGIAAPDSQRIAFYRLLDEFF"

polyA_signal	2192..2240

/label=poly(A) signal

/note="synthetic polyadenylation signal"

promoter	2546..2650

/gene="<i>bla</i>"

/label=AmpR promoter

CDS	2651..3511

/codon_start=1

/gene="<i>bla</i>"

/product="beta-lactamase"

/label=AmpR

/note="confers resistance to ampicillin, carbenicillin,

and related antibiotics"

/translation="MSIQHFRVALIPFFAAFCLPVFAHPETLVKVKDAEDQLGARVGYI

ELDLNSGKILESFRPEERFPMMSTFKVLLCGAVLSRIDAGQEQLGRRIHYSQNDLVEYS

PVTEKHLTDGMTVRELCSAAITMSDNTAANLLLTTIGGPKELTAFLHNMGDHVTRLDRW

EPELNEAIPNDERDTTMPVAMATTLRKLLTGELLTLASRQQLIDWMEADKVAGPLLRSA

LPAGWFIADKSGAGERGSRGIIAALGPDGKPSRIVVIYTTGSQATMDERNRQIAEIGAS

LIKHW"

rep_origin	3682..4270

/direction=RIGHT

/label=ori

/note="high-copy-number ColE1/pMB1/pBR322/pUC origin of

replication"

/note="/snapgene_direction=RIGHT"

enhancer	4481..4860

/label=CMV enhancer

/note="human cytomegalovirus immediate early enhancer"

promoter	4861..5072

/label=CMV promoter

/note="human cytomegalovirus (CMV) immediate early

promoter"

intron	5233..5365

/label=chimeric intron

/note="chimera between introns from human&nbsp;beta-globin

and immunoglobulin heavy chain genes"

regulatory	5417..5422

/label=Kozak sequence

/note="vertebrate consensus sequence for strong initiation

of translation (<a

href=""https://www.ncbi.nlm.nih.gov/pubmed/3313277""

title=""https://www.ncbi.nlm.nih.gov/pubmed/3313277"">Koza

k, 1987</a>)"

/regulatory_class="other"

misc_feature	5423..6691

/label=VSV_N

/label=nonstandard type: ORF

/note="/created_by=mcmarques"

/note="/modified_by=mcmarques"

ORIGIN

1 gcggccgctt ccctttagtg agggttaatg cttcgagcag acatgataag atacattgat

61 gagtttggac aaaccacaac tagaatgcag tgaaaaaaat gctttatttg tgaaatttgt

121 gatgctattg ctttatttgt aaccattata agctgcaata aacaagttaa caacaacaat

181 tgcattcatt ttatgtttca ggttcagggg gagatgtggg aggtttttta aagcaagtaa

241 aacctctaca aatgtggtaa aatccgataa ggatcgatcc gggctggcgt aatagcgaag

301 aggcccgcac cgatcgccct tcccaacagt tgcgcagcct gaatggcgaa tggacgcgcc

361 ctgtagcggc gcattaagcg cggcgggtgt ggtggttacg cgcagcgtga ccgctacact

421 tgccagcgcc ctagcgcccg ctcctttcgc tttcttccct tcctttctcg ccacgttcgc

481 cggctttccc cgtcaagctc taaatcgggg gctcccttta gggttccgat ttagtgcttt

541 acggcacctc gaccccaaaa aacttgatta gggtgatggt tcacgtagtg ggccatcgcc

601 ctgatagacg gtttttcgcc ctttgacgtt ggagtccacg ttctttaata gtggactctt

661 gttccaaact ggaacaacac tcaaccctat ctcggtctat tcttttgatt tataagggat

721 tttgccgatt tcggcctatt ggttaaaaaa tgagctgatt taacaaaaat ttaacgcgaa

781 ttttaacaaa atattaacgc ttacaatttc ctgatgcggt attttctcct tacgcatctg

841 tgcggtattt cacaccgcat acgcggatct gcgcagcacc atggcctgaa ataacctctg

901 aaagaggaac ttggttaggt accttctgag gcggaaagaa ccagctgtgg aatgtgtgtc

961 agttagggtg tggaaagtcc ccaggctccc cagcaggcag aagtatgcaa agcatgcatc

1021 tcaattagtc agcaaccagg tgtggaaagt ccccaggctc cccagcaggc agaagtatgc

1081 aaagcatgca tctcaattag tcagcaacca tagtcccgcc cctaactccg cccatcccgc

1141 ccctaactcc gcccagttcc gcccattctc cgccccatgg ctgactaatt ttttttattt

1201 atgcagaggc cgaggccgcc tcggcctctg agctattcca gaagtagtga ggaggctttt

1261 ttggaggcct aggcttttgc aaaaagcttg attcttctga cacaacagtc tcgaacttaa

1321 ggctagagcc accatgattg aacaagatgg attgcacgca ggttctccgg ccgcttgggt

1381 ggagaggcta ttcggctatg actgggcaca acagacaatc ggctgctctg atgccgccgt

1441 gttccggctg tcagcgcagg ggcgcccggt tctttttgtc aagaccgacc tgtccggtgc

1501 cctgaatgaa ctgcaggacg aggcagcgcg gctatcgtgg ctggccacga cgggcgttcc

1561 ttgcgcagct gtgctcgacg ttgtcactga agcgggaagg gactggctgc tattgggcga

1621 agtgccgggg caggatctcc tgtcatctca ccttgctcct gccgagaaag tatccatcat

1681 ggctgatgca atgcggcggc tgcatacgct tgatccggct acctgcccat tcgaccacca

1741 agcgaaacat cgcatcgagc gagcacgtac tcggatggaa gccggtcttg tcgatcagga

1801 tgatctggac gaagagcatc aggggctcgc gccagccgaa ctgttcgcca ggctcaaggc

1861 gcgcatgccc gacggcgagg atctcgtcgt gacccatggc gatgcctgct tgccgaatat

1921 catggtggaa aatggccgct tttctggatt catcgactgt ggccggctgg gtgtggcgga

1981 ccgctatcag gacatagcgt tggctacccg tgatattgct gaagagcttg gcggcgaatg

2041 ggctgaccgc ttcctcgtgc tttacggtat cgccgctccc gattcgcagc gcatcgcctt

2101 ctatcgcctt cttgacgagt tcttctgagc gggactctgg ggttcgaaat gaccgaccaa

2161 gcgacgccca acctgccatc acgatggccg caataaaata tctttatttt cattacatct

2221 gtgtgttggt tttttgtgtg aatcgatagc gataaggatc cgcgtatggt gcactctcag

2281 tacaatctgc tctgatgccg catagttaag ccagccccga cacccgccaa cacccgctga

2341 cgcgccctga cgggcttgtc tgctcccggc atccgcttac agacaagctg tgaccgtctc

2401 cgggagctgc atgtgtcaga ggttttcacc gtcatcaccg aaacgcgcga gacgaaaggg

2461 cctcgtgata cgcctatttt tataggttaa tgtcatgata ataatggttt cttagacgtc

2521 aggtggcact tttcggggaa atgtgcgcgg aacccctatt tgtttatttt tctaaataca

2581 ttcaaatatg tatccgctca tgagacaata accctgataa atgcttcaat aatattgaaa

2641 aaggaagagt atgagtattc aacatttccg tgtcgccctt attccctttt ttgcggcatt

2701 ttgccttcct gtttttgctc acccagaaac gctggtgaaa gtaaaagatg ctgaagatca

2761 gttgggtgca cgagtgggtt acatcgaact ggatctcaac agcggtaaga tccttgagag

2821 ttttcgcccc gaagaacgtt ttccaatgat gagcactttt aaagttctgc tatgtggcgc

2881 ggtattatcc cgtattgacg ccgggcaaga gcaactcggt cgccgcatac actattctca

2941 gaatgacttg gttgagtact caccagtcac agaaaagcat cttacggatg gcatgacagt

3001 aagagaatta tgcagtgctg ccataaccat gagtgataac actgcggcca acttacttct

3061 gacaacgatc ggaggaccga aggagctaac cgcttttttg cacaacatgg gggatcatgt

3121 aactcgcctt gatcgttggg aaccggagct gaatgaagcc ataccaaacg acgagcgtga

3181 caccacgatg cctgtagcaa tggcaacaac gttgcgcaaa ctattaactg gcgaactact

3241 tactctagct tcccggcaac aattaataga ctggatggag gcggataaag ttgcaggacc

3301 acttctgcgc tcggcccttc cggctggctg gtttattgct gataaatctg gagccggtga

3361 gcgtgggtct cgcggtatca ttgcagcact ggggccagat ggtaagccct cccgtatcgt

3421 agttatctac acgacgggga gtcaggcaac tatggatgaa cgaaatagac agatcgctga

3481 gataggtgcc tcactgatta agcattggta actgtcagac caagtttact catatatact

3541 ttagattgat ttaaaacttc atttttaatt taaaaggatc taggtgaaga tcctttttga

3601 taatctcatg accaaaatcc cttaacgtga gttttcgttc cactgagcgt cagaccccgt

3661 agaaaagatc aaaggatctt cttgagatcc tttttttctg cgcgtaatct gctgcttgca

3721 aacaaaaaaa ccaccgctac cagcggtggt ttgtttgccg gatcaagagc taccaactct

3781 ttttccgaag gtaactggct tcagcagagc gcagatacca aatactgttc ttctagtgta

3841 gccgtagtta ggccaccact tcaagaactc tgtagcaccg cctacatacc tcgctctgct

3901 aatcctgtta ccagtggctg ctgccagtgg cgataagtcg tgtcttaccg ggttggactc

3961 aagacgatag ttaccggata aggcgcagcg gtcgggctga acggggggtt cgtgcacaca

4021 gcccagcttg gagcgaacga cctacaccga actgagatac ctacagcgtg agctatgaga

4081 aagcgccacg cttcccgaag ggagaaaggc ggacaggtat ccggtaagcg gcagggtcgg

4141 aacaggagag cgcacgaggg agcttccagg gggaaacgcc tggtatcttt atagtcctgt

4201 cgggtttcgc cacctctgac ttgagcgtcg atttttgtga tgctcgtcag gggggcggag

4261 cctatggaaa aacgccagca acgcggcctt tttacggttc ctggcctttt gctggccttt

4321 tgctcacatg gctcgacaga tcttcaatat tggccattag ccatattatt cattggttat

4381 atagcataaa tcaatattgg ctattggcca ttgcatacgt tgtatctata tcataatatg

4441 tacatttata ttggctcatg tccaatatga ccgccatgtt ggcattgatt attgactagt

4501 tattaatagt aatcaattac ggggtcatta gttcatagcc catatatgga gttccgcgtt

4561 acataactta cggtaaatgg cccgcctggc tgaccgccca acgacccccg cccattgacg

4621 tcaataatga cgtatgttcc catagtaacg ccaataggga ctttccattg acgtcaatgg

4681 gtggagtatt tacggtaaac tgcccacttg gcagtacatc aagtgtatca tatgccaagt

4741 ccgcccccta ttgacgtcaa tgacggtaaa tggcccgcct ggcattatgc ccagtacatg

4801 accttacggg actttcctac ttggcagtac atctacgtat tagtcatcgc tattaccatg

4861 gtgatgcggt tttggcagta caccaatggg cgtggatagc ggtttgactc acggggattt

4921 ccaagtctcc accccattga cgtcaatggg agtttgtttt ggcaccaaaa tcaacgggac

4981 tttccaaaat gtcgtaacaa ctgcgatcgc ccgccccgtt gacgcaaatg ggcggtaggc

5041 gtgtacggtg ggaggtctat ataagcagag ctcgtttagt gaaccgtcag atcactagaa

5101 gctttattgc ggtagtttat cacagttaaa ttgctaacgc agtcagtgct tctgacacaa

5161 cagtctcgaa cttaagctgc agtgactctc ttaaggtagc cttgcagaag ttggtcgtga

5221 ggcactgggc aggtaagtat caaggttaca agacaggttt aaggagacca atagaaactg

5281 ggcttgtcga gacagagaag actcttgcgt ttctgatagg cacctattgg tcttactgac

5341 atccactttg cctttctctc cacaggtgtc cactcccagt tcaattacag ctcttaaggc

5401 tagagtactt aagctagcca ccatgagcgt gaccgtgaaa agaatcatcg ataataccgt

5461 ggtggtcccc aagctgcccg ccaacgagga ccccgtggaa tatcctgccg actacttcag

5521 aaaatccaag gaaatcccac tgtacatcaa caccacaaag agcctgtctg atctgcgcgg

5581 ttatgtgtac cagggcctta agagcggcaa cgtgtctatc atccacgtga acagctacct

5641 gtacggcgcc ctgaaggata tcagaggaaa actggataag gactggagca gcttcggcat

5701 caatatcggc aaggctggcg acaccatcgg aatcttcgac ctcgtgtccc tgaaggccct

5761 ggacggggtg ctgcctgatg gcgtgtccga cgccagcaga acctctgccg acgacaagtg

5821 gctgccactg tacctgctgg gcctgtacag agtgggcaga acacagatgc ctgaataccg

5881 gaagaaactg atggacggcc tgaccaatca atgtaaaatg atcaatgagc agtttgagcc

5941 tctggtgccc gagggccggg acatctttga tgtgtgggga aacgacagca actacaccaa

6001 gatcgtggcc gccgtggaca tgttcttcca catgtttaag aagcacgagt gcgccagctt

6061 ccggtacggc accattgtga gcagattcaa ggactgcgcc gctctggcca ccttcggcca

6121 cctgtgcaag atcaccggca tgagcaccga ggacgtgaca acctggatcc tgaacagaga

6181 agtggccgac gagatggtcc agatgatgct gcctggccag gagatcgaca aggccgacag

6241 ctacatgccc tacctgatcg acttcggcct atcttctaag agcccataca gcagcgtgaa

6301 aaaccccgct tttcatttct ggggccagct gacagccctg ctgctgaggt ccacaagagc

6361 cagaaacgcc agacagcctg acgacatcga gtacacaagc ctgaccaccg ccggcctgct

6421 gtatgcatac gccgtcggat ctagcgccga tctggctcag cagttctgcg tgggcgataa

6481 caagtacaca cctgacgata gcacaggcgg cctgaccacc aacgcccctc ctcagggcag

6541 agatgttgtg gaatggctgg gctggttcga ggatcagaac cggaaaccta cccctgacat

6601 gatgcaatac gctaagcggg ccgtgatgag cctgcaggga ctgcgggaaa agactattgg

6661 aaagtacgcc aagagcgagt tcgataagtg a

//

LOCUS	pCMV-Popt	6220 bp DNA	circular SYN 04-NOV-2023

DEFINITION synthetic circular DNA

ACCESSION	.

VERSION	.

KEYWORDS	pCMV-Popt

SOURCE	synthetic DNA construct

ORGANISM synthetic DNA construct

REFERENCE	1 (bases 1 to 6220)

AUTHORS	.

TITLE	Direct Submission

JOURNAL	Exported Nov 4, 2023 from SnapGene Viewer 7.0.3

https://www.snapgene.com

FEATURES	Location/Qualifiers

source	1..6220

/mol_type="other DNA"

/organism="synthetic DNA construct"

polyA_signal	47..168

/label=SV40 poly(A) signal

/note="SV40 polyadenylation signal"

rep_origin	354..809

/direction=RIGHT

/label=f1 ori

/note="f1 bacteriophage origin of replication; arrow

indicates direction of (+) strand synthesis"

promoter	926..1283

/label=SV40 promoter

/note="SV40 enhancer and early promoter"

enhancer	926..1117

/label=SV40 enhancer

/note="enhancer for the SV40 early promoter (Herr, 1993)"

rep_origin	1134..1269

/label=SV40 ori

/note="SV40 origin of replication"

CDS	1334..2128

/codon_start=1

/gene="aph(3')-II (or nptII)"

/product="aminoglycoside phosphotransferase from Tn5"

/label=NeoR/KanR

/note="confers resistance to neomycin, kanamycin, and G418

(Geneticin(R))"

/translation="MIEQDGLHAGSPAAWVERLFGYDWAQQTIGCSDAAVFRLSAQGRP

VLFVKTDLSGALNELQDEAARLSWLATTGVPCAAVLDVVTEAGRDWLLLGEVPGQDLLS

SHLAPAEKVSIMADAMRRLHTLDPATCPFDHQAKHRIERARTRMEAGLVDQDDLDEEHQ

GLAPAELFARLKARMPDGEDLVVTHGDACLPNIMVENGRFSGFIDCGRLGVADRYQDIA

LATRDIAEELGGEWADRFLVLYGIAAPDSQRIAFYRLLDEFF"

polyA_signal	2192..2240

/note="synthetic polyadenylation signal"

promoter	2546..2650

/gene="bla"

/label=AmpR promoter

CDS	2651..3511

/codon_start=1

/gene="bla"

/product="beta-lactamase"

/label=AmpR

/note="confers resistance to ampicillin, carbenicillin, and

related antibiotics"

/translation="MSIQHFRVALIPFFAAFCLPVFAHPETLVKVKDAEDQLGARVGYI

ELDLNSGKILESFRPEERFPMMSTFKVLLCGAVLSRIDAGQEQLGRRIHYSQNDLVEYS

PVTEKHLTDGMTVRELCSAAITMSDNTAANLLLTTIGGPKELTAFLHNMGDHVTRLDRW

EPELNEAIPNDERDTTMPVAMATTLRKLLTGELLTLASRQQLIDWMEADKVAGPLLRSA

LPAGWFIADKSGAGERGSRGIIAALGPDGKPSRIVVIYTTGSQATMDERNRQIAEIGAS

LIKHW"

rep_origin	3682..4270

/direction=RIGHT

/label=ori

/note="high-copy-number ColE1/pMB1/pBR322/pUC origin of

replication"

enhancer	4481..4860

/label=CMV enhancer

/note="human cytomegalovirus immediate early enhancer"

promoter	4861..5072

/label=CMV promoter

/note="human cytomegalovirus (CMV) immediate early

promoter"

intron	5233..5365

/label=chimeric intron

/note="chimera between introns from human&nbsp;beta-globin

and immunoglobulin heavy chain genes"

regulatory	5417..5426

/label=Kozak sequence

/note="vertebrate consensus sequence for strong initiation

of translation (Kozak, 1987)"

/regulatory_class="other"

regulatory	5417..5422

/label=Kozak sequence

/note="vertebrate consensus sequence for strong initiation

of translation (<a

href=""https://www.ncbi.nlm.nih.gov/pubmed/3313277""

title=""https://www.ncbi.nlm.nih.gov/pubmed/3313277"">Koza

k, 1987</a>)"

/regulatory_class="other"

misc_feature	5423..6220

/label=VSV_P

/label=nonstandard type: ORF

/note="/created_by=mcmarques"

/note="/modified_by=mcmarques"

ORIGIN

1 gcggccgctt ccctttagtg agggttaatg cttcgagcag acatgataag atacattgat

61 gagtttggac aaaccacaac tagaatgcag tgaaaaaaat gctttatttg tgaaatttgt

121 gatgctattg ctttatttgt aaccattata agctgcaata aacaagttaa caacaacaat

181 tgcattcatt ttatgtttca ggttcagggg gagatgtggg aggtttttta aagcaagtaa

241 aacctctaca aatgtggtaa aatccgataa ggatcgatcc gggctggcgt aatagcgaag

301 aggcccgcac cgatcgccct tcccaacagt tgcgcagcct gaatggcgaa tggacgcgcc

361 ctgtagcggc gcattaagcg cggcgggtgt ggtggttacg cgcagcgtga ccgctacact

421 tgccagcgcc ctagcgcccg ctcctttcgc tttcttccct tcctttctcg ccacgttcgc

481 cggctttccc cgtcaagctc taaatcgggg gctcccttta gggttccgat ttagtgcttt

541 acggcacctc gaccccaaaa aacttgatta gggtgatggt tcacgtagtg ggccatcgcc

601 ctgatagacg gtttttcgcc ctttgacgtt ggagtccacg ttctttaata gtggactctt

661 gttccaaact ggaacaacac tcaaccctat ctcggtctat tcttttgatt tataagggat

721 tttgccgatt tcggcctatt ggttaaaaaa tgagctgatt taacaaaaat ttaacgcgaa

781 ttttaacaaa atattaacgc ttacaatttc ctgatgcggt attttctcct tacgcatctg

841 tgcggtattt cacaccgcat acgcggatct gcgcagcacc atggcctgaa ataacctctg

901 aaagaggaac ttggttaggt accttctgag gcggaaagaa ccagctgtgg aatgtgtgtc

961 agttagggtg tggaaagtcc ccaggctccc cagcaggcag aagtatgcaa agcatgcatc

1021 tcaattagtc agcaaccagg tgtggaaagt ccccaggctc cccagcaggc agaagtatgc

1081 aaagcatgca tctcaattag tcagcaacca tagtcccgcc cctaactccg cccatcccgc

1141 ccctaactcc gcccagttcc gcccattctc cgccccatgg ctgactaatt ttttttattt

1201 atgcagaggc cgaggccgcc tcggcctctg agctattcca gaagtagtga ggaggctttt

1261 ttggaggcct aggcttttgc aaaaagcttg attcttctga cacaacagtc tcgaacttaa

1321 ggctagagcc accatgattg aacaagatgg attgcacgca ggttctccgg ccgcttgggt

1381 ggagaggcta ttcggctatg actgggcaca acagacaatc ggctgctctg atgccgccgt

1441 gttccggctg tcagcgcagg ggcgcccggt tctttttgtc aagaccgacc tgtccggtgc

1501 cctgaatgaa ctgcaggacg aggcagcgcg gctatcgtgg ctggccacga cgggcgttcc

1561 ttgcgcagct gtgctcgacg ttgtcactga agcgggaagg gactggctgc tattgggcga

1621 agtgccgggg caggatctcc tgtcatctca ccttgctcct gccgagaaag tatccatcat

1681 ggctgatgca atgcggcggc tgcatacgct tgatccggct acctgcccat tcgaccacca

1741 agcgaaacat cgcatcgagc gagcacgtac tcggatggaa gccggtcttg tcgatcagga

1801 tgatctggac gaagagcatc aggggctcgc gccagccgaa ctgttcgcca ggctcaaggc

1861 gcgcatgccc gacggcgagg atctcgtcgt gacccatggc gatgcctgct tgccgaatat

1921 catggtggaa aatggccgct tttctggatt catcgactgt ggccggctgg gtgtggcgga

1981 ccgctatcag gacatagcgt tggctacccg tgatattgct gaagagcttg gcggcgaatg

2041 ggctgaccgc ttcctcgtgc tttacggtat cgccgctccc gattcgcagc gcatcgcctt

2101 ctatcgcctt cttgacgagt tcttctgagc gggactctgg ggttcgaaat gaccgaccaa

2161 gcgacgccca acctgccatc acgatggccg caataaaata tctttatttt cattacatct

2221 gtgtgttggt tttttgtgtg aatcgatagc gataaggatc cgcgtatggt gcactctcag

2281 tacaatctgc tctgatgccg catagttaag ccagccccga cacccgccaa cacccgctga

2341 cgcgccctga cgggcttgtc tgctcccggc atccgcttac agacaagctg tgaccgtctc

2401 cgggagctgc atgtgtcaga ggttttcacc gtcatcaccg aaacgcgcga gacgaaaggg

2461 cctcgtgata cgcctatttt tataggttaa tgtcatgata ataatggttt cttagacgtc

2521 aggtggcact tttcggggaa atgtgcgcgg aacccctatt tgtttatttt tctaaataca

2581 ttcaaatatg tatccgctca tgagacaata accctgataa atgcttcaat aatattgaaa

2641 aaggaagagt atgagtattc aacatttccg tgtcgccctt attccctttt ttgcggcatt

2701 ttgccttcct gtttttgctc acccagaaac gctggtgaaa gtaaaagatg ctgaagatca

2761 gttgggtgca cgagtgggtt acatcgaact ggatctcaac agcggtaaga tccttgagag

2821 ttttcgcccc gaagaacgtt ttccaatgat gagcactttt aaagttctgc tatgtggcgc

2881 ggtattatcc cgtattgacg ccgggcaaga gcaactcggt cgccgcatac actattctca

2941 gaatgacttg gttgagtact caccagtcac agaaaagcat cttacggatg gcatgacagt

3001 aagagaatta tgcagtgctg ccataaccat gagtgataac actgcggcca acttacttct

3061 gacaacgatc ggaggaccga aggagctaac cgcttttttg cacaacatgg gggatcatgt

3121 aactcgcctt gatcgttggg aaccggagct gaatgaagcc ataccaaacg acgagcgtga

3181 caccacgatg cctgtagcaa tggcaacaac gttgcgcaaa ctattaactg gcgaactact

3241 tactctagct tcccggcaac aattaataga ctggatggag gcggataaag ttgcaggacc

3301 acttctgcgc tcggcccttc cggctggctg gtttattgct gataaatctg gagccggtga

3361 gcgtgggtct cgcggtatca ttgcagcact ggggccagat ggtaagccct cccgtatcgt

3421 agttatctac acgacgggga gtcaggcaac tatggatgaa cgaaatagac agatcgctga

3481 gataggtgcc tcactgatta agcattggta actgtcagac caagtttact catatatact

3541 ttagattgat ttaaaacttc atttttaatt taaaaggatc taggtgaaga tcctttttga

3601 taatctcatg accaaaatcc cttaacgtga gttttcgttc cactgagcgt cagaccccgt

3661 agaaaagatc aaaggatctt cttgagatcc tttttttctg cgcgtaatct gctgcttgca

3721 aacaaaaaaa ccaccgctac cagcggtggt ttgtttgccg gatcaagagc taccaactct

3781 ttttccgaag gtaactggct tcagcagagc gcagatacca aatactgttc ttctagtgta

3841 gccgtagtta ggccaccact tcaagaactc tgtagcaccg cctacatacc tcgctctgct

3901 aatcctgtta ccagtggctg ctgccagtgg cgataagtcg tgtcttaccg ggttggactc

3961 aagacgatag ttaccggata aggcgcagcg gtcgggctga acggggggtt cgtgcacaca

4021 gcccagcttg gagcgaacga cctacaccga actgagatac ctacagcgtg agctatgaga

4081 aagcgccacg cttcccgaag ggagaaaggc ggacaggtat ccggtaagcg gcagggtcgg

4141 aacaggagag cgcacgaggg agcttccagg gggaaacgcc tggtatcttt atagtcctgt

4201 cgggtttcgc cacctctgac ttgagcgtcg atttttgtga tgctcgtcag gggggcggag

4261 cctatggaaa aacgccagca acgcggcctt tttacggttc ctggcctttt gctggccttt

4321 tgctcacatg gctcgacaga tcttcaatat tggccattag ccatattatt cattggttat

4381 atagcataaa tcaatattgg ctattggcca ttgcatacgt tgtatctata tcataatatg

4441 tacatttata ttggctcatg tccaatatga ccgccatgtt ggcattgatt attgactagt

4501 tattaatagt aatcaattac ggggtcatta gttcatagcc catatatgga gttccgcgtt

4561 acataactta cggtaaatgg cccgcctggc tgaccgccca acgacccccg cccattgacg

4621 tcaataatga cgtatgttcc catagtaacg ccaataggga ctttccattg acgtcaatgg

4681 gtggagtatt tacggtaaac tgcccacttg gcagtacatc aagtgtatca tatgccaagt

4741 ccgcccccta ttgacgtcaa tgacggtaaa tggcccgcct ggcattatgc ccagtacatg

4801 accttacggg actttcctac ttggcagtac atctacgtat tagtcatcgc tattaccatg

4861 gtgatgcggt tttggcagta caccaatggg cgtggatagc ggtttgactc acggggattt

4921 ccaagtctcc accccattga cgtcaatggg agtttgtttt ggcaccaaaa tcaacgggac

4981 tttccaaaat gtcgtaacaa ctgcgatcgc ccgccccgtt gacgcaaatg ggcggtaggc

5041 gtgtacggtg ggaggtctat ataagcagag ctcgtttagt gaaccgtcag atcactagaa

5101 gctttattgc ggtagtttat cacagttaaa ttgctaacgc agtcagtgct tctgacacaa

5161 cagtctcgaa cttaagctgc agtgactctc ttaaggtagc cttgcagaag ttggtcgtga

5221 ggcactgggc aggtaagtat caaggttaca agacaggttt aaggagacca atagaaactg

5281 ggcttgtcga gacagagaag actcttgcgt ttctgatagg cacctattgg tcttactgac

5341 atccactttg cctttctctc cacaggtgtc cactcccagt tcaattacag ctcttaaggc

5401 tagagtactt aagctagcca ccatggacaa cctgacaaag gtgcgggaat acctgaagtc

5461 ttatagccgg ctcgatcagg ccgtcggcga aatcgacgag attgaagccc agagagccga

5521 aaagagcaac tacgagctgt ttcaggagga cggcgttgag gaacacacca agcctagcta

5581 cttccaggcc gccgacgatt ctgacaccga gagcgagccc gagatcgagg ataaccaggg

5641 cctgtacgcc cctgatccag aggccgagca agtggaaggc ttcatccagg gacctctgga

5701 cgactacgcc gatgaagagg tggacgtggt gttcaccagc gattggaagc agcctgaact

5761 ggaaagcgac gagcatggca agaccctgag actgacaagc cctgagggcc tgtccggaga

5821 acagaaatct cagtggctga gcacaatcaa ggccgtggtg cagtccgcca agtactggaa

5881 tctggctgag tgcaccttcg aggccagcgg agagggcgtg atcatgaaag agagacagat

5941 cacccccgac gtctacaagg tgacccctgt gatgaacacc caccccagcc agagcgaggc

6001 tgtgtccgac gtgtggagcc tgtccaagac aagcatgaca tttcaaccaa agaaagccag

6061 cctgcaacct ctgaccatca gcctggacga actgttcagc agtagaggcg agttcatcag

6121 cgtgggcggc gacggcagaa tgagccacaa ggaagctatc ctgctgggcc tgcggtacaa

6181 aaagctgtac aaccaggcca gggtgaagta ctctctgtga

//

LOCUS	pCVM-Lopt	11768 bp DNA	circular UNA 04-NOV-2023

DEFINITION natural circular DNA

ACCESSION	.

VERSION	.

KEYWORDS	pCMV-Lopt

SOURCE	natural DNA sequence

ORGANISM unspecified

REFERENCE	1 (bases 1 to 11768)

AUTHORS	.

TITLE	Direct Submission

JOURNAL	Exported Nov 4, 2023 from SnapGene Viewer 7.0.3

https://www.snapgene.com

FEATURES	Location/Qualifiers

source	1..11768

/mol_type="genomic DNA"

/organism="unspecified"

polyA_signal	58..179

/label=SV40 poly

/label=SV40 poly(A) signal

/note="SV40 polyadenylation signal"

rep_origin	365..820

/label=f1 ori

/note="f1 bacteriophage origin of replication; arrow

indicates direction of (+) strand synthesis"

promoter	937..1294

/label=SV40 promoter SV40 enhancer and early promoter

/label=SV40 promoter

/note="SV40 enhancer and early promoter"

enhancer	937..1128

/label=SV40 enhancer

/note="enhancer for the SV40 early promoter (Herr, 1993)"

rep_origin	1145..1280

/label=SV40 ori

/note="SV40 origin of replication"

CDS	1345..2139

/codon_start=1

/gene="aph(3')-II (or nptII)"

/product="aminoglycoside phosphotransferase from Tn5"

/label=NeoR/KanR

/note="confers resistance to neomycin, kanamycin, and G418

(Geneticin(R))"

/translation="MIEQDGLHAGSPAAWVERLFGYDWAQQTIGCSDAAVFRLSAQGRP

VLFVKTDLSGALNELQDEAARLSWLATTGVPCAAVLDVVTEAGRDWLLLGEVPGQDLLS

SHLAPAEKVSIMADAMRRLHTLDPATCPFDHQAKHRIERARTRMEAGLVDQDDLDEEHQ

GLAPAELFARLKARMPDGEDLVVTHGDACLPNIMVENGRFSGFIDCGRLGVADRYQDIA

LATRDIAEELGGEWADRFLVLYGIAAPDSQRIAFYRLLDEFF"

polyA_signal	2203..2251

/label=poly

/label=poly(A) signal

/note="synthetic polyadenylation signal"

promoter	2557..2661

/gene="bla"

/label=AmpR promoter

CDS	2662..3522

/codon_start=1

/gene="bla"

/product="beta-lactamase"

/label=AmpR

/note="confers resistance to ampicillin, carbenicillin, and

related antibiotics"

/translation="MSIQHFRVALIPFFAAFCLPVFAHPETLVKVKDAEDQLGARVGYI

ELDLNSGKILESFRPEERFPMMSTFKVLLCGAVLSRIDAGQEQLGRRIHYSQNDLVEYS

PVTEKHLTDGMTVRELCSAAITMSDNTAANLLLTTIGGPKELTAFLHNMGDHVTRLDRW

EPELNEAIPNDERDTTMPVAMATTLRKLLTGELLTLASRQQLIDWMEADKVAGPLLRSA

LPAGWFIADKSGAGERGSRGIIAALGPDGKPSRIVVIYTTGSQATMDERNRQIAEIGAS

LIKHW"

rep_origin	3693..4281

/label=ori

/note="high-copy-number ColE1/pMB1/pBR322/pUC origin of

replication"

enhancer	4492..4871

/label=CMV enhancer human cytomegalovirus immediate ea

/label=CMV enhancer

/note="human cytomegalovirus immediate early enhancer"

promoter	4872..5083

/label=CMV promoter

/note="human cytomegalovirus (CMV) immediate early

promoter"

intron	5244..5376

/label=chimeric intron

/note="chimera between introns from human beta-globin and

immunoglobulin heavy chain genes"

regulatory	5428..5437

/label=Kozak sequence

/note="vertebrate consensus sequence for strong initiation

of translation (Kozak, 1987)"

/regulatory_class="other"

misc_feature	5428..5433

/label=Kozak sequence

/note="vertebrate consensus sequence for strong initiation

of translation

(https://www.ncbi.nlm.nih.gov/pubmed/3313277""

title=""https://www.ncbi.nlm.nih.gov/pubmed/3313277"">Koza

k, 1987)"

CDS	5434..11763

/codon_start=1

/label=Lopt

/translation="MEVHDFETDEFNDFNEDDYATREFLNPDERMTYLNHADYNLNSPL

ISDDIDNLIRKFNSLPIPSMWDSKNWDGVLEMLTSCQANPIPTSQMHKWMGSWLMSDNH

DASQGYSFLHEVDKEAEITFDVVETFIRGWGNKPIEYIKKERWTDSFKILAYLCQKFLD

LHKLTLILNAVSEVELLNLARTFKGKVRRSSHGTNICRIRVPSLGPTFISEGWAYFKKL

DILMDRNFLLMVKDVIIGRMQTVLSMVCRIDNLFSEQDIFSLLNIYRIGDKIVERQGNF

SYDLIKMVEPICNLKLMKLARESRPLVPQFPHFENHIKTSVDEGAKIDRGIRFLHDQIM

SVKTVDLTLVIYGSFRHWGHPFIDYYTGLEKLHSQVTMKKDIDVSYAKALASDLARIVL

FQQFNDHKKWFVNGDLLPHDHPFKSHVKENTWPTAAQVQDFGDKWHELPLIKCFEIPDL

LDPSIIYSDKSHSMNRSEVLKHVRMNPNTPIPSKKVLQTMLDTKATNWKEFLKEIDEKG

LDDDDLIIGLKGKERELKLAGRFFSLMSWKLREYFVITEYLIKTHFVPMFKGLTMADDL

TAVIKKMLDSSSGQGLKSYEAICIANHIDYEKWNNHQRKLSNGPVFRVMGQFLGYPSLI

ERTHEFFEKSLIYYNGRPDLMRVHNNTLINSTSQRVCWQGQEGGLEGLRQKGWSILNLL

VIQREAKIRNTAVKVLAQGDNQVICTQYKTKKSRNVVELQGALNQMVSNNEKIMTAIKI

GTGKLGLLINDDETMQSADYLNYGKIPIFRGVIRGLETKRWSRVTCVTNDQIPTCANIM

SSVSTNALTVAHFAENPINAMIQYNYFGTFARLLLMMHDPALRQSLYEVQDKIPGLHSS

TFKYAMLYLDPSIGGVSGMSLSRFLIRAFPDPVTESLSFWRFIHVHARSEHLKEMSAVF

GNPEIAKFRITHIDKLVEDPTSLNIAMGMSPANLLKTEVKKCLIESRQTIRNQVIKDAT

IYLYHEEDRLRSFLWSINPLFPRFLSEFKSGTFLGVADGLISLFQNSRTIRNSFKKKYH

RELDDLIVRSEVSSLTHLGKLHLRRGSCKMWTCSATHADTLRYKSWGRTVIGTTVPHPL

EMLGPQHRKETPCAPCNTSGFNYVSVHCPDGIHDVFSSRGPLPAYLGSKTSESTSILQP

WERESKVPLIKRATRLRDAISWFVEPDSKLAMTILSNIHSLTGEEWTKRQHGFKRTGSA

LHRFSTSRMSHGGFASQSTAALTRLMATTDTMRDLGDQNFDFLFQATLLYAQITTTVAR

DGWITSCTDHYHIACKSCLRPIEEITLDSSMDYTPPDVSHVLKTWRNGEGSWGQEIKQI

YPLEGNWKNLAPAEQSYQVGRCIGFLYGDLAYRKSTHAEDSSLFPLSIQGRIRGRGFLK

GLLDGLMRASCCQVIHRRSLAHLKRPANAVYGGLIYLIDKLSVSPPFLSLTRSGPIRDE

LETIPHKIPTSYPTSNRDMGVIVRNYFKYQCRLIEKGKYRSHYSQLWLFSDVLSIDFIG

PFSISTTLLQILYKPFLSGKDKNELRELANLSSLLRSGEGWEDIHVKFFTKDILLCPEE

IRHACKFGIAKDNNKDMSYPPWGRESRGTITTIPVYYTTTPYPKMLEMPPRIQNPLLSG

IRLGQLPTGAHYKIRSILHGMGIHYRDFLSCGDGSGGMTAALLRENVHSRGIFNSLLEL

SGSVMRGASPEPPSALETLGGDKSRCVNGETCWEYPSDLCDPRTWDYFLRLKAGLGLQI

DLIVMDMEVRDSSTSLKIETNVRNYVHRILDEQGVLIYKTYGTYICESEKNAVTILGPM

FKTVDLVQTEFSSSQTSEVYMVCKGLKKLIDEPNPDWSSINESWKNLYAFQSSEQEFAR

AKKVSTYFTLTGIPSQFIPDPFVNIETMLQIFGVPTGVSHAAALKSSDRPADLLTISLF

YMAIISYYNINHIRVGPIPPNPPSDGIAQNVGIAITGISFWLSLMEKDIPLYQQCLAVI

QQSFPIRWEAVSVKGGYKQKWSTRGDGLPKDTRISDSLAPIGNWIRSLELVRNQVRLNP

FNEILFNQLCRTVDNHLKWSNLRRNTGMIEWINRRISKEDRSILMLKSDLHEENSWRD"

ORIGIN

1 agtcgacccg ggcggccgct tccctttagt gagggttaat gcttcgagca gacatgataa

61 gatacattga tgagtttgga caaaccacaa ctagaatgca gtgaaaaaaa tgctttattt

121 gtgaaatttg tgatgctatt gctttatttg taaccattat aagctgcaat aaacaagtta

181 acaacaacaa ttgcattcat tttatgtttc aggttcaggg ggagatgtgg gaggtttttt

241 aaagcaagta aaacctctac aaatgtggta aaatccgata aggatcgatc cgggctggcg

301 taatagcgaa gaggcccgca ccgatcgccc ttcccaacag ttgcgcagcc tgaatggcga

361 atggacgcgc cctgtagcgg cgcattaagc gcggcgggtg tggtggttac gcgcagcgtg

421 accgctacac ttgccagcgc cctagcgccc gctcctttcg ctttcttccc ttcctttctc

481 gccacgttcg ccggctttcc ccgtcaagct ctaaatcggg ggctcccttt agggttccga

541 tttagtgctt tacggcacct cgaccccaaa aaacttgatt agggtgatgg ttcacgtagt

601 gggccatcgc cctgatagac ggtttttcgc cctttgacgt tggagtccac gttctttaat

661 agtggactct tgttccaaac tggaacaaca ctcaacccta tctcggtcta ttcttttgat

721 ttataaggga ttttgccgat ttcggcctat tggttaaaaa atgagctgat ttaacaaaaa

781 tttaacgcga attttaacaa aatattaacg cttacaattt cctgatgcgg tattttctcc

841 ttacgcatct gtgcggtatt tcacaccgca tacgcggatc tgcgcagcac catggcctga

901 aataacctct gaaagaggaa cttggttagg taccttctga ggcggaaaga accagctgtg

961 gaatgtgtgt cagttagggt gtggaaagtc cccaggctcc ccagcaggca gaagtatgca

1021 aagcatgcat ctcaattagt cagcaaccag gtgtggaaag tccccaggct ccccagcagg

1081 cagaagtatg caaagcatgc atctcaatta gtcagcaacc atagtcccgc ccctaactcc

1141 gcccatcccg cccctaactc cgcccagttc cgcccattct ccgccccatg gctgactaat

1201 tttttttatt tatgcagagg ccgaggccgc ctcggcctct gagctattcc agaagtagtg

1261 aggaggcttt tttggaggcc taggcttttg caaaaagctt gattcttctg acacaacagt

1321 ctcgaactta aggctagagc caccatgatt gaacaagatg gattgcacgc aggttctccg

1381 gccgcttggg tggagaggct attcggctat gactgggcac aacagacaat cggctgctct

1441 gatgccgccg tgttccggct gtcagcgcag gggcgcccgg ttctttttgt caagaccgac

1501 ctgtccggtg ccctgaatga actgcaggac gaggcagcgc ggctatcgtg gctggccacg

1561 acgggcgttc cttgcgcagc tgtgctcgac gttgtcactg aagcgggaag ggactggctg

1621 ctattgggcg aagtgccggg gcaggatctc ctgtcatctc accttgctcc tgccgagaaa

1681 gtatccatca tggctgatgc aatgcggcgg ctgcatacgc ttgatccggc tacctgccca

1741 ttcgaccacc aagcgaaaca tcgcatcgag cgagcacgta ctcggatgga agccggtctt

1801 gtcgatcagg atgatctgga cgaagagcat caggggctcg cgccagccga actgttcgcc

1861 aggctcaagg cgcgcatgcc cgacggcgag gatctcgtcg tgacccatgg cgatgcctgc

1921 ttgccgaata tcatggtgga aaatggccgc ttttctggat tcatcgactg tggccggctg

1981 ggtgtggcgg accgctatca ggacatagcg ttggctaccc gtgatattgc tgaagagctt

2041 ggcggcgaat gggctgaccg cttcctcgtg ctttacggta tcgccgctcc cgattcgcag

2101 cgcatcgcct tctatcgcct tcttgacgag ttcttctgag cgggactctg gggttcgaaa

2161 tgaccgacca agcgacgccc aacctgccat cacgatggcc gcaataaaat atctttattt

2221 tcattacatc tgtgtgttgg ttttttgtgt gaatcgatag cgataaggat ccgcgtatgg

2281 tgcactctca gtacaatctg ctctgatgcc gcatagttaa gccagccccg acacccgcca

2341 acacccgctg acgcgccctg acgggcttgt ctgctcccgg catccgctta cagacaagct

2401 gtgaccgtct ccgggagctg catgtgtcag aggttttcac cgtcatcacc gaaacgcgcg

2461 agacgaaagg gcctcgtgat acgcctattt ttataggtta atgtcatgat aataatggtt

2521 tcttagacgt caggtggcac ttttcgggga aatgtgcgcg gaacccctat ttgtttattt

2581 ttctaaatac attcaaatat gtatccgctc atgagacaat aaccctgata aatgcttcaa

2641 taatattgaa aaaggaagag tatgagtatt caacatttcc gtgtcgccct tattcccttt

2701 tttgcggcat tttgccttcc tgtttttgct cacccagaaa cgctggtgaa agtaaaagat

2761 gctgaagatc agttgggtgc acgagtgggt tacatcgaac tggatctcaa cagcggtaag

2821 atccttgaga gttttcgccc cgaagaacgt tttccaatga tgagcacttt taaagttctg

2881 ctatgtggcg cggtattatc ccgtattgac gccgggcaag agcaactcgg tcgccgcata

2941 cactattctc agaatgactt ggttgagtac tcaccagtca cagaaaagca tcttacggat

3001 ggcatgacag taagagaatt atgcagtgct gccataacca tgagtgataa cactgcggcc

3061 aacttacttc tgacaacgat cggaggaccg aaggagctaa ccgctttttt gcacaacatg

3121 ggggatcatg taactcgcct tgatcgttgg gaaccggagc tgaatgaagc cataccaaac

3181 gacgagcgtg acaccacgat gcctgtagca atggcaacaa cgttgcgcaa actattaact

3241 ggcgaactac ttactctagc ttcccggcaa caattaatag actggatgga ggcggataaa

3301 gttgcaggac cacttctgcg ctcggccctt ccggctggct ggtttattgc tgataaatct

3361 ggagccggtg agcgtgggtc tcgcggtatc attgcagcac tggggccaga tggtaagccc

3421 tcccgtatcg tagttatcta cacgacgggg agtcaggcaa ctatggatga acgaaataga

3481 cagatcgctg agataggtgc ctcactgatt aagcattggt aactgtcaga ccaagtttac

3541 tcatatatac tttagattga tttaaaactt catttttaat ttaaaaggat ctaggtgaag

3601 atcctttttg ataatctcat gaccaaaatc ccttaacgtg agttttcgtt ccactgagcg

3661 tcagaccccg tagaaaagat caaaggatct tcttgagatc ctttttttct gcgcgtaatc

3721 tgctgcttgc aaacaaaaaa accaccgcta ccagcggtgg tttgtttgcc ggatcaagag

3781 ctaccaactc tttttccgaa ggtaactggc ttcagcagag cgcagatacc aaatactgtt

3841 cttctagtgt agccgtagtt aggccaccac ttcaagaact ctgtagcacc gcctacatac

3901 ctcgctctgc taatcctgtt accagtggct gctgccagtg gcgataagtc gtgtcttacc

3961 gggttggact caagacgata gttaccggat aaggcgcagc ggtcgggctg aacggggggt

4021 tcgtgcacac agcccagctt ggagcgaacg acctacaccg aactgagata cctacagcgt

4081 gagctatgag aaagcgccac gcttcccgaa gggagaaagg cggacaggta tccggtaagc

4141 ggcagggtcg gaacaggaga gcgcacgagg gagcttccag ggggaaacgc ctggtatctt

4201 tatagtcctg tcgggtttcg ccacctctga cttgagcgtc gatttttgtg atgctcgtca

4261 ggggggcgga gcctatggaa aaacgccagc aacgcggcct ttttacggtt cctggccttt

4321 tgctggcctt ttgctcacat ggctcgacag atcttcaata ttggccatta gccatattat

4381 tcattggtta tatagcataa atcaatattg gctattggcc attgcatacg ttgtatctat

4441 atcataatat gtacatttat attggctcat gtccaatatg accgccatgt tggcattgat

4501 tattgactag ttattaatag taatcaatta cggggtcatt agttcatagc ccatatatgg

4561 agttccgcgt tacataactt acggtaaatg gcccgcctgg ctgaccgccc aacgaccccc

4621 gcccattgac gtcaataatg acgtatgttc ccatagtaac gccaataggg actttccatt

4681 gacgtcaatg ggtggagtat ttacggtaaa ctgcccactt ggcagtacat caagtgtatc

4741 atatgccaag tccgccccct attgacgtca atgacggtaa atggcccgcc tggcattatg

4801 cccagtacat gaccttacgg gactttccta cttggcagta catctacgta ttagtcatcg

4861 ctattaccat ggtgatgcgg ttttggcagt acaccaatgg gcgtggatag cggtttgact

4921 cacggggatt tccaagtctc caccccattg acgtcaatgg gagtttgttt tggcaccaaa

4981 atcaacggga ctttccaaaa tgtcgtaaca actgcgatcg cccgccccgt tgacgcaaat

5041 gggcggtagg cgtgtacggt gggaggtcta tataagcaga gctcgtttag tgaaccgtca

5101 gatcactaga agctttattg cggtagttta tcacagttaa attgctaacg cagtcagtgc

5161 ttctgacaca acagtctcga acttaagctg cagtgactct cttaaggtag ccttgcagaa

5221 gttggtcgtg aggcactggg caggtaagta tcaaggttac aagacaggtt taaggagacc

5281 aatagaaact gggcttgtcg agacagagaa gactcttgcg tttctgatag gcacctattg

5341 gtcttactga catccacttt gcctttctct ccacaggtgt ccactcccag ttcaattaca

5401 gctcttaagg ctagagtact taagctagcc accatggaag tgcacgactt cgagactgac

5461 gagtttaatg atttcaacga ggacgattac gccacaagag agttcctgaa tccagacgag

5521 cgaatgacgt acctgaacca cgctgattac aacctgaaca gccctctgat cagcgacgac

5581 atcgacaatc tgatcagaaa gttcaatagc ctgcccatac cctctatgtg ggacagcaag

5641 aactgggacg gagtcctgga aatgctcacc tcttgccagg ccaatcctat ccccacaagt

5701 cagatgcaca agtggatggg cagttggctg atgagcgaca atcacgacgc atcccagggc

5761 tacagcttcc tgcacgaggt ggacaaggaa gccgagatca ccttcgatgt ggtggaaacc

5821 ttcatcagag gctggggaaa caagcctatc gagtacatca aaaaagagag atggacagac

5881 tctttcaaga tccttgccta cctgtgccaa aagttcttgg acctgcataa gctgacactg

5941 atcctgaacg ctgtgtctga agtggaattg ctgaacctgg cgcgaacctt taaaggcaaa

6001 gtgcggagat ccagccatgg aaccaacatc tgcagaatca gagtgccttc tctcggccca

6061 acctttatca gcgaaggctg ggcctacttt aagaagctgg acatcctgat ggacaggaac

6121 ttcctgttga tggttaagga cgttatcatc ggacgcatgc agaccgtgct gagcatggtc

6181 tgcagaatcg ataacctgtt tagcgagcaa gacatcttta gcctgctgaa catttatagg

6241 atcggcgaca agatcgtcga aagacaggga aacttcagct acgacctgat caaaatggtg

6301 gagcctatct gcaacctgaa gctcatgaag ctggcccggg agagcagacc tctggtcccc

6361 cagttccctc acttcgagaa ccacatcaag acaagcgtgg acgagggcgc caagattgac

6421 cggggtatcc ggtttctgca tgatcagatc atgagcgtga agaccgtgga cctcacgctg

6481 gtgatttacg gaagcttcag gcactggggc caccctttca tcgattacta caccggactt

6541 gagaagctgc actcgcaggt cacgatgaaa aaagacatcg atgtgtctta cgccaaggct

6601 ctggcctccg acctagccag aatcgtgctg ttccagcagt tcaacgacca caagaagtgg

6661 ttcgtgaacg gcgacctgtt gccccacgac caccccttca agtcacatgt gaaagagaac

6721 acctggccta cagctgctca ggtgcaggac ttcggcgaca agtggcacga gctgcctctg

6781 attaaatgct tcgagatccc cgacctgctg gacccctcca tcatctacag cgataaatcc

6841 cactctatga accggagcga ggtgctgaag cacgtgagaa tgaaccccaa cactcccatc

6901 ccttctaaaa aggtgctgca gaccatgctg gatacaaagg ccacaaattg gaaggaattc

6961 ctgaaggaaa ttgacgagaa gggcctggac gacgatgacc tgataatcgg gctgaaaggc

7021 aaggagcgcg aactgaaact ggctggccgg ttcttctctc tcatgagctg gaaactccgg

7081 gaatacttcg tgatcaccga atatctgatc aagacccact tcgtccccat gtttaaaggc

7141 ctgaccatgg ccgatgacct gaccgccgtg atcaagaaga tgctggatag cagcagcggg

7201 cagggcctga aaagttacga agccatttgc atcgccaacc acatcgatta cgagaagtgg

7261 aacaatcatc agagaaagct gagcaacggc cctgtgtttc gggtcatggg ccagttcctg

7321 ggctatccta gcctgatcga gagaacccat gaattcttcg agaagagcct gatctattac

7381 aacggaagac ctgacctgat gcgtgtgcac aacaacacac tgataaacag caccagccaa

7441 agagtttgct ggcagggcca ggagggcggt ttggagggcc tccgccagaa gggctggtcc

7501 atcctgaacc tgctggtgat ccagagagaa gccaagatcc gaaataccgc tgttaaagtg

7561 ctggcccagg gcgataacca ggtgatctgt acccagtata agaccaagaa gagccggaat

7621 gtggtggaac tgcagggggc cctgaatcaa atggtgtcta acaacgagaa gatcatgacc

7681 gccatcaaga tcggcacagg caagctgggc ctcctgatca acgatgatga gacaatgcag

7741 tccgccgact acctcaacta cgggaagatc cctattttca gaggcgtgat ccggggcctg

7801 gaaaccaaga gatggagccg ggtcacctgt gtgaccaacg accagatccc cacctgtgcc

7861 aatatcatgt caagcgtgag cactaacgcc ctgaccgtcg cccactttgc tgagaacccc

7921 atcaacgcca tgatccagta caactacttc ggcacctttg cgcgcttact gctgatgatg

7981 cacgaccccg ccctgcggca gagcctgtac gaggtccagg ataagatccc tggcctgcat

8041 agctctacct ttaagtacgc gatgctgtac ctggacccaa gcatcggcgg cgtgtctggc

8101 atgagcttaa gcaggttcct gattagagct ttccctgatc ctgtgacaga gagcctgtca

8161 ttctggcggt tcatccacgt gcacgcgaga tccgagcacc tgaaggaaat gtccgccgtg

8221 ttcggaaacc ctgagatcgc caagttccgg attacccaca tcgataagct tgtggaagat

8281 cctactagcc tcaatattgc aatgggcatg tcgccggcaa acctgctgaa gaccgaggtg

8341 aagaagtgcc tcatcgagtc gcgccagaca atcagaaacc aggtgataaa ggatgctaca

8401 atttacctgt accacgagga ggatagactg cggtcttttc tgtggagtat caaccccctg

8461 tttcctagat ttctgtccga gttcaagagc ggcacattcc tgggcgtggc tgacggactg

8521 atctctctgt tccagaacag cagaaccatc agaaacagct ttaaaaagaa gtaccacaga

8581 gagctagatg atctgatcgt gcggtcagag gtgagctctc tgactcacct gggaaaactg

8641 cacttaagac ggggcagctg caagatgtgg acctgcagcg ccactcacgc cgataccctg

8701 agatacaaaa gctggggccg gaccgtgatc ggcacgaccg tgccccaccc tctggaaatg

8761 ctgggccctc aacacagaaa ggaaacacct tgcgcccctt gtaatacctc cggcttcaat

8821 tacgtgagtg tgcactgccc cgacgggatc cacgacgtat tctctagcag aggccctctg

8881 cctgcctacc tgggttccaa aaccagtgaa tccacaagca tcctgcaacc ttgggagaga

8941 gaaagcaagg tgcccctaat caaaagggcc acaagacttc gggatgccat ctcctggttc

9001 gtggagcccg atagcaagct ggccatgacc atcctgtcta acatccattc gcttaccggc

9061 gaagagtgga ccaagcggca gcacggcttc aaaagaacag gctcagccct gcaccggttc

9121 agcacatcaa gaatgagcca cggcggattc gccagccaaa gcactgccgc cctgacccgg

9181 ctgatggcta caaccgatac catgcgggac ctgggcgatc agaatttcga tttcttgttt

9241 caggccaccc tgctttatgc ccagatcaca acaaccgttg ccagggacgg ctggatcaca

9301 agctgcaccg accactatca catcgcctgc aagtcttgcc tgaggccaat cgaagagatt

9361 accctggaca gctctatgga ttacacacct cccgacgtgt cccacgtgct gaagacctgg

9421 cggaatgggg aaggcagctg gggccaggag atcaagcaga tctaccctct ggagggaaat

9481 tggaagaacc tggcccctgc cgaacagtcc taccaggtcg gaagatgcat cggcttcctg

9541 tacggcgacc tggcttacag aaagtccacc cacgccgaag atagcagctt attccctctg

9601 agtatccaag gtaggatccg aggtcgtgga ttccttaaag gcctgctgga cggcctgatg

9661 cgggccagct gctgccaggt gattcaccgg agatctctgg ctcacctgaa gcggcctgcc

9721 aacgccgtat acggcggcct gatctacctg atcgataagc tgagcgtgtc ccctcctttc

9781 ctatccttga cccggagcgg accaatacgg gatgaactgg aaaccatccc gcacaaaatc

9841 cctacctctt accctacatc caatagagat atgggcgtta tcgtgcggaa ctacttcaag

9901 taccagtgca gattaatcga gaagggtaaa tacagaagcc actactctca gctgtggctg

9961 ttttctgatg tgctgagcat cgattttatc ggccccttca gcatctccac cacgctgctg

10021 cagatcctgt acaagccctt cctgtctgga aaggacaaga atgagctgcg ggagctggcc

10081 aatctgagct ctctcctgag aagtggcgaa ggatgggagg acatccacgt taaattcttc

10141 accaaggaca tcctgctgtg tcctgaggaa atccggcacg cctgtaaatt tggcatcgct

10201 aaagacaaca acaaggacat gtcctaccct ccttggggca gagagagcag aggcaccatt

10261 accacaatcc ccgtgtacta caccacaacc ccttacccca agatgctgga aatgcctcct

10321 agaatccaga accctctgct ctcaggtata agactgggcc agctgccaac cggcgctcac

10381 tacaagatta ggagcatcct gcacggcatg ggcatccatt acagagactt cctatcttgc

10441 ggcgacggct ccggcggaat gactgccgcc ctgctgagag aaaacgtgca cagccggggc

10501 atcttcaact cgttactcga attaagcggc tccgtaatga gaggcgccag ccctgaacct

10561 ccaagcgccc ttgaaaccct gggcggcgat aagtcacggt gcgtgaacgg cgagacttgc

10621 tgggaatacc ctagcgactt atgcgacccc cggacctggg attacttcct gcggctcaag

10681 gcgggcttgg gcctgcagat cgatctgatt gtgatggaca tggaagtgcg agactccagc

10741 accagcctaa agatcgaaac caacgtcaga aactacgtgc acagaatcct ggacgagcag

10801 ggggtcctga tctataagac ctacggcacc tacatctgtg aatctgaaaa gaacgctgtg

10861 acgatcctgg gcccaatgtt caagacagtg gatctggtgc aaacagaatt cagctccagc

10921 cagacctcag aagtctacat ggtctgcaag ggcctcaaga agctgatcga tgaacctaac

10981 cctgactggt cttccatcaa tgaaagctgg aagaatctgt acgccttcca gagcagcgag

11041 caggagttcg ccagagcgaa aaaggtgtcc acctacttta cactgacagg catcccttcc

11101 cagtttatcc ctgacccctt cgtgaacatc gagacaatgt tgcagatctt tggagtgcca

11161 accggcgtat ctcacgctgc cgccctgaag agctcagaca gaccagccga tctgctgaca

11221 attagccttt tctacatggc catcatctct tactacaaca tcaaccacat cagggtgggc

11281 cctatcccac ctaaccctcc ttctgacggc atcgcacaga acgtcggcat cgccataacc

11341 gggatcagct tctggctgag tctgatggaa aaagacatcc ccctctatca gcagtgtctg

11401 gccgtgatcc agcagagctt ccccatccgg tgggaggccg tgagcgtgaa aggaggatac

11461 aaacagaagt ggagcacaag aggcgacggc ctgcccaagg acacaagaat ctccgactcg

11521 ctggccccta tcggaaactg gatcagatcc ctggaactcg tgagaaacca ggtgcggctc

11581 aacccgttca atgaaatcct gttcaaccag ctgtgtcgga ctgtggacaa ccacctgaag

11641 tggtcaaacc tccgcaggaa caccggaatg atcgagtgga tcaacagaag aatcagcaaa

11701 gaggacagaa gcattctgat gctgaagtcc gacctgcacg aggagaacag ctggagagac

11761 tgatctag

//

LOCUS	pVSVdG-GFP-linker	13492 bp DNA	circular SYN 04-NOV-2023

DEFINITION synthetic circular DNA

ACCESSION	.

VERSION	.

KEYWORDS	pVSV-Delta-G-GFP-linker

SOURCE	synthetic DNA construct

ORGANISM synthetic DNA construct

REFERENCE	1 (bases 1 to 13492)

AUTHORS	.

TITLE	Direct Submission

JOURNAL	Exported Nov 4, 2023 from SnapGene Viewer 7.0.3

https://www.snapgene.com

FEATURES	Location/Qualifiers

source	1..13492

/mol_type="other DNA"

/organism="synthetic DNA construct"

rep_origin	complement(33..461)

/direction=LEFT

/label=f1 ori

/note="f1 bacteriophage origin of replication; arrow

indicates direction of (+) strand synthesis"

primer_bind	603..619

/label=M13 fwd

/note="common sequencing primer, one of multiple similar

variants"

promoter	626..644

/label=T7 promoter

/note="promoter for bacteriophage T7 RNA polymerase"

CDS	708..1976

/codon_start=1

/label=N

/translation="MSVTVKRIIDNTVVVPKLPANEDPVEYPADYFRKSKEIPLYINTT

KSLSDLRGYVYQGLKSGNVSIIHVNSYLYGALKDIRGKLDKDWSSFGINIGKAGDTIGI

FDLVSLKALDGVLPDGVSDASRTSADDKWLPLYLLGLYRVGRTQMPEYRKKLMDGLTNQ

CKMINEQFEPLVPEGRDIFDVWGNDSNYTKIVAAVDMFFHMFKKHECASFRYGTIVSRF

KDCAALATFGHLCKITGMSTEDVTTWILNREVADEMVQMMLPGQEIDKADSYMPYLIDF

GLSSKSPYSSVKNPAFHFWGQLTALLLRSTRARNARQPDDIEYTSLTTAGLLYAYAVGS

SADLAQQFCVGDNKYTPDDSTGGLTTNAPPQGRDVVEWLGWFEDQNRKPTPDMMQYAKR

AVMSLQGLREKTIGKYAKSEFDK"

CDS	2040..2837

/codon_start=1

/label=P

/translation="MDNLTKVREYLKSYSRLDQAVGEIDEIEAQRAEKSNYELFQEDGV

EEHTKPSYFQAADDSDTESEPEIEDNQGLYAPDPEAEQVEGFIQGPLDDYADEEVDVVF

TSDWKQPELESDEHGKTLRLTSPEGLSGEQKSQWLSTIKAVVQSAKYWNLAECTFEASG

EGVIMKERQITPDVYKVTPVMNTHPSQSEAVSDVWSLSKTSMTFQPKKASLQPLTISLD

ELFSSRGEFISVGGDGRMSHKEAILLGLRYKKLYNQARVKYSL"

misc_feature	2840..2848

/label=GE

misc_feature	2853..2862

/label=GS

misc_feature	2894..3583

/label=Mprotein

primer_bind	3640..3664

/label=VSV_3664F

misc_feature	3680..3688

/label=GE

misc_feature	3693..3702

/label=GS

misc_feature	3712..3729

/label=LINKER

misc_feature	3746..3756

/label=GE

misc_feature	3759..3768

/label=GS

misc_feature	3777..4501

/direction=RIGHT

/note="note=""Fragment extracted from

VSV_GFP_with_genes.xdna [725 nt] : (XhoI[4656] /

NheI[5385])"

primer_bind	complement(3778..3797)

/label=VSV_rc_7552R

CDS	3781..4497

/codon_start=1

/product="enhanced GFP"

/label=EGFP

/translation="MVSKGEELFTGVVPILVELDGDVNGHKFSVSGEGEGDATYGKLTL

KFICTTGKLPVPWPTLVTTLTYGVQCFSRYPDHMKQHDFFKSAMPEGYVQERTIFFKDD

GNYKTRAEVKFEGDTLVNRIELKGIDFKEDGNILGHKLEYNYNSHNVYIMADKQKNGIK

VNFKIRHNIEDGSVQLADHYQQNTPIGDGPVLLPDNHYLSTQSALSKDPNEKRDHMVLL

EFVTAAGITLGMDELYK"

primer_bind	complement(4191..4211)

/label=GFP_410_R

misc_feature	4587..4595

/label=GE

misc_feature	4600..4609

/label=GS

misc_feature	4610..10939

/label=L

misc_feature	10969..10977

/label=GE

terminator	11192..11239

/label=T7 terminator

/note="transcription terminator for bacteriophage T7 RNA

polymerase"

promoter	complement(11307..11325)

/label=T3 promoter

/note="promoter for bacteriophage T3 RNA polymerase"

primer_bind	complement(11346..11362)

/label=M13 rev

/note="common sequencing primer, one of multiple similar

variants"

protein_bind	11370..11386

/label=lac operator

/bound_moiety="lac repressor encoded by lacI"

/note="The lac repressor binds to the lac operator to

inhibit transcription in E. coli. This inhibition can be

relieved by adding lactose or

isopropyl-beta-D-thiogalactopyranoside (IPTG)."

promoter	complement(11394..11424)

/label=lac promoter

/note="promoter for the E. coli lac operon"

protein_bind	11439..11460

/label=CAP binding site

/bound_moiety="E. coli catabolite activator protein"

/note="CAP binding activates transcription in the presence

of cAMP."

rep_origin	complement(11748..12336)

/direction=LEFT

/label=ori

/note="high-copy-number ColE1/pMB1/pBR322/pUC origin of

replication"

CDS	complement(12507..13367)

/codon_start=1

/gene="bla"

/product="beta-lactamase"

/label=AmpR

/note="confers resistance to ampicillin, carbenicillin, and

related antibiotics"

/translation="MSIQHFRVALIPFFAAFCLPVFAHPETLVKVKDAEDQLGARVGYI

ELDLNSGKILESFRPEERFPMMSTFKVLLCGAVLSRIDAGQEQLGRRIHYSQNDLVEYS

PVTEKHLTDGMTVRELCSAAITMSDNTAANLLLTTIGGPKELTAFLHNMGDHVTRLDRW

EPELNEAIPNDERDTTMPVAMATTLRKLLTGELLTLASRQQLIDWMEADKVAGPLLRSA

LPAGWFIADKSGAGERGSRGIIAALGPDGKPSRIVVIYTTGSQATMDERNRQIAEIGAS

LIKHW"

promoter	complement(13368..13472)

/gene="bla"

/label=AmpR promoter

ORIGIN

1 cacctaaatt gtaagcgtta atattttgtt aaaattcgcg ttaaattttt gttaaatcag

61 ctcatttttt aaccaatagg ccgaaatcgg caaaatccct tataaatcaa aagaatagac

121 cgagataggg ttgagtgttg ttccagtttg gaacaagagt ccactattaa agaacgtgga

181 ctccaacgtc aaagggcgaa aaaccgtcta tcagggcgat ggcccactac gtgaaccatc

241 accctaatca agttttttgg ggtcgaggtg ccgtaaagca ctaaatcgga accctaaagg

301 gagcccccga tttagagctt gacggggaaa gccggcgaac gtggcgagaa aggaagggaa

361 gaaagcgaaa ggagcgggcg ctagggcgct ggcaagtgta gcggtcacgc tgcgcgtaac

421 caccacaccc gccgcgctta atgcgccgct acagggcgcg tcccattcgc cattcaggct

481 gcgcaactgt tgggaagggc gatcggtgcg ggcctcttcg ctattacgcc agctggcgaa

541 agggggatgt gctgcaaggc gattaagttg ggtaacgcca gggttttccc agtcacgacg

601 ttgtaaaacg acggccagtg aattgtaata cgactcacta taggacgaag acaaacaaac

661 cattattatc attaaaaggc tcaggagaaa ctttaacagt aatcaaaatg tctgttacag

721 tcaagagaat cattgacaac acagtcgtag ttccaaaact tcctgcaaat gaggatccag

781 tggaataccc ggcagattac ttcagaaaat caaaggagat tcctctttac atcaatacta

841 caaaaagttt gtcagatcta agaggatatg tctaccaagg cctcaaatcc ggaaatgtat

901 caatcataca tgtcaacagc tacttgtatg gagcattaaa ggacatccgg ggtaagttgg

961 ataaagattg gtcaagtttc ggaataaaca tcgggaaagc aggggataca atcggaatat

1021 ttgaccttgt atccttgaaa gccctggacg gcgtacttcc agatggagta tcggatgctt

1081 ccagaaccag cgcagatgac aaatggttgc ctttgtatct acttggctta tacagagtgg

1141 gcagaacaca aatgcctgaa tacagaaaaa agctcatgga tgggctgaca aatcaatgca

1201 aaatgatcaa tgaacagttt gaacctcttg tgccagaagg tcgtgacatt tttgatgtgt

1261 ggggaaatga cagtaattac acaaaaattg tcgctgcagt ggacatgttc ttccacatgt

1321 tcaaaaaaca tgaatgtgcc tcgttcagat acggaactat tgtttccaga ttcaaagatt

1381 gtgctgcatt ggcaacattt ggacacctct gcaaaataac cggaatgtct acagaagatg

1441 taacgacctg gatcttgaac cgagaagttg cagatgaaat ggtccaaatg atgcttccag

1501 gccaagaaat tgacaaggcc gattcataca tgccttattt gatcgacttt ggattgtctt

1561 ctaagtctcc atattcttcc gtcaaaaacc ctgccttcca cttctggggg caattgacag

1621 ctcttctgct cagatccacc agagcaagga atgcccgaca gcctgatgac attgagtata

1681 catctcttac tacagcaggt ttgttgtacg cttatgcagt aggatcctct gccgacttgg

1741 cacaacagtt ttgtgttgga gataacaaat acactccaga tgatagtacc ggaggattga

1801 cgactaatgc accgccacaa ggcagagatg tggtcgaatg gctcggatgg tttgaagatc

1861 aaaacagaaa accgactcct gatatgatgc agtatgcgaa aagagcagtc atgtcactgc

1921 aaggcctaag agagaagaca attggcaagt atgctaagtc agaatttgac aaatgaccct

1981 ataattctca gatcacctat tatatattat gctacatatg aaaaaaacta acagatatca

2041 tggataatct cacaaaagtt cgtgagtatc tcaagtccta ttctcgtctg gatcaggcgg

2101 taggagagat agatgagatc gaagcacaac gagctgaaaa gtccaattat gagttgttcc

2161 aagaggatgg agtggaagag catactaagc cctcttattt tcaggcagca gatgattctg

2221 acacagaatc tgaaccagaa attgaagaca atcaaggctt gtatgcacca gatccagaag

2281 ctgagcaagt tgaaggcttt atacaggggc ctttagatga ctatgcagat gaggaagtgg

2341 atgttgtatt tacttcggac tggaaacagc ctgagcttga atctgacgag catggaaaga

2401 ccttacggtt gacatcgcca gagggtttaa gtggagagca gaaatcccag tggctttcga

2461 cgattaaagc agtcgtgcaa agtgccaaat actggaatct ggcagagtgc acatttgaag

2521 catcgggaga aggggtcatt atgaaggagc gccagataac tccggatgta tataaggtca

2581 ctccagtgat gaacacacat ccgtcccaat cagaagcagt atcagatgtt tggtctctct

2641 caaagacatc catgactttc caacccaaga aagcaagtct tcagcctctc accatatcct

2701 tggatgaatt gttctcatct agaggagagt tcatctctgt cggaggtgac ggacgaatgt

2761 ctcataaaga ggccatcctg ctcggcctga gatacaaaaa gttgtacaat caggcgagag

2821 tcaaatattc tctgtagact atgaaaaaaa gtaacagata tcacgatcta agtgttatcc

2881 caatccattc atcatgagtt ccttaaagaa gattctcggt ctgaagggga aaggtaagaa

2941 atctaagaaa ttagggatcg caccaccccc ttatgaagag gacactagca tggagtatgc

3001 tccgagcgct ccaattgaca aatcctattt tggagttgac gagatggaca cctatgatcc

3061 gaatcaatta agatatgaga aattcttctt tacagtgaaa atgacggtta gatctaatcg

3121 tccgttcaga acatactcag atgtggcagc cgctgtatcc cattgggatc acatgtacat

3181 cggaatggca gggaaacgtc ccttctacaa aatcttggct tttttgggtt cttctaatct

3241 aaaggccact ccagcggtat tggcagatca aggtcaacca gagtatcacg ctcactgcga

3301 aggcagggct tatttgccac ataggatggg gaagacccct cccatgctca atgtaccaga

3361 gcacttcaga agaccattca atataggtct ttacaaggga acgattgagc tcacaatgac

3421 catctacgat gatgagtcac tggaagcagc tcctatgatc tgggatcatt tcaattcttc

3481 caaattttct gatttcagag agaaggcctt aatgtttggc ctgattgtcg agaaaaaggc

3541 atctggagcg tgggtcctgg actctatcgg ccacttcaaa tgagctagtc taacttctag

3601 cttctgaaca atccccggtt tactcagtct cccctaattc cagcctctcg aacaactaat

3661 atcctgtctt ttctatccct atgaaaaaaa ctaacagaga tcgatctgtt tacgcgtcac

3721 tttaattaac tcaaatcctg ctaggtatga aaaaaactaa cagatatcac gctcgagacc

3781 atggtgagca agggcgagga gctgttcacc ggggtggtgc ccatcctggt cgagctggac

3841 ggcgacgtaa acggccacaa gttcagcgtg tccggcgagg gcgagggcga tgccacctac

3901 ggcaagctga ccctgaagtt catctgcacc accggcaagc tgcccgtgcc ctggcccacc

3961 ctcgtgacca ccctgaccta cggcgtgcag tgcttcagcc gctaccccga ccacatgaag

4021 cagcacgact tcttcaagtc cgccatgccc gaaggctacg tccaggagcg caccatcttc

4081 ttcaaggacg acggcaacta caagacccgc gccgaggtga agttcgaggg cgacaccctg

4141 gtgaaccgca tcgagctgaa gggcatcgac ttcaaggagg acggcaacat cctggggcac

4201 aagctggagt acaactacaa cagccacaac gtctatatca tggccgacaa gcagaagaac

4261 ggcatcaagg tgaacttcaa gatccgccac aacatcgagg acggcagcgt gcagctcgcc

4321 gaccactacc agcagaacac ccccatcggc gacggccccg tgctgctgcc cgacaaccac

4381 tacctgagca cccagtccgc cctgagcaaa gaccccaacg agaagcgcga tcacatggtc

4441 ctgctggagt tcgtgaccgc cgccgggatc actctcggca tggacgagct gtacaagtaa

4501 gctagccaga ttcttcatgt ttggaccaaa tcaacttgtg ataccatgct caaagaggcc

4561 tcaattatat ttgagttttt aatttttatg aaaaaaacta acagcaatca tggaagtcca

4621 cgattttgag accgacgagt tcaatgattt caatgaagat gactatgcca caagagaatt

4681 cctgaatccc gatgagcgca tgacgtactt gaatcatgct gattacaacc tgaattctcc

4741 tctaattagt gatgatattg acaatttaat caggaaattc aattctcttc caattccctc

4801 gatgtgggat agtaagaact gggatggagt tcttgagatg ttaacgtcat gtcaagccaa

4861 tcccatccca acatctcaga tgcataaatg gatgggaagt tggttaatgt ctgataatca

4921 tgatgccagt caagggtata gttttttaca tgaagtggac aaagaggcag aaataacatt

4981 tgacgtggtg gagaccttca tccgcggctg gggcaacaaa ccaattgaat acatcaaaaa

5041 ggaaagatgg actgactcat tcaaaattct cgcttatttg tgtcaaaagt ttttggactt

5101 acacaagttg acattaatct taaatgctgt ctctgaggtg gaattgctca acttggcgag

5161 gactttcaaa ggcaaagtca gaagaagttc tcatggaacg aacatatgca ggattagggt

5221 tcccagcttg ggtcctactt ttatttcaga aggatgggct tacttcaaga aacttgatat

5281 tctaatggac cgaaactttc tgttaatggt caaagatgtg attataggga ggatgcaaac

5341 ggtgctatcc atggtatgta gaatagacaa cctgttctca gagcaagaca tcttctccct

5401 tctaaatatc tacagaattg gagataaaat tgtggagagg cagggaaatt tttcttatga

5461 cttgattaaa atggtggaac cgatatgcaa cttgaagctg atgaaattag caagagaatc

5521 aaggccttta gtcccacaat tccctcattt tgaaaatcat atcaagactt ctgttgatga

5581 aggggcaaaa attgaccgag gtataagatt cctccatgat cagataatga gtgtgaaaac

5641 agtggatctc acactggtga tttatggatc gttcagacat tggggtcatc cttttataga

5701 ttattacact ggactagaaa aattacattc ccaagtaacc atgaagaaag atattgatgt

5761 gtcatatgca aaagcacttg caagtgattt agctcggatt gttctatttc aacagttcaa

5821 tgatcataaa aagtggttcg tgaatggaga cttgctccct catgatcatc cctttaaaag

5881 tcatgttaaa gaaaatacat ggcccacagc tgctcaagtt caagattttg gagataaatg

5941 gcatgaactt ccgctgatta aatgttttga aatacccgac ttactagacc catcgataat

6001 atactctgac aaaagtcatt caatgaatag gtcagaggtg ttgaaacatg tccgaatgaa

6061 tccgaacact cctatcccta gtaaaaaggt gttgcagact atgttggaca caaaggctac

6121 caattggaaa gaatttctta aagagattga tgagaagggc ttagatgatg atgatctaat

6181 tattggtctt aaaggaaagg agagggaact gaagttggca ggtagatttt tctccctaat

6241 gtcttggaaa ttgcgagaat actttgtaat taccgaatat ttgataaaga ctcatttcgt

6301 ccctatgttt aaaggcctga caatggcgga cgatctaact gcagtcatta aaaagatgtt

6361 agattcctca tccggccaag gattgaagtc atatgaggca atttgcatag ccaatcacat

6421 tgattacgaa aaatggaata accaccaaag gaagttatca aacggcccag tgttccgagt

6481 tatgggccag ttcttaggtt atccatcctt aatcgagaga actcatgaat tttttgagaa

6541 aagtcttata tactacaatg gaagaccaga cttgatgcgt gttcacaaca acacactgat

6601 caattcaacc tcccaacgag tttgttggca aggacaagag ggtggactgg aaggtctacg

6661 gcaaaaagga tggagtatcc tcaatctact ggttattcaa agagaggcta aaatcagaaa

6721 cactgctgtc aaagtcttgg cacaaggtga taatcaagtt atttgcacac agtataaaac

6781 gaagaaatcg agaaacgttg tagaattaca gggtgctctc aatcaaatgg tttctaataa

6841 tgagaaaatt atgactgcaa tcaaaatagg gacagggaag ttaggacttt tgataaatga

6901 cgatgagact atgcaatctg cagattactt gaattatgga aaaataccga ttttccgtgg

6961 agtgattaga gggttagaga ccaagagatg gtcacgagtg acttgtgtca ccaatgacca

7021 aatacccact tgtgctaata taatgagctc agtttccaca aatgctctca ccgtagctca

7081 ttttgctgag aacccaatca atgccatgat acagtacaat tattttggga catttgctag

7141 actcttgttg atgatgcatg atcctgctct tcgtcaatca ttgtatgaag ttcaagataa

7201 gataccgggc ttgcacagtt ctactttcaa atacgccatg ttgtatttgg acccttccat

7261 tggaggagtg tcgggcatgt ctttgtccag gtttttgatt agagccttcc cagatcccgt

7321 aacagaaagt ctctcattct ggagattcat ccatgtacat gctcgaagtg agcatctgaa

7381 ggagatgagt gcagtatttg gaaaccccga gatagccaag tttcgaataa ctcacataga

7441 caagctagta gaagatccaa cctctctgaa catcgctatg ggaatgagtc cagcgaactt

7501 gttaaagact gaggttaaaa aatgcttaat cgaatcaaga caaaccatca ggaaccaggt

7561 gattaaggat gcaaccatat atttgtatca tgaagaggat cggctcagaa gtttcttatg

7621 gtcaataaat cctctgttcc ctagattttt aagtgaattc aaatcaggca cttttttggg

7681 agtcgcagac gggctcatca gtctatttca aaattctcgt actattcgga actcctttaa

7741 gaaaaagtat catagggaat tggatgattt gattgtgagg agtgaggtat cctctttgac

7801 acatttaggg aaacttcatt tgagaagggg atcatgtaaa atgtggacat gttcagctac

7861 tcatgctgac acattaagat acaaatcctg gggccgtaca gttattggga caactgtacc

7921 ccatccatta gaaatgttgg gtccacaaca tcgaaaagag actccttgtg caccatgtaa

7981 cacatcaggg ttcaattatg tttctgtgca ttgtccagac gggatccatg acgtctttag

8041 ttcacgggga ccattgcctg cttatctagg gtctaaaaca tctgaatcta catctatttt

8101 gcagccttgg gaaagggaaa gcaaagtccc actgattaaa agagctacac gtcttagaga

8161 tgctatctct tggtttgttg aacccgactc taaactagca atgactatac tttctaacat

8221 ccactcttta acaggcgaag aatggaccaa aaggcagcat gggttcaaaa gaacagggtc

8281 tgcccttcat aggttttcga catctcggat gagccatggt gggttcgcat ctcagagcac

8341 tgcagcattg accaggttga tggcaactac agacaccatg agggatctgg gagatcagaa

8401 tttcgacttt ttattccaag caacgttgct ctatgctcaa attaccacca ctgttgcaag

8461 agacggatgg atcaccagtt gtacagatca ttatcatatt gcctgtaagt cctgtttgag

8521 acccatagaa gagatcaccc tggactcaag tatggactac acgcccccag atgtatccca

8581 tgtgctgaag acatggagga atggggaagg ttcgtgggga caagagataa aacagatcta

8641 tcctttagaa gggaattgga agaatttagc acctgctgag caatcctatc aagtcggcag

8701 atgtataggt tttctatatg gagacttggc gtatagaaaa tctactcatg ccgaggacag

8761 ttctctattt cctctatcta tacaaggtcg tattagaggt cgaggtttct taaaagggtt

8821 gctagacgga ttaatgagag caagttgctg ccaagtaata caccggagaa gtctggctca

8881 tttgaagagg ccggccaacg cagtgtacgg aggtttgatt tacttgattg ataaattgag

8941 tgtatcacct ccattccttt ctcttactag atcaggacct attagagacg aattagaaac

9001 gattccccac aagatcccaa cctcctatcc gacaagcaac cgtgatatgg gggtgattgt

9061 cagaaattac ttcaaatacc aatgccgtct aattgaaaag ggaaaataca gatcacatta

9121 ttcacaatta tggttattct cagatgtctt atccatagac ttcattggac cattctctat

9181 ttccaccacc ctcttgcaaa tcctatacaa gccattttta tctgggaaag ataagaatga

9241 gttgagagag ctggcaaatc tttcttcatt gctaagatca ggagaggggt gggaagacat

9301 acatgtgaaa ttcttcacca aggacatatt attgtgtcca gaggaaatca gacatgcttg

9361 caagttcggg attgctaagg ataataataa agacatgagc tatccccctt ggggaaggga

9421 atccagaggg acaattacaa caatccctgt ttattatacg accacccctt acccaaagat

9481 gctagagatg cctccaagaa tccaaaatcc cctgctgtcc ggaatcaggt tgggccaatt

9541 accaactggc gctcattata aaattcggag tatattacat ggaatgggaa tccattacag

9601 ggacttcttg agttgtggag acggctccgg agggatgact gctgcattac tacgagaaaa

9661 tgtgcatagc agaggaatat tcaatagtct gttagaatta tcagggtcag tcatgcgagg

9721 cgcctctcct gagcccccca gtgccctaga aactttagga ggagataaat cgagatgtgt

9781 aaatggtgaa acatgttggg aatatccatc tgacttatgt gacccaagga cttgggacta

9841 tttcctccga ctcaaagcag gcttggggct tcaaattgat ttaattgtaa tggatatgga

9901 agttcgggat tcttctacta gcctgaaaat tgagacgaat gttagaaatt atgtgcaccg

9961 gattttggat gagcaaggag ttttaatcta caagacttat ggaacatata tttgtgagag

10021 cgaaaagaat gcagtaacaa tccttggtcc catgttcaag acggtcgact tagttcaaac

10081 agaatttagt agttctcaaa cgtctgaagt atatatggta tgtaaaggtt tgaagaaatt

10141 aatcgatgaa cccaatcccg attggtcttc catcaatgaa tcctggaaaa acctgtacgc

10201 attccagtca tcagaacagg aatttgccag agcaaagaag gttagtacat actttacctt

10261 gacaggtatt ccctcccaat tcattcctga tccttttgta aacattgaga ctatgctaca

10321 aatattcgga gtacccacgg gtgtgtctca tgcggctgcc ttaaaatcat ctgatagacc

10381 tgcagattta ttgaccatta gcctttttta tatggcgatt atatcgtatt ataacatcaa

10441 tcatatcaga gtaggaccga tacctccgaa ccccccatca gatggaattg cacaaaatgt

10501 ggggatcgct ataactggta taagcttttg gctgagtttg atggagaaag acattccact

10561 atatcaacag tgtttagcag ttatccagca atcattcccg attaggtggg aggctgtttc

10621 agtaaaagga ggatacaagc agaagtggag tactagaggt gatgggctcc caaaagatac

10681 ccgaatttca gactccttgg ccccaatcgg gaactggatc agatctctgg aattggtccg

10741 aaaccaagtt cgtctaaatc cattcaatga gatcttgttc aatcagctat gtcgtacagt

10801 ggataatcat ttgaaatggt caaatttgcg aagaaacaca ggaatgattg aatggatcaa

10861 tagacgaatt tcaaaagaag accggtctat actgatgttg aagagtgacc tacacgagga

10921 aaactcttgg agagattaaa aaatcatgag gagactccaa actttaagta tgaaaaaaac

10981 tttgatcctt aagaccctct tgtggttttt attttttatc tggttttgtg gtcttcgtgg

11041 gtcggcatgg catctccacc tcctcgcggt ccgacctggg catccgaagg aggacgtcgt

11101 ccactcggat ggctaaggga ggggcccccg cggggctgct aacaaagccc gaaaggaagc

11161 tgagttggct gctgccaccg ctgagcaata actagcataa ccccttgggg cctctaaacg

11221 ggtcttgagg ggttttttgc tgaaaggagg aactatatcc ggatcgagac ctcgatacta

11281 gtgcggtgga gctccagctt ttgttccctt tagtgagggt taatttcgag cttggcgtaa

11341 tcatggtcat agctgtttcc tgtgtgaaat tgttatccgc tcacaattcc acacaacata

11401 cgagccggaa gcataaagtg taaagcctgg ggtgcctaat gagtgagcta actcacatta

11461 attgcgttgc gctcactgcc cgctttccag tcgggaaacc tgtcgtgcca gctgcattaa

11521 tgaatcggcc aacgcgcggg gagaggcggt ttgcgtattg ggcgctcttc cgcttcctcg

11581 ctcactgact cgctgcgctc ggtcgttcgg ctgcggcgag cggtatcagc tcactcaaag

11641 gcggtaatac ggttatccac agaatcaggg gataacgcag gaaagaacat gtgagcaaaa

11701 ggccagcaaa aggccaggaa ccgtaaaaag gccgcgttgc tggcgttttt ccataggctc

11761 cgcccccctg acgagcatca caaaaatcga cgctcaagtc agaggtggcg aaacccgaca

11821 ggactataaa gataccaggc gtttccccct ggaagctccc tcgtgcgctc tcctgttccg

11881 accctgccgc ttaccggata cctgtccgcc tttctccctt cgggaagcgt ggcgctttct

11941 catagctcac gctgtaggta tctcagttcg gtgtaggtcg ttcgctccaa gctgggctgt

12001 gtgcacgaac cccccgttca gcccgaccgc tgcgccttat ccggtaacta tcgtcttgag

12061 tccaacccgg taagacacga cttatcgcca ctggcagcag ccactggtaa caggattagc

12121 agagcgaggt atgtaggcgg tgctacagag ttcttgaagt ggtggcctaa ctacggctac

12181 actagaagga cagtatttgg tatctgcgct ctgctgaagc cagttacctt cggaaaaaga

12241 gttggtagct cttgatccgg caaacaaacc accgctggta gcggtggttt ttttgtttgc

12301 aagcagcaga ttacgcgcag aaaaaaagga tctcaagaag atcctttgat cttttctacg

12361 gggtctgacg ctcagtggaa cgaaaactca cgttaaggga ttttggtcat gagattatca

12421 aaaaggatct tcacctagat ccttttaaat taaaaatgaa gttttaaatc aatctaaagt

12481 atatatgagt aaacttggtc tgacagttac caatgcttaa tcagtgaggc acctatctca

12541 gcgatctgtc tatttcgttc atccatagtt gcctgactcc ccgtcgtgta gataactacg

12601 atacgggagg gcttaccatc tggccccagt gctgcaatga taccgcgaga cccacgctca

12661 ccggctccag atttatcagc aataaaccag ccagccggaa gggccgagcg cagaagtggt

12721 cctgcaactt tatccgcctc catccagtct attaattgtt gccgggaagc tagagtaagt

12781 agttcgccag ttaatagttt gcgcaacgtt gttgccattg ctacaggcat cgtggtgtca

12841 cgctcgtcgt ttggtatggc ttcattcagc tccggttccc aacgatcaag gcgagttaca

12901 tgatccccca tgttgtgcaa aaaagcggtt agctccttcg gtcctccgat cgttgtcaga

12961 agtaagttgg ccgcagtgtt atcactcatg gttatggcag cactgcataa ttctcttact

13021 gtcatgccat ccgtaagatg cttttctgtg actggtgagt actcaaccaa gtcattctga

13081 gaatagtgta tgcggcgacc gagttgctct tgcccggcgt caatacggga taataccgcg

13141 ccacatagca gaactttaaa agtgctcatc attggaaaac gttcttcggg gcgaaaactc

13201 tcaaggatct taccgctgtt gagatccagt tcgatgtaac ccactcgtgc acccaactga

13261 tcttcagcat cttttacttt caccagcgtt tctgggtgag caaaaacagg aaggcaaaat

13321 gccgcaaaaa agggaataag ggcgacacgg aaatgttgaa tactcatact cttccttttt

13381 caatattatt gaagcattta tcagggttat tgtctcatga gcggatacat atttgaatgt

13441 atttagaaaa ataaacaaat aggggttccg cgcacatttc cccgaaaagt gc

//

LOCUS	pVSVdG-mCherry-Linker	13477 bp DNA	circular SYN 11-DEC-2023

DEFINITION synthetic circular DNA

ACCESSION	.

VERSION	.

KEYWORDS	pVSVdG-mCherry-linker

SOURCE	synthetic DNA construct

ORGANISM synthetic DNA construct

REFERENCE	1 (bases 1 to 13477)

AUTHORS	.

TITLE	Direct Submission

JOURNAL	Exported Dec 11, 2023 from SnapGene Viewer 7.0.3

https://www.snapgene.com

FEATURES	Location/Qualifiers

source	1..13477

/mol_type="other DNA"

/organism="synthetic DNA construct"

rep_origin	complement(33..461)

/direction=LEFT

/label=f1 ori

/note="f1 bacteriophage origin of replication; arrow

indicates direction of (+) strand synthesis"

CDS	complement(463..531)

/label=LacZ alpha

primer_bind	603..619

/label=M13 fwd

/note="common sequencing primer, one of multiple similar

variants"

promoter	626..644

/label=T7 promoter

/note="promoter for bacteriophage T7 RNA polymerase"

CDS	708..1976

/codon_start=1

/label=N

/translation="MSVTVKRIIDNTVVVPKLPANEDPVEYPADYFRKSKEIPLYINTT

KSLSDLRGYVYQGLKSGNVSIIHVNSYLYGALKDIRGKLDKDWSSFGINIGKAGDTIGI

FDLVSLKALDGVLPDGVSDASRTSADDKWLPLYLLGLYRVGRTQMPEYRKKLMDGLTNQ

CKMINEQFEPLVPEGRDIFDVWGNDSNYTKIVAAVDMFFHMFKKHECASFRYGTIVSRF

KDCAALATFGHLCKITGMSTEDVTTWILNREVADEMVQMMLPGQEIDKADSYMPYLIDF

GLSSKSPYSSVKNPAFHFWGQLTALLLRSTRARNARQPDDIEYTSLTTAGLLYAYAVGS

SADLAQQFCVGDNKYTPDDSTGGLTTNAPPQGRDVVEWLGWFEDQNRKPTPDMMQYAKR

AVMSLQGLREKTIGKYAKSEFDK"

CDS	2040..2837

/codon_start=1

/label=P

/translation="MDNLTKVREYLKSYSRLDQAVGEIDEIEAQRAEKSNYELFQEDGV

EEHTKPSYFQAADDSDTESEPEIEDNQGLYAPDPEAEQVEGFIQGPLDDYADEEVDVVF

TSDWKQPELESDEHGKTLRLTSPEGLSGEQKSQWLSTIKAVVQSAKYWNLAECTFEASG

EGVIMKERQITPDVYKVTPVMNTHPSQSEAVSDVWSLSKTSMTFQPKKASLQPLTISLD

ELFSSRGEFISVGGDGRMSHKEAILLGLRYKKLYNQARVKYSL"

CDS	2894..3583

/codon_start=1

/label=M

/translation="MSSLKKILGLKGKGKKSKKLGIAPPPYEEDTSMEYAPSAPIDKSY

FGVDEMDTYDPNQLRYEKFFFTVKMTVRSNRPFRTYSDVAAAVSHWDHMYIGMAGKRPF

YKILAFLGSSNLKATPAVLADQGQPEYHAHCEGRAYLPHRMGKTPPMLNVPEHFRRPFN

IGLYKGTIELTMTIYDDESLEAAPMIWDHFNSSKFSDFREKALMFGLIVEKKASGAWVL

DSIGHFK"

misc_feature	3712..3729

/label=stuffer

misc_feature	3777..4486

/direction=RIGHT

/note="note=""Fragment extracted from

VSV_GFP_with_genes.xdna [725 nt] : (XhoI[4656] /

NheI[5385])"

CDS	3781..4485

/label=mCherry

misc_feature	4595..10924

/label=L Protein

CDS	4595..10924

/codon_start=1

/label=L

/translation="MEVHDFETDEFNDFNEDDYATREFLNPDERMTYLNHADYNLNSPL

ISDDIDNLIRKFNSLPIPSMWDSKNWDGVLEMLTSCQANPIPTSQMHKWMGSWLMSDNH

DASQGYSFLHEVDKEAEITFDVVETFIRGWGNKPIEYIKKERWTDSFKILAYLCQKFLD

LHKLTLILNAVSEVELLNLARTFKGKVRRSSHGTNICRIRVPSLGPTFISEGWAYFKKL

DILMDRNFLLMVKDVIIGRMQTVLSMVCRIDNLFSEQDIFSLLNIYRIGDKIVERQGNF

SYDLIKMVEPICNLKLMKLARESRPLVPQFPHFENHIKTSVDEGAKIDRGIRFLHDQIM

SVKTVDLTLVIYGSFRHWGHPFIDYYTGLEKLHSQVTMKKDIDVSYAKALASDLARIVL

FQQFNDHKKWFVNGDLLPHDHPFKSHVKENTWPTAAQVQDFGDKWHELPLIKCFEIPDL

LDPSIIYSDKSHSMNRSEVLKHVRMNPNTPIPSKKVLQTMLDTKATNWKEFLKEIDEKG

LDDDDLIIGLKGKERELKLAGRFFSLMSWKLREYFVITEYLIKTHFVPMFKGLTMADDL

TAVIKKMLDSSSGQGLKSYEAICIANHIDYEKWNNHQRKLSNGPVFRVMGQFLGYPSLI

ERTHEFFEKSLIYYNGRPDLMRVHNNTLINSTSQRVCWQGQEGGLEGLRQKGWSILNLL

VIQREAKIRNTAVKVLAQGDNQVICTQYKTKKSRNVVELQGALNQMVSNNEKIMTAIKI

GTGKLGLLINDDETMQSADYLNYGKIPIFRGVIRGLETKRWSRVTCVTNDQIPTCANIM

SSVSTNALTVAHFAENPINAMIQYNYFGTFARLLLMMHDPALRQSLYEVQDKIPGLHSS

TFKYAMLYLDPSIGGVSGMSLSRFLIRAFPDPVTESLSFWRFIHVHARSEHLKEMSAVF

GNPEIAKFRITHIDKLVEDPTSLNIAMGMSPANLLKTEVKKCLIESRQTIRNQVIKDAT

IYLYHEEDRLRSFLWSINPLFPRFLSEFKSGTFLGVADGLISLFQNSRTIRNSFKKKYH

RELDDLIVRSEVSSLTHLGKLHLRRGSCKMWTCSATHADTLRYKSWGRTVIGTTVPHPL

EMLGPQHRKETPCAPCNTSGFNYVSVHCPDGIHDVFSSRGPLPAYLGSKTSESTSILQP

WERESKVPLIKRATRLRDAISWFVEPDSKLAMTILSNIHSLTGEEWTKRQHGFKRTGSA

LHRFSTSRMSHGGFASQSTAALTRLMATTDTMRDLGDQNFDFLFQATLLYAQITTTVAR

DGWITSCTDHYHIACKSCLRPIEEITLDSSMDYTPPDVSHVLKTWRNGEGSWGQEIKQI

YPLEGNWKNLAPAEQSYQVGRCIGFLYGDLAYRKSTHAEDSSLFPLSIQGRIRGRGFLK

GLLDGLMRASCCQVIHRRSLAHLKRPANAVYGGLIYLIDKLSVSPPFLSLTRSGPIRDE

LETIPHKIPTSYPTSNRDMGVIVRNYFKYQCRLIEKGKYRSHYSQLWLFSDVLSIDFIG

PFSISTTLLQILYKPFLSGKDKNELRELANLSSLLRSGEGWEDIHVKFFTKDILLCPEE

IRHACKFGIAKDNNKDMSYPPWGRESRGTITTIPVYYTTTPYPKMLEMPPRIQNPLLSG

IRLGQLPTGAHYKIRSILHGMGIHYRDFLSCGDGSGGMTAALLRENVHSRGIFNSLLEL

SGSVMRGASPEPPSALETLGGDKSRCVNGETCWEYPSDLCDPRTWDYFLRLKAGLGLQI

DLIVMDMEVRDSSTSLKIETNVRNYVHRILDEQGVLIYKTYGTYICESEKNAVTILGPM

FKTVDLVQTEFSSSQTSEVYMVCKGLKKLIDEPNPDWSSINESWKNLYAFQSSEQEFAR

AKKVSTYFTLTGIPSQFIPDPFVNIETMLQIFGVPTGVSHAAALKSSDRPADLLTISLF

YMAIISYYNINHIRVGPIPPNPPSDGIAQNVGIAITGISFWLSLMEKDIPLYQQCLAVI

QQSFPIRWEAVSVKGGYKQKWSTRGDGLPKDTRISDSLAPIGNWIRSLELVRNQVRLNP

FNEILFNQLCRTVDNHLKWSNLRRNTGMIEWINRRISKEDRSILMLKSDLHEENSWRD"

terminator	11177..11224

/label=T7 terminator

/note="transcription terminator for bacteriophage T7 RNA

polymerase"

primer_bind	complement(11290..11309)

/label=T3

promoter	complement(11292..11310)

/label=T3 promoter

/note="promoter for bacteriophage T3 RNA polymerase"

primer_bind	complement(11327..11347)

/label=M13-rev

primer_bind	complement(11331..11347)

/label=M13 rev

/note="common sequencing primer, one of multiple similar

variants"

misc_binding	complement(11353..11375)

/label=LacO

protein_bind	11355..11371

/label=lac operator

/bound_moiety="lac repressor encoded by lacI"

/note="The lac repressor binds to the lac operator to

inhibit transcription in E. coli. This inhibition can be

relieved by adding lactose or

isopropyl-beta-D-thiogalactopyranoside (IPTG)."

promoter	complement(11379..11409)

/label=lac promoter

/note="promoter for the E. coli lac operon"

promoter	complement(11380..11409)

/label=lac

protein_bind	11424..11445

/label=CAP binding site

/bound_moiety="E. coli catabolite activator protein"

/note="CAP binding activates transcription in the presence

of cAMP."

rep_origin	complement(11715..12343)

/direction=LEFT

/label=ColE1 origin

rep_origin	complement(11733..12321)

/direction=LEFT

/label=ori

/note="high-copy-number ColE1/pMB1/pBR322/pUC origin of

replication"

CDS	complement(12492..13352)

/codon_start=1

/gene="bla"

/product="beta-lactamase"

/label=AmpR

/note="confers resistance to ampicillin, carbenicillin, and

related antibiotics"

/translation="MSIQHFRVALIPFFAAFCLPVFAHPETLVKVKDAEDQLGARVGYI

ELDLNSGKILESFRPEERFPMMSTFKVLLCGAVLSRIDAGQEQLGRRIHYSQNDLVEYS

PVTEKHLTDGMTVRELCSAAITMSDNTAANLLLTTIGGPKELTAFLHNMGDHVTRLDRW

EPELNEAIPNDERDTTMPVAMATTLRKLLTGELLTLASRQQLIDWMEADKVAGPLLRSA

LPAGWFIADKSGAGERGSRGIIAALGPDGKPSRIVVIYTTGSQATMDERNRQIAEIGAS

LIKHW"

promoter	complement(13353..13457)

/gene="bla"

/label=AmpR promoter

ORIGIN

1 cacctaaatt gtaagcgtta atattttgtt aaaattcgcg ttaaattttt gttaaatcag

61 ctcatttttt aaccaatagg ccgaaatcgg caaaatccct tataaatcaa aagaatagac

121 cgagataggg ttgagtgttg ttccagtttg gaacaagagt ccactattaa agaacgtgga

181 ctccaacgtc aaagggcgaa aaaccgtcta tcagggcgat ggcccactac gtgaaccatc

241 accctaatca agttttttgg ggtcgaggtg ccgtaaagca ctaaatcgga accctaaagg

301 gagcccccga tttagagctt gacggggaaa gccggcgaac gtggcgagaa aggaagggaa

361 gaaagcgaaa ggagcgggcg ctagggcgct ggcaagtgta gcggtcacgc tgcgcgtaac

421 caccacaccc gccgcgctta atgcgccgct acagggcgcg tcccattcgc cattcaggct

481 gcgcaactgt tgggaagggc gatcggtgcg ggcctcttcg ctattacgcc agctggcgaa

541 agggggatgt gctgcaaggc gattaagttg ggtaacgcca gggttttccc agtcacgacg

601 ttgtaaaacg acggccagtg aattgtaata cgactcacta taggacgaag acaaacaaac

661 cattattatc attaaaaggc tcaggagaaa ctttaacagt aatcaaaatg tctgttacag

721 tcaagagaat cattgacaac acagtcgtag ttccaaaact tcctgcaaat gaggatccag

781 tggaataccc ggcagattac ttcagaaaat caaaggagat tcctctttac atcaatacta

841 caaaaagttt gtcagatcta agaggatatg tctaccaagg cctcaaatcc ggaaatgtat

901 caatcataca tgtcaacagc tacttgtatg gagcattaaa ggacatccgg ggtaagttgg

961 ataaagattg gtcaagtttc ggaataaaca tcgggaaagc aggggataca atcggaatat

1021 ttgaccttgt atccttgaaa gccctggacg gcgtacttcc agatggagta tcggatgctt

1081 ccagaaccag cgcagatgac aaatggttgc ctttgtatct acttggctta tacagagtgg

1141 gcagaacaca aatgcctgaa tacagaaaaa agctcatgga tgggctgaca aatcaatgca

1201 aaatgatcaa tgaacagttt gaacctcttg tgccagaagg tcgtgacatt tttgatgtgt

1261 ggggaaatga cagtaattac acaaaaattg tcgctgcagt ggacatgttc ttccacatgt

1321 tcaaaaaaca tgaatgtgcc tcgttcagat acggaactat tgtttccaga ttcaaagatt

1381 gtgctgcatt ggcaacattt ggacacctct gcaaaataac cggaatgtct acagaagatg

1441 taacgacctg gatcttgaac cgagaagttg cagatgaaat ggtccaaatg atgcttccag

1501 gccaagaaat tgacaaggcc gattcataca tgccttattt gatcgacttt ggattgtctt

1561 ctaagtctcc atattcttcc gtcaaaaacc ctgccttcca cttctggggg caattgacag

1621 ctcttctgct cagatccacc agagcaagga atgcccgaca gcctgatgac attgagtata

1681 catctcttac tacagcaggt ttgttgtacg cttatgcagt aggatcctct gccgacttgg

1741 cacaacagtt ttgtgttgga gataacaaat acactccaga tgatagtacc ggaggattga

1801 cgactaatgc accgccacaa ggcagagatg tggtcgaatg gctcggatgg tttgaagatc

1861 aaaacagaaa accgactcct gatatgatgc agtatgcgaa aagagcagtc atgtcactgc

1921 aaggcctaag agagaagaca attggcaagt atgctaagtc agaatttgac aaatgaccct

1981 ataattctca gatcacctat tatatattat gctacatatg aaaaaaacta acagatatca

2041 tggataatct cacaaaagtt cgtgagtatc tcaagtccta ttctcgtctg gatcaggcgg

2101 taggagagat agatgagatc gaagcacaac gagctgaaaa gtccaattat gagttgttcc

2161 aagaggatgg agtggaagag catactaagc cctcttattt tcaggcagca gatgattctg

2221 acacagaatc tgaaccagaa attgaagaca atcaaggctt gtatgcacca gatccagaag

2281 ctgagcaagt tgaaggcttt atacaggggc ctttagatga ctatgcagat gaggaagtgg

2341 atgttgtatt tacttcggac tggaaacagc ctgagcttga atctgacgag catggaaaga

2401 ccttacggtt gacatcgcca gagggtttaa gtggagagca gaaatcccag tggctttcga

2461 cgattaaagc agtcgtgcaa agtgccaaat actggaatct ggcagagtgc acatttgaag

2521 catcgggaga aggggtcatt atgaaggagc gccagataac tccggatgta tataaggtca

2581 ctccagtgat gaacacacat ccgtcccaat cagaagcagt atcagatgtt tggtctctct

2641 caaagacatc catgactttc caacccaaga aagcaagtct tcagcctctc accatatcct

2701 tggatgaatt gttctcatct agaggagagt tcatctctgt cggaggtgac ggacgaatgt

2761 ctcataaaga ggccatcctg ctcggcctga gatacaaaaa gttgtacaat caggcgagag

2821 tcaaatattc tctgtagact atgaaaaaaa gtaacagata tcacgatcta agtgttatcc

2881 caatccattc atcatgagtt ccttaaagaa gattctcggt ctgaagggga aaggtaagaa

2941 atctaagaaa ttagggatcg caccaccccc ttatgaagag gacactagca tggagtatgc

3001 tccgagcgct ccaattgaca aatcctattt tggagttgac gagatggaca cctatgatcc

3061 gaatcaatta agatatgaga aattcttctt tacagtgaaa atgacggtta gatctaatcg

3121 tccgttcaga acatactcag atgtggcagc cgctgtatcc cattgggatc acatgtacat

3181 cggaatggca gggaaacgtc ccttctacaa aatcttggct tttttgggtt cttctaatct

3241 aaaggccact ccagcggtat tggcagatca aggtcaacca gagtatcacg ctcactgcga

3301 aggcagggct tatttgccac ataggatggg gaagacccct cccatgctca atgtaccaga

3361 gcacttcaga agaccattca atataggtct ttacaaggga acgattgagc tcacaatgac

3421 catctacgat gatgagtcac tggaagcagc tcctatgatc tgggatcatt tcaattcttc

3481 caaattttct gatttcagag agaaggcctt aatgtttggc ctgattgtcg agaaaaaggc

3541 atctggagcg tgggtcctgg actctatcgg ccacttcaaa tgagctagtc taacttctag

3601 cttctgaaca atccccggtt tactcagtct cccctaattc cagcctctcg aacaactaat

3661 atcctgtctt ttctatccct atgaaaaaaa ctaacagaga tcgatctgtt tacgcgtcac

3721 tttaattaac tcaaatcctg ctaggtatga aaaaaactaa cagatatcac gctcgagacc

3781 gtgagcaagg gcgaggagga taacatggcc atcatcaagg agttcatgcg cttcaaggtg

3841 cacatggagg gctccgtgaa cggccacgag ttcgagatcg agggcgaggg cgagggccgc

3901 ccctacgagg gcacccagac cgccaagctg aaggtgacca agggtggccc cctgcccttc

3961 gcctgggaca tcctgtcccc tcagttcatg tacggctcca aggcctacgt gaagcacccc

4021 gccgacatcc ccgactactt gaagctgtcc ttccccgagg gcttcaagtg ggagcgcgtg

4081 atgaacttcg aggacggcgg cgtggtgacc gtgacccagg actcctccct gcaggacggc

4141 gagttcatct acaaggtgaa gctgcgcggc accaacttcc cctccgacgg ccccgtaatg

4201 cagaagaaga ccatgggctg ggaggcctcc tccgagcgga tgtaccccga ggacggcgcc

4261 ctgaagggcg agatcaagca gaggctgaag ctgaaggacg gcggccacta cgacgctgag

4321 gtcaagacca cctacaaggc caagaagccc gtgcagctgc ccggcgccta caacgtcaac

4381 atcaagttgg acatcacctc ccacaacgag gactacacca tcgtggaaca gtacgaacgc

4441 gccgagggcc gccactccac cggcggcatg gacgagctgt acaaggctag ccagattctt

4501 catgtttgga ccaaatcaac ttgtgatacc atgctcaaag aggcctcaat tatatttgag

4561 tttttaattt ttatgaaaaa aactaacagc aatcatggaa gtccacgatt ttgagaccga

4621 cgagttcaat gatttcaatg aagatgacta tgccacaaga gaattcctga atcccgatga

4681 gcgcatgacg tacttgaatc atgctgatta caacctgaat tctcctctaa ttagtgatga

4741 tattgacaat ttaatcagga aattcaattc tcttccaatt ccctcgatgt gggatagtaa

4801 gaactgggat ggagttcttg agatgttaac gtcatgtcaa gccaatccca tcccaacatc

4861 tcagatgcat aaatggatgg gaagttggtt aatgtctgat aatcatgatg ccagtcaagg

4921 gtatagtttt ttacatgaag tggacaaaga ggcagaaata acatttgacg tggtggagac

4981 cttcatccgc ggctggggca acaaaccaat tgaatacatc aaaaaggaaa gatggactga

5041 ctcattcaaa attctcgctt atttgtgtca aaagtttttg gacttacaca agttgacatt

5101 aatcttaaat gctgtctctg aggtggaatt gctcaacttg gcgaggactt tcaaaggcaa

5161 agtcagaaga agttctcatg gaacgaacat atgcaggatt agggttccca gcttgggtcc

5221 tacttttatt tcagaaggat gggcttactt caagaaactt gatattctaa tggaccgaaa

5281 ctttctgtta atggtcaaag atgtgattat agggaggatg caaacggtgc tatccatggt

5341 atgtagaata gacaacctgt tctcagagca agacatcttc tcccttctaa atatctacag

5401 aattggagat aaaattgtgg agaggcaggg aaatttttct tatgacttga ttaaaatggt

5461 ggaaccgata tgcaacttga agctgatgaa attagcaaga gaatcaaggc ctttagtccc

5521 acaattccct cattttgaaa atcatatcaa gacttctgtt gatgaagggg caaaaattga

5581 ccgaggtata agattcctcc atgatcagat aatgagtgtg aaaacagtgg atctcacact

5641 ggtgatttat ggatcgttca gacattgggg tcatcctttt atagattatt acactggact

5701 agaaaaatta cattcccaag taaccatgaa gaaagatatt gatgtgtcat atgcaaaagc

5761 acttgcaagt gatttagctc ggattgttct atttcaacag ttcaatgatc ataaaaagtg

5821 gttcgtgaat ggagacttgc tccctcatga tcatcccttt aaaagtcatg ttaaagaaaa

5881 tacatggccc acagctgctc aagttcaaga ttttggagat aaatggcatg aacttccgct

5941 gattaaatgt tttgaaatac ccgacttact agacccatcg ataatatact ctgacaaaag

6001 tcattcaatg aataggtcag aggtgttgaa acatgtccga atgaatccga acactcctat

6061 ccctagtaaa aaggtgttgc agactatgtt ggacacaaag gctaccaatt ggaaagaatt

6121 tcttaaagag attgatgaga agggcttaga tgatgatgat ctaattattg gtcttaaagg

6181 aaaggagagg gaactgaagt tggcaggtag atttttctcc ctaatgtctt ggaaattgcg

6241 agaatacttt gtaattaccg aatatttgat aaagactcat ttcgtcccta tgtttaaagg

6301 cctgacaatg gcggacgatc taactgcagt cattaaaaag atgttagatt cctcatccgg

6361 ccaaggattg aagtcatatg aggcaatttg catagccaat cacattgatt acgaaaaatg

6421 gaataaccac caaaggaagt tatcaaacgg cccagtgttc cgagttatgg gccagttctt

6481 aggttatcca tccttaatcg agagaactca tgaatttttt gagaaaagtc ttatatacta

6541 caatggaaga ccagacttga tgcgtgttca caacaacaca ctgatcaatt caacctccca

6601 acgagtttgt tggcaaggac aagagggtgg actggaaggt ctacggcaaa aaggatggag

6661 tatcctcaat ctactggtta ttcaaagaga ggctaaaatc agaaacactg ctgtcaaagt

6721 cttggcacaa ggtgataatc aagttatttg cacacagtat aaaacgaaga aatcgagaaa

6781 cgttgtagaa ttacagggtg ctctcaatca aatggtttct aataatgaga aaattatgac

6841 tgcaatcaaa atagggacag ggaagttagg acttttgata aatgacgatg agactatgca

6901 atctgcagat tacttgaatt atggaaaaat accgattttc cgtggagtga ttagagggtt

6961 agagaccaag agatggtcac gagtgacttg tgtcaccaat gaccaaatac ccacttgtgc

7021 taatataatg agctcagttt ccacaaatgc tctcaccgta gctcattttg ctgagaaccc

7081 aatcaatgcc atgatacagt acaattattt tgggacattt gctagactct tgttgatgat

7141 gcatgatcct gctcttcgtc aatcattgta tgaagttcaa gataagatac cgggcttgca

7201 cagttctact ttcaaatacg ccatgttgta tttggaccct tccattggag gagtgtcggg

7261 catgtctttg tccaggtttt tgattagagc cttcccagat cccgtaacag aaagtctctc

7321 attctggaga ttcatccatg tacatgctcg aagtgagcat ctgaaggaga tgagtgcagt

7381 atttggaaac cccgagatag ccaagtttcg aataactcac atagacaagc tagtagaaga

7441 tccaacctct ctgaacatcg ctatgggaat gagtccagcg aacttgttaa agactgaggt

7501 taaaaaatgc ttaatcgaat caagacaaac catcaggaac caggtgatta aggatgcaac

7561 catatatttg tatcatgaag aggatcggct cagaagtttc ttatggtcaa taaatcctct

7621 gttccctaga tttttaagtg aattcaaatc aggcactttt ttgggagtcg cagacgggct

7681 catcagtcta tttcaaaatt ctcgtactat tcggaactcc tttaagaaaa agtatcatag

7741 ggaattggat gatttgattg tgaggagtga ggtatcctct ttgacacatt tagggaaact

7801 tcatttgaga aggggatcat gtaaaatgtg gacatgttca gctactcatg ctgacacatt

7861 aagatacaaa tcctggggcc gtacagttat tgggacaact gtaccccatc cattagaaat

7921 gttgggtcca caacatcgaa aagagactcc ttgtgcacca tgtaacacat cagggttcaa

7981 ttatgtttct gtgcattgtc cagacgggat ccatgacgtc tttagttcac ggggaccatt

8041 gcctgcttat ctagggtcta aaacatctga atctacatct attttgcagc cttgggaaag

8101 ggaaagcaaa gtcccactga ttaaaagagc tacacgtctt agagatgcta tctcttggtt

8161 tgttgaaccc gactctaaac tagcaatgac tatactttct aacatccact ctttaacagg

8221 cgaagaatgg accaaaaggc agcatgggtt caaaagaaca gggtctgccc ttcataggtt

8281 ttcgacatct cggatgagcc atggtgggtt cgcatctcag agcactgcag cattgaccag

8341 gttgatggca actacagaca ccatgaggga tctgggagat cagaatttcg actttttatt

8401 ccaagcaacg ttgctctatg ctcaaattac caccactgtt gcaagagacg gatggatcac

8461 cagttgtaca gatcattatc atattgcctg taagtcctgt ttgagaccca tagaagagat

8521 caccctggac tcaagtatgg actacacgcc cccagatgta tcccatgtgc tgaagacatg

8581 gaggaatggg gaaggttcgt ggggacaaga gataaaacag atctatcctt tagaagggaa

8641 ttggaagaat ttagcacctg ctgagcaatc ctatcaagtc ggcagatgta taggttttct

8701 atatggagac ttggcgtata gaaaatctac tcatgccgag gacagttctc tatttcctct

8761 atctatacaa ggtcgtatta gaggtcgagg tttcttaaaa gggttgctag acggattaat

8821 gagagcaagt tgctgccaag taatacaccg gagaagtctg gctcatttga agaggccggc

8881 caacgcagtg tacggaggtt tgatttactt gattgataaa ttgagtgtat cacctccatt

8941 cctttctctt actagatcag gacctattag agacgaatta gaaacgattc cccacaagat

9001 cccaacctcc tatccgacaa gcaaccgtga tatgggggtg attgtcagaa attacttcaa

9061 ataccaatgc cgtctaattg aaaagggaaa atacagatca cattattcac aattatggtt

9121 attctcagat gtcttatcca tagacttcat tggaccattc tctatttcca ccaccctctt

9181 gcaaatccta tacaagccat ttttatctgg gaaagataag aatgagttga gagagctggc

9241 aaatctttct tcattgctaa gatcaggaga ggggtgggaa gacatacatg tgaaattctt

9301 caccaaggac atattattgt gtccagagga aatcagacat gcttgcaagt tcgggattgc

9361 taaggataat aataaagaca tgagctatcc cccttgggga agggaatcca gagggacaat

9421 tacaacaatc cctgtttatt atacgaccac cccttaccca aagatgctag agatgcctcc

9481 aagaatccaa aatcccctgc tgtccggaat caggttgggc caattaccaa ctggcgctca

9541 ttataaaatt cggagtatat tacatggaat gggaatccat tacagggact tcttgagttg

9601 tggagacggc tccggaggga tgactgctgc attactacga gaaaatgtgc atagcagagg

9661 aatattcaat agtctgttag aattatcagg gtcagtcatg cgaggcgcct ctcctgagcc

9721 ccccagtgcc ctagaaactt taggaggaga taaatcgaga tgtgtaaatg gtgaaacatg

9781 ttgggaatat ccatctgact tatgtgaccc aaggacttgg gactatttcc tccgactcaa

9841 agcaggcttg gggcttcaaa ttgatttaat tgtaatggat atggaagttc gggattcttc

9901 tactagcctg aaaattgaga cgaatgttag aaattatgtg caccggattt tggatgagca

9961 aggagtttta atctacaaga cttatggaac atatatttgt gagagcgaaa agaatgcagt

10021 aacaatcctt ggtcccatgt tcaagacggt cgacttagtt caaacagaat ttagtagttc

10081 tcaaacgtct gaagtatata tggtatgtaa aggtttgaag aaattaatcg atgaacccaa

10141 tcccgattgg tcttccatca atgaatcctg gaaaaacctg tacgcattcc agtcatcaga

10201 acaggaattt gccagagcaa agaaggttag tacatacttt accttgacag gtattccctc

10261 ccaattcatt cctgatcctt ttgtaaacat tgagactatg ctacaaatat tcggagtacc

10321 cacgggtgtg tctcatgcgg ctgccttaaa atcatctgat agacctgcag atttattgac

10381 cattagcctt ttttatatgg cgattatatc gtattataac atcaatcata tcagagtagg

10441 accgatacct ccgaaccccc catcagatgg aattgcacaa aatgtgggga tcgctataac

10501 tggtataagc ttttggctga gtttgatgga gaaagacatt ccactatatc aacagtgttt

10561 agcagttatc cagcaatcat tcccgattag gtgggaggct gtttcagtaa aaggaggata

10621 caagcagaag tggagtacta gaggtgatgg gctcccaaaa gatacccgaa tttcagactc

10681 cttggcccca atcgggaact ggatcagatc tctggaattg gtccgaaacc aagttcgtct

10741 aaatccattc aatgagatct tgttcaatca gctatgtcgt acagtggata atcatttgaa

10801 atggtcaaat ttgcgaagaa acacaggaat gattgaatgg atcaatagac gaatttcaaa

10861 agaagaccgg tctatactga tgttgaagag tgacctacac gaggaaaact cttggagaga

10921 ttaaaaaatc atgaggagac tccaaacttt aagtatgaaa aaaactttga tccttaagac

10981 cctcttgtgg tttttatttt ttatctggtt ttgtggtctt cgtgggtcgg catggcatct

11041 ccacctcctc gcggtccgac ctgggcatcc gaaggaggac gtcgtccact cggatggcta

11101 agggaggggc ccccgcgggg ctgctaacaa agcccgaaag gaagctgagt tggctgctgc

11161 caccgctgag caataactag cataacccct tggggcctct aaacgggtct tgaggggttt

11221 tttgctgaaa ggaggaacta tatccggatc gagacctcga tactagtgcg gtggagctcc

11281 agcttttgtt ccctttagtg agggttaatt tcgagcttgg cgtaatcatg gtcatagctg

11341 tttcctgtgt gaaattgtta tccgctcaca attccacaca acatacgagc cggaagcata

11401 aagtgtaaag cctggggtgc ctaatgagtg agctaactca cattaattgc gttgcgctca

11461 ctgcccgctt tccagtcggg aaacctgtcg tgccagctgc attaatgaat cggccaacgc

11521 gcggggagag gcggtttgcg tattgggcgc tcttccgctt cctcgctcac tgactcgctg

11581 cgctcggtcg ttcggctgcg gcgagcggta tcagctcact caaaggcggt aatacggtta

11641 tccacagaat caggggataa cgcaggaaag aacatgtgag caaaaggcca gcaaaaggcc

11701 aggaaccgta aaaaggccgc gttgctggcg tttttccata ggctccgccc ccctgacgag

11761 catcacaaaa atcgacgctc aagtcagagg tggcgaaacc cgacaggact ataaagatac

11821 caggcgtttc cccctggaag ctccctcgtg cgctctcctg ttccgaccct gccgcttacc

11881 ggatacctgt ccgcctttct cccttcggga agcgtggcgc tttctcatag ctcacgctgt

11941 aggtatctca gttcggtgta ggtcgttcgc tccaagctgg gctgtgtgca cgaacccccc

12001 gttcagcccg accgctgcgc cttatccggt aactatcgtc ttgagtccaa cccggtaaga

12061 cacgacttat cgccactggc agcagccact ggtaacagga ttagcagagc gaggtatgta

12121 ggcggtgcta cagagttctt gaagtggtgg cctaactacg gctacactag aaggacagta

12181 tttggtatct gcgctctgct gaagccagtt accttcggaa aaagagttgg tagctcttga

12241 tccggcaaac aaaccaccgc tggtagcggt ggtttttttg tttgcaagca gcagattacg

12301 cgcagaaaaa aaggatctca agaagatcct ttgatctttt ctacggggtc tgacgctcag

12361 tggaacgaaa actcacgtta agggattttg gtcatgagat tatcaaaaag gatcttcacc

12421 tagatccttt taaattaaaa atgaagtttt aaatcaatct aaagtatata tgagtaaact

12481 tggtctgaca gttaccaatg cttaatcagt gaggcaccta tctcagcgat ctgtctattt

12541 cgttcatcca tagttgcctg actccccgtc gtgtagataa ctacgatacg ggagggctta

12601 ccatctggcc ccagtgctgc aatgataccg cgagacccac gctcaccggc tccagattta

12661 tcagcaataa accagccagc cggaagggcc gagcgcagaa gtggtcctgc aactttatcc

12721 gcctccatcc agtctattaa ttgttgccgg gaagctagag taagtagttc gccagttaat

12781 agtttgcgca acgttgttgc cattgctaca ggcatcgtgg tgtcacgctc gtcgtttggt

12841 atggcttcat tcagctccgg ttcccaacga tcaaggcgag ttacatgatc ccccatgttg

12901 tgcaaaaaag cggttagctc cttcggtcct ccgatcgttg tcagaagtaa gttggccgca

12961 gtgttatcac tcatggttat ggcagcactg cataattctc ttactgtcat gccatccgta

13021 agatgctttt ctgtgactgg tgagtactca accaagtcat tctgagaata gtgtatgcgg

13081 cgaccgagtt gctcttgccc ggcgtcaata cgggataata ccgcgccaca tagcagaact

13141 ttaaaagtgc tcatcattgg aaaacgttct tcggggcgaa aactctcaag gatcttaccg

13201 ctgttgagat ccagttcgat gtaacccact cgtgcaccca actgatcttc agcatctttt

13261 actttcacca gcgtttctgg gtgagcaaaa acaggaaggc aaaatgccgc aaaaaaggga

13321 ataagggcga cacggaaatg ttgaatactc atactcttcc tttttcaata ttattgaagc

13381 atttatcagg gttattgtct catgagcgga tacatatttg aatgtattta gaaaaataaa

13441 caaatagggg ttccgcgcac atttccccga aaagtgc

//

LOCUS	pVSV-mCherry-rose	15011 bp DNA	circular SYN 22-NOV-2023

DEFINITION mCherry cloned into pVSV_XN2 from John Rose.

ACCESSION	.

VERSION	.

KEYWORDS	pVSV-mCherry

SOURCE	synthetic DNA construct

ORGANISM synthetic DNA construct

REFERENCE	1 (bases 1 to 15011)

AUTHORS	.

TITLE	Direct Submission

JOURNAL	Exported Nov 22, 2023 from SnapGene Viewer 7.0.3

https://www.snapgene.com

COMMENT	mCherry cloned into pVSV_XN2 from John Rose

FEATURES	Location/Qualifiers

source	1..15011

/mol_type="other DNA"

/organism="synthetic DNA construct"

rep_origin	16..456

/direction=RIGHT

/label=F1 ori

rep_origin	complement(33..461)

/direction=LEFT

/label=f1 ori

/note="f1 bacteriophage origin of replication; arrow

indicates direction of (+) strand synthesis"

CDS	complement(463..531)

/label=LacZ alpha

primer_bind	602..619

/label=M13-fwd

primer_bind	603..619

/label=M13 fwd

/note="common sequencing primer, one of multiple similar

variants"

promoter	626..644

/label=T7 promoter

/note="promoter for bacteriophage T7 RNA polymerase"

CDS	708..1976

/codon_start=1

/label=N

/translation="MSVTVKRIIDNTVVVPKLPANEDPVEYPADYFRKSKEIPLYINTT

KSLSDLRGYVYQGLKSGNVSIIHVNSYLYGALKDIRGKLDKDWSSFGINIGKAGDTIGI

FDLVSLKALDGVLPDGVSDASRTSADDKWLPLYLLGLYRVGRTQMPEYRKKLMDGLTNQ

CKMINEQFEPLVPEGRDIFDVWGNDSNYTKIVAAVDMFFHMFKKHECASFRYGTIVSRF

KDCAALATFGHLCKITGMSTEDVTTWILNREVADEMVQMMLPGQEIDKADSYMPYLIDF

GLSSKSPYSSVKNPAFHFWGQLTALLLRSTRARNARQPDDIEYTSLTTAGLLYAYAVGS

SADLAQQFCVGDNKYTPDDSTGGLTTNAPPQGRDVVEWLGWFEDQNRKPTPDMMQYAKR

AVMSLQGLREKTIGKYAKSEFDK"

CDS	2040..2837

/codon_start=1

/label=P

/translation="MDNLTKVREYLKSYSRLDQAVGEIDEIEAQRAEKSNYELFQEDGV

EEHTKPSYFQAADDSDTESEPEIEDNQGLYAPDPEAEQVEGFIQGPLDDYADEEVDVVF

TSDWKQPELESDEHGKTLRLTSPEGLSGEQKSQWLSTIKAVVQSAKYWNLAECTFEASG

EGVIMKERQITPDVYKVTPVMNTHPSQSEAVSDVWSLSKTSMTFQPKKASLQPLTISLD

ELFSSRGEFISVGGDGRMSHKEAILLGLRYKKLYNQARVKYSL"

CDS	2894..3583

/codon_start=1

/label=P

/translation="MSSLKKILGLKGKGKKSKKLGIAPPPYEEDTSMEYAPSAPIDKSY

FGVDEMDTYDPNQLRYEKFFFTVKMTVRSNRPFRTYSDVAAAVSHWDHMYIGMAGKRPF

YKILAFLGSSNLKATPAVLADQGQPEYHAHCEGRAYLPHRMGKTPPMLNVPEHFRRPFN

IGLYKGTIELTMTIYDDESLEAAPMIWDHFNSSKFSDFREKALMFGLIVEKKASGAWVL

DSIGHFK"

CDS	3722..5257

/codon_start=1

/product="vesicular stomatitis virus G glycoprotein"

/label=VSV-G

/note="Indiana strain"

/translation="MKCLLYLAFLFIGVNCKFTIVFPHNQKGNWKNVPSNYHYCPSSSD

LNWHNDLIGTALQVKMPKSHKAIQADGWMCHASKWVTTCDFRWYGPKYITHSIRSFTPS

VEQCKESIEQTKQGTWLNPGFPPQSCGYATVTDAEAVIVQVTPHHVLVDEYTGEWVDSQ

FINGKCSNYICPTVHNSTTWHSDYKVKGLCDSNLISMDITFFSEDGELSSLGKEGTGFR

SNYFAYETGGKACKMQYCKHWGVRLPSGVWFEMADKDLFAAARFPECPEGSSISAPSQT

SVDVSLIQDVERILDYSLCQETWSKIRAGLPISPVDLSYLAPKNPGTGPAFTIINGTLK

YFETRYIRVDIAAPILSRMVGMISGTTTERELWDDWAPYEDVEIGPNGVLRTSSGYKFP

LYMIGHGMLDSDLHLSSKAQVFEHPHIQDAASQLPDDESLFFGDTGLSKNPIELVEGWF

SSWKSSIASFFFIIGLIIGLFLVLRVGIHLCIKLKHTKKRQIYTDIEMNRLGK"

CDS	5309..6019

/codon_start=1

/product="monomeric derivative of DsRed fluorescent protein

(Shaner et al., 2004)"

/label=mCherry

/note="mammalian codon-optimized"

/translation="MVSKGEEDNMAIIKEFMRFKVHMEGSVNGHEFEIEGEGEGRPYEG

TQTAKLKVTKGGPLPFAWDILSPQFMYGSKAYVKHPADIPDYLKLSFPEGFKWERVMNF

EDGGVVTVTQDSSLQDGEFIYKVKLRGTNFPSDGPVMQKKTMGWEASSERMYPEDGALK

GEIKQRLKLKDGGHYDAEVKTTYKAKKPVQLPGAYNVNIKLDITSHNEDYTIVEQYERA

EGRHSTGGMDELYK"

CDS	6129..12458

/codon_start=1

/label=L

/translation="MEVHDFETDEFNDFNEDDYATREFLNPDERMTYLNHADYNLNSPL

ISDDIDNLIRKFNSLPIPSMWDSKNWDGVLEMLTSCQANPIPTSQMHKWMGSWLMSDNH

DASQGYSFLHEVDKEAEITFDVVETFIRGWGNKPIEYIKKERWTDSFKILAYLCQKFLD

LHKLTLILNAVSEVELLNLARTFKGKVRRSSHGTNICRIRVPSLGPTFISEGWAYFKKL

DILMDRNFLLMVKDVIIGRMQTVLSMVCRIDNLFSEQDIFSLLNIYRIGDKIVERQGNF

SYDLIKMVEPICNLKLMKLARESRPLVPQFPHFENHIKTSVDEGAKIDRGIRFLHDQIM

SVKTVDLTLVIYGSFRHWGHPFIDYYTGLEKLHSQVTMKKDIDVSYAKALASDLARIVL

FQQFNDHKKWFVNGDLLPHDHPFKSHVKENTWPTAAQVQDFGDKWHELPLIKCFEIPDL

LDPSIIYSDKSHSMNRSEVLKHVRMNPNTPIPSKKVLQTMLDTKATNWKEFLKEIDEKG

LDDDDLIIGLKGKERELKLAGRFFSLMSWKLREYFVITEYLIKTHFVPMFKGLTMADDL

TAVIKKMLDSSSGQGLKSYEAICIANHIDYEKWNNHQRKLSNGPVFRVMGQFLGYPSLI

ERTHEFFEKSLIYYNGRPDLMRVHNNTLINSTSQRVCWQGQEGGLEGLRQKGWSILNLL

VIQREAKIRNTAVKVLAQGDNQVICTQYKTKKSRNVVELQGALNQMVSNNEKIMTAIKI

GTGKLGLLINDDETMQSADYLNYGKIPIFRGVIRGLETKRWSRVTCVTNDQIPTCANIM

SSVSTNALTVAHFAENPINAMIQYNYFGTFARLLLMMHDPALRQSLYEVQDKIPGLHSS

TFKYAMLYLDPSIGGVSGMSLSRFLIRAFPDPVTESLSFWRFIHVHARSEHLKEMSAVF

GNPEIAKFRITHIDKLVEDPTSLNIAMGMSPANLLKTEVKKCLIESRQTIRNQVIKDAT

IYLYHEEDRLRSFLWSINPLFPRFLSEFKSGTFLGVADGLISLFQNSRTIRNSFKKKYH

RELDDLIVRSEVSSLTHLGKLHLRRGSCKMWTCSATHADTLRYKSWGRTVIGTTVPHPL

EMLGPQHRKETPCAPCNTSGFNYVSVHCPDGIHDVFSSRGPLPAYLGSKTSESTSILQP

WERESKVPLIKRATRLRDAISWFVEPDSKLAMTILSNIHSLTGEEWTKRQHGFKRTGSA

LHRFSTSRMSHGGFASQSTAALTRLMATTDTMRDLGDQNFDFLFQATLLYAQITTTVAR

DGWITSCTDHYHIACKSCLRPIEEITLDSSMDYTPPDVSHVLKTWRNGEGSWGQEIKQI

YPLEGNWKNLAPAEQSYQVGRCIGFLYGDLAYRKSTHAEDSSLFPLSIQGRIRGRGFLK

GLLDGLMRASCCQVIHRRSLAHLKRPANAVYGGLIYLIDKLSVSPPFLSLTRSGPIRDE

LETIPHKIPTSYPTSNRDMGVIVRNYFKYQCRLIEKGKYRSHYSQLWLFSDVLSIDFIG

PFSISTTLLQILYKPFLSGKDKNELRELANLSSLLRSGEGWEDIHVKFFTKDILLCPEE

IRHACKFGIAKDNNKDMSYPPWGRESRGTITTIPVYYTTTPYPKMLEMPPRIQNPLLSG

IRLGQLPTGAHYKIRSILHGMGIHYRDFLSCGDGSGGMTAALLRENVHSRGIFNSLLEL

SGSVMRGASPEPPSALETLGGDKSRCVNGETCWEYPSDLCDPRTWDYFLRLKAGLGLQI

DLIVMDMEVRDSSTSLKIETNVRNYVHRILDEQGVLIYKTYGTYICESEKNAVTILGPM

FKTVDLVQTEFSSSQTSEVYMVCKGLKKLIDEPNPDWSSINESWKNLYAFQSSEQEFAR

AKKVSTYFTLTGIPSQFIPDPFVNIETMLQIFGVPTGVSHAAALKSSDRPADLLTISLF

YMAIISYYNINHIRVGPIPPNPPSDGIAQNVGIAITGISFWLSLMEKDIPLYQQCLAVI

QQSFPIRWEAVSVKGGYKQKWSTRGDGLPKDTRISDSLAPIGNWIRSLELVRNQVRLNP

FNEILFNQLCRTVDNHLKWSNLRRNTGMIEWINRRISKEDRSILMLKSDLHEENSWRD"

terminator	12711..12758

/label=T7 terminator

/note="transcription terminator for bacteriophage T7 RNA

polymerase"

primer_bind	complement(12824..12843)

/label=T3

promoter	complement(12826..12844)

/label=T3 promoter

/note="promoter for bacteriophage T3 RNA polymerase"

primer_bind	complement(12861..12881)

/label=M13-rev

primer_bind	complement(12865..12881)

/label=M13 rev

/note="common sequencing primer, one of multiple similar

variants"

misc_binding	complement(12887..12909)

/label=LacO

protein_bind	12889..12905

/label=lac operator

/bound_moiety="lac repressor encoded by lacI"

/note="The lac repressor binds to the lac operator to

inhibit transcription in E. coli. This inhibition can be

relieved by adding lactose or

isopropyl-beta-D-thiogalactopyranoside (IPTG)."

promoter	complement(12913..12943)

/label=lac promoter

/note="promoter for the E. coli lac operon"

promoter	complement(12914..12943)

/label=lac

protein_bind	12958..12979

/label=CAP binding site

/bound_moiety="E. coli catabolite activator protein"

/note="CAP binding activates transcription in the presence

of cAMP."

rep_origin	complement(13249..13877)

/direction=LEFT

/label=ColE1 origin

rep_origin	complement(13267..13855)

/direction=LEFT

/label=ori

/note="high-copy-number ColE1/pMB1/pBR322/pUC origin of

replication"

CDS	complement(14026..14886)

/codon_start=1

/gene="bla"

/product="beta-lactamase"

/label=AmpR

/note="confers resistance to ampicillin, carbenicillin, and

related antibiotics"

/translation="MSIQHFRVALIPFFAAFCLPVFAHPETLVKVKDAEDQLGARVGYI

ELDLNSGKILESFRPEERFPMMSTFKVLLCGAVLSRIDAGQEQLGRRIHYSQNDLVEYS

PVTEKHLTDGMTVRELCSAAITMSDNTAANLLLTTIGGPKELTAFLHNMGDHVTRLDRW

EPELNEAIPNDERDTTMPVAMATTLRKLLTGELLTLASRQQLIDWMEADKVAGPLLRSA

LPAGWFIADKSGAGERGSRGIIAALGPDGKPSRIVVIYTTGSQATMDERNRQIAEIGAS

LIKHW"

CDS	complement(14029..14688)

/label=AmpR

promoter	complement(14887..14991)

/gene="bla"

/label=AmpR promoter

promoter	complement(14928..14956)

/label=Amp prom

ORIGIN

1 cacctaaatt gtaagcgtta atattttgtt aaaattcgcg ttaaattttt gttaaatcag

61 ctcatttttt aaccaatagg ccgaaatcgg caaaatccct tataaatcaa aagaatagac

121 cgagataggg ttgagtgttg ttccagtttg gaacaagagt ccactattaa agaacgtgga

181 ctccaacgtc aaagggcgaa aaaccgtcta tcagggcgat ggcccactac gtgaaccatc

241 accctaatca agttttttgg ggtcgaggtg ccgtaaagca ctaaatcgga accctaaagg

301 gagcccccga tttagagctt gacggggaaa gccggcgaac gtggcgagaa aggaagggaa

361 gaaagcgaaa ggagcgggcg ctagggcgct ggcaagtgta gcggtcacgc tgcgcgtaac

421 caccacaccc gccgcgctta atgcgccgct acagggcgcg tcccattcgc cattcaggct

481 gcgcaactgt tgggaagggc gatcggtgcg ggcctcttcg ctattacgcc agctggcgaa

541 agggggatgt gctgcaaggc gattaagttg ggtaacgcca gggttttccc agtcacgacg

601 ttgtaaaacg acggccagtg aattgtaata cgactcacta taggacgaag acaaacaaac

661 cattattatc attaaaaggc tcaggagaaa ctttaacagt aatcaaaatg tctgttacag

721 tcaagagaat cattgacaac acagtcgtag ttccaaaact tcctgcaaat gaggatccag

781 tggaataccc ggcagattac ttcagaaaat caaaggagat tcctctttac atcaatacta

841 caaaaagttt gtcagatcta agaggatatg tctaccaagg cctcaaatcc ggaaatgtat

901 caatcataca tgtcaacagc tacttgtatg gagcattaaa ggacatccgg ggtaagttgg

961 ataaagattg gtcaagtttc ggaataaaca tcgggaaagc aggggataca atcggaatat

1021 ttgaccttgt atccttgaaa gccctggacg gcgtacttcc agatggagta tcggatgctt

1081 ccagaaccag cgcagatgac aaatggttgc ctttgtatct acttggctta tacagagtgg

1141 gcagaacaca aatgcctgaa tacagaaaaa agctcatgga tgggctgaca aatcaatgca

1201 aaatgatcaa tgaacagttt gaacctcttg tgccagaagg tcgtgacatt tttgatgtgt

1261 ggggaaatga cagtaattac acaaaaattg tcgctgcagt ggacatgttc ttccacatgt

1321 tcaaaaaaca tgaatgtgcc tcgttcagat acggaactat tgtttccaga ttcaaagatt

1381 gtgctgcatt ggcaacattt ggacacctct gcaaaataac cggaatgtct acagaagatg

1441 taacgacctg gatcttgaac cgagaagttg cagatgaaat ggtccaaatg atgcttccag

1501 gccaagaaat tgacaaggcc gattcataca tgccttattt gatcgacttt ggattgtctt

1561 ctaagtctcc atattcttcc gtcaaaaacc ctgccttcca cttctggggg caattgacag

1621 ctcttctgct cagatccacc agagcaagga atgcccgaca gcctgatgac attgagtata

1681 catctcttac tacagcaggt ttgttgtacg cttatgcagt aggatcctct gccgacttgg

1741 cacaacagtt ttgtgttgga gataacaaat acactccaga tgatagtacc ggaggattga

1801 cgactaatgc accgccacaa ggcagagatg tggtcgaatg gctcggatgg tttgaagatc

1861 aaaacagaaa accgactcct gatatgatgc agtatgcgaa aagagcagtc atgtcactgc

1921 aaggcctaag agagaagaca attggcaagt atgctaagtc agaatttgac aaatgaccct

1981 ataattctca gatcacctat tatatattat gctacatatg aaaaaaacta acagatatca

2041 tggataatct cacaaaagtt cgtgagtatc tcaagtccta ttctcgtctg gatcaggcgg

2101 taggagagat agatgagatc gaagcacaac gagctgaaaa gtccaattat gagttgttcc

2161 aagaggatgg agtggaagag catactaagc cctcttattt tcaggcagca gatgattctg

2221 acacagaatc tgaaccagaa attgaagaca atcaaggctt gtatgcacca gatccagaag

2281 ctgagcaagt tgaaggcttt atacaggggc ctttagatga ctatgcagat gaggaagtgg

2341 atgttgtatt tacttcggac tggaaacagc ctgagcttga atctgacgag catggaaaga

2401 ccttacggtt gacatcgcca gagggtttaa gtggagagca gaaatcccag tggctttcga

2461 cgattaaagc agtcgtgcaa agtgccaaat actggaatct ggcagagtgc acatttgaag

2521 catcgggaga aggggtcatt atgaaggagc gccagataac tccggatgta tataaggtca

2581 ctccagtgat gaacacacat ccgtcccaat cagaagcagt atcagatgtt tggtctctct

2641 caaagacatc catgactttc caacccaaga aagcaagtct tcagcctctc accatatcct

2701 tggatgaatt gttctcatct agaggagagt tcatctctgt cggaggtgac ggacgaatgt

2761 ctcataaaga ggccatcctg ctcggcctga gatacaaaaa gttgtacaat caggcgagag

2821 tcaaatattc tctgtagact atgaaaaaaa gtaacagata tcacgatcta agtgttatcc

2881 caatccattc atcatgagtt ccttaaagaa gattctcggt ctgaagggga aaggtaagaa

2941 atctaagaaa ttagggatcg caccaccccc ttatgaagag gacactagca tggagtatgc

3001 tccgagcgct ccaattgaca aatcctattt tggagttgac gagatggaca cctatgatcc

3061 gaatcaatta agatatgaga aattcttctt tacagtgaaa atgacggtta gatctaatcg

3121 tccgttcaga acatactcag atgtggcagc cgctgtatcc cattgggatc acatgtacat

3181 cggaatggca gggaaacgtc ccttctacaa aatcttggct tttttgggtt cttctaatct

3241 aaaggccact ccagcggtat tggcagatca aggtcaacca gagtatcacg ctcactgcga

3301 aggcagggct tatttgccac ataggatggg gaagacccct cccatgctca atgtaccaga

3361 gcacttcaga agaccattca atataggtct ttacaaggga acgattgagc tcacaatgac

3421 catctacgat gatgagtcac tggaagcagc tcctatgatc tgggatcatt tcaattcttc

3481 caaattttct gatttcagag agaaggcctt aatgtttggc ctgattgtcg agaaaaaggc

3541 atctggagcg tgggtcctgg actctatcgg ccacttcaaa tgagctagtc taacttctag

3601 cttctgaaca atccccggtt tactcagtct cccctaattc cagcctctcg aacaactaat

3661 atcctgtctt ttctatccct atgaaaaaaa ctaacagaga tcgatctgtt tacgcgtcac

3721 tatgaagtgc cttttgtact tagccttttt attcattggg gtgaattgca agttcaccat

3781 agtttttcca cacaaccaaa aaggaaactg gaaaaatgtt ccttctaatt accattattg

3841 cccgtcaagc tcagatttaa attggcataa tgacttaata ggcacagcct tacaagtcaa

3901 aatgcccaag agtcacaagg ctattcaagc agacggttgg atgtgtcatg cttccaaatg

3961 ggtcactact tgtgatttcc gctggtatgg accgaagtat ataacacatt ccatccgatc

4021 cttcactcca tctgtagaac aatgcaagga aagcattgaa caaacgaaac aaggaacttg

4081 gctgaatcca ggcttccctc ctcaaagttg tggatatgca actgtgacgg atgccgaagc

4141 agtgattgtc caggtgactc ctcaccatgt gctggttgat gaatacacag gagaatgggt

4201 tgattcacag ttcatcaacg gaaaatgcag caattacata tgccccactg tccataactc

4261 tacaacctgg cattctgact ataaggtcaa agggctatgt gattctaacc tcatttccat

4321 ggacatcacc ttcttctcag aggacggaga gctatcatcc ctgggaaagg agggcacagg

4381 gttcagaagt aactactttg cttatgaaac tggaggcaag gcctgcaaaa tgcaatactg

4441 caagcattgg ggagtcagac tcccatcagg tgtctggttc gagatggctg ataaggatct

4501 ctttgctgca gccagattcc ctgaatgccc agaagggtca agtatctctg ctccatctca

4561 gacctcagtg gatgtaagtc taattcagga cgttgagagg atcttggatt attccctctg

4621 ccaagaaacc tggagcaaaa tcagagcggg tcttccaatc tctccagtgg atctcagcta

4681 tcttgctcct aaaaacccag gaaccggtcc tgctttcacc ataatcaatg gtaccctaaa

4741 atactttgag accagataca tcagagtcga tattgctgct ccaatcctct caagaatggt

4801 cggaatgatc agtggaacta ccacagaaag ggaactgtgg gatgactggg caccatatga

4861 agacgtggaa attggaccca atggagttct gaggaccagt tcaggatata agtttccttt

4921 atacatgatt ggacatggta tgttggactc cgatcttcat cttagctcaa aggctcaggt

4981 gttcgaacat cctcacattc aagacgctgc ttcgcaactt cctgatgatg agagtttatt

5041 ttttggtgat actgggctat ccaaaaatcc aatcgagctt gtagaaggtt ggttcagtag

5101 ttggaaaagc tctattgcct cttttttctt tatcataggg ttaatcattg gactattctt

5161 ggttctccga gttggtatcc atctttgcat taaattaaag cacaccaaga aaagacagat

5221 ttatacagac atagagatga accgacttgg aaagtaactc aaatcctgct aggtatgaaa

5281 aaaactaaca gatatcacgc tcgagaccat ggtgagcaag ggcgaggagg ataacatggc

5341 catcatcaag gagttcatgc gcttcaaggt gcacatggag ggctccgtga acggccacga

5401 gttcgagatc gagggcgagg gcgagggccg cccctacgag ggcacccaga ccgccaagct

5461 gaaggtgacc aagggtggcc ccctgccctt cgcctgggac atcctgtccc ctcagttcat

5521 gtacggctcc aaggcctacg tgaagcaccc cgccgacatc cccgactact tgaagctgtc

5581 cttccccgag ggcttcaagt gggagcgcgt gatgaacttc gaggacggcg gcgtggtgac

5641 cgtgacccag gactcctccc tgcaggacgg cgagttcatc tacaaggtga agctgcgcgg

5701 caccaacttc ccctccgacg gccccgtaat gcagaagaag accatgggct gggaggcctc

5761 ctccgagcgg atgtaccccg aggacggcgc cctgaagggc gagatcaagc agaggctgaa

5821 gctgaaggac ggcggccact acgacgctga ggtcaagacc acctacaagg ccaagaagcc

5881 cgtgcagctg cccggcgcct acaacgtcaa catcaagttg gacatcacct cccacaacga

5941 ggactacacc atcgtggaac agtacgaacg cgccgagggc cgccactcca ccggcggcat

6001 ggacgagctg tacaagtaag ctagccagat tcttcatgtt tggaccaaat caacttgtga

6061 taccatgctc aaagaggcct caattatatt tgagttttta atttttatga aaaaaactaa

6121 cagcaatcat ggaagtccac gattttgaga ccgacgagtt caatgatttc aatgaagatg

6181 actatgccac aagagaattc ctgaatcccg atgagcgcat gacgtacttg aatcatgctg

6241 attacaacct gaattctcct ctaattagtg atgatattga caatttaatc aggaaattca

6301 attctcttcc aattccctcg atgtgggata gtaagaactg ggatggagtt cttgagatgt

6361 taacgtcatg tcaagccaat cccatcccaa catctcagat gcataaatgg atgggaagtt

6421 ggttaatgtc tgataatcat gatgccagtc aagggtatag ttttttacat gaagtggaca

6481 aagaggcaga aataacattt gacgtggtgg agaccttcat ccgcggctgg ggcaacaaac

6541 caattgaata catcaaaaag gaaagatgga ctgactcatt caaaattctc gcttatttgt

6601 gtcaaaagtt tttggactta cacaagttga cattaatctt aaatgctgtc tctgaggtgg

6661 aattgctcaa cttggcgagg actttcaaag gcaaagtcag aagaagttct catggaacga

6721 acatatgcag gattagggtt cccagcttgg gtcctacttt tatttcagaa ggatgggctt

6781 acttcaagaa acttgatatt ctaatggacc gaaactttct gttaatggtc aaagatgtga

6841 ttatagggag gatgcaaacg gtgctatcca tggtatgtag aatagacaac ctgttctcag

6901 agcaagacat cttctccctt ctaaatatct acagaattgg agataaaatt gtggagaggc

6961 agggaaattt ttcttatgac ttgattaaaa tggtggaacc gatatgcaac ttgaagctga

7021 tgaaattagc aagagaatca aggcctttag tcccacaatt ccctcatttt gaaaatcata

7081 tcaagacttc tgttgatgaa ggggcaaaaa ttgaccgagg tataagattc ctccatgatc

7141 agataatgag tgtgaaaaca gtggatctca cactggtgat ttatggatcg ttcagacatt

7201 ggggtcatcc ttttatagat tattacactg gactagaaaa attacattcc caagtaacca

7261 tgaagaaaga tattgatgtg tcatatgcaa aagcacttgc aagtgattta gctcggattg

7321 ttctatttca acagttcaat gatcataaaa agtggttcgt gaatggagac ttgctccctc

7381 atgatcatcc ctttaaaagt catgttaaag aaaatacatg gcccacagct gctcaagttc

7441 aagattttgg agataaatgg catgaacttc cgctgattaa atgttttgaa atacccgact

7501 tactagaccc atcgataata tactctgaca aaagtcattc aatgaatagg tcagaggtgt

7561 tgaaacatgt ccgaatgaat ccgaacactc ctatccctag taaaaaggtg ttgcagacta

7621 tgttggacac aaaggctacc aattggaaag aatttcttaa agagattgat gagaagggct

7681 tagatgatga tgatctaatt attggtctta aaggaaagga gagggaactg aagttggcag

7741 gtagattttt ctccctaatg tcttggaaat tgcgagaata ctttgtaatt accgaatatt

7801 tgataaagac tcatttcgtc cctatgttta aaggcctgac aatggcggac gatctaactg

7861 cagtcattaa aaagatgtta gattcctcat ccggccaagg attgaagtca tatgaggcaa

7921 tttgcatagc caatcacatt gattacgaaa aatggaataa ccaccaaagg aagttatcaa

7981 acggcccagt gttccgagtt atgggccagt tcttaggtta tccatcctta atcgagagaa

8041 ctcatgaatt ttttgagaaa agtcttatat actacaatgg aagaccagac ttgatgcgtg

8101 ttcacaacaa cacactgatc aattcaacct cccaacgagt ttgttggcaa ggacaagagg

8161 gtggactgga aggtctacgg caaaaaggat ggagtatcct caatctactg gttattcaaa

8221 gagaggctaa aatcagaaac actgctgtca aagtcttggc acaaggtgat aatcaagtta

8281 tttgcacaca gtataaaacg aagaaatcga gaaacgttgt agaattacag ggtgctctca

8341 atcaaatggt ttctaataat gagaaaatta tgactgcaat caaaataggg acagggaagt

8401 taggactttt gataaatgac gatgagacta tgcaatctgc agattacttg aattatggaa

8461 aaataccgat tttccgtgga gtgattagag ggttagagac caagagatgg tcacgagtga

8521 cttgtgtcac caatgaccaa atacccactt gtgctaatat aatgagctca gtttccacaa

8581 atgctctcac cgtagctcat tttgctgaga acccaatcaa tgccatgata cagtacaatt

8641 attttgggac atttgctaga ctcttgttga tgatgcatga tcctgctctt cgtcaatcat

8701 tgtatgaagt tcaagataag ataccgggct tgcacagttc tactttcaaa tacgccatgt

8761 tgtatttgga cccttccatt ggaggagtgt cgggcatgtc tttgtccagg tttttgatta

8821 gagccttccc agatcccgta acagaaagtc tctcattctg gagattcatc catgtacatg

8881 ctcgaagtga gcatctgaag gagatgagtg cagtatttgg aaaccccgag atagccaagt

8941 ttcgaataac tcacatagac aagctagtag aagatccaac ctctctgaac atcgctatgg

9001 gaatgagtcc agcgaacttg ttaaagactg aggttaaaaa atgcttaatc gaatcaagac

9061 aaaccatcag gaaccaggtg attaaggatg caaccatata tttgtatcat gaagaggatc

9121 ggctcagaag tttcttatgg tcaataaatc ctctgttccc tagattttta agtgaattca

9181 aatcaggcac ttttttggga gtcgcagacg ggctcatcag tctatttcaa aattctcgta

9241 ctattcggaa ctcctttaag aaaaagtatc atagggaatt ggatgatttg attgtgagga

9301 gtgaggtatc ctctttgaca catttaggga aacttcattt gagaagggga tcatgtaaaa

9361 tgtggacatg ttcagctact catgctgaca cattaagata caaatcctgg ggccgtacag

9421 ttattgggac aactgtaccc catccattag aaatgttggg tccacaacat cgaaaagaga

9481 ctccttgtgc accatgtaac acatcagggt tcaattatgt ttctgtgcat tgtccagacg

9541 ggatccatga cgtctttagt tcacggggac cattgcctgc ttatctaggg tctaaaacat

9601 ctgaatctac atctattttg cagccttggg aaagggaaag caaagtccca ctgattaaaa

9661 gagctacacg tcttagagat gctatctctt ggtttgttga acccgactct aaactagcaa

9721 tgactatact ttctaacatc cactctttaa caggcgaaga atggaccaaa aggcagcatg

9781 ggttcaaaag aacagggtct gcccttcata ggttttcgac atctcggatg agccatggtg

9841 ggttcgcatc tcagagcact gcagcattga ccaggttgat ggcaactaca gacaccatga

9901 gggatctggg agatcagaat ttcgactttt tattccaagc aacgttgctc tatgctcaaa

9961 ttaccaccac tgttgcaaga gacggatgga tcaccagttg tacagatcat tatcatattg

10021 cctgtaagtc ctgtttgaga cccatagaag agatcaccct ggactcaagt atggactaca

10081 cgcccccaga tgtatcccat gtgctgaaga catggaggaa tggggaaggt tcgtggggac

10141 aagagataaa acagatctat cctttagaag ggaattggaa gaatttagca cctgctgagc

10201 aatcctatca agtcggcaga tgtataggtt ttctatatgg agacttggcg tatagaaaat

10261 ctactcatgc cgaggacagt tctctatttc ctctatctat acaaggtcgt attagaggtc

10321 gaggtttctt aaaagggttg ctagacggat taatgagagc aagttgctgc caagtaatac

10381 accggagaag tctggctcat ttgaagaggc cggccaacgc agtgtacgga ggtttgattt

10441 acttgattga taaattgagt gtatcacctc cattcctttc tcttactaga tcaggaccta

10501 ttagagacga attagaaacg attccccaca agatcccaac ctcctatccg acaagcaacc

10561 gtgatatggg ggtgattgtc agaaattact tcaaatacca atgccgtcta attgaaaagg

10621 gaaaatacag atcacattat tcacaattat ggttattctc agatgtctta tccatagact

10681 tcattggacc attctctatt tccaccaccc tcttgcaaat cctatacaag ccatttttat

10741 ctgggaaaga taagaatgag ttgagagagc tggcaaatct ttcttcattg ctaagatcag

10801 gagaggggtg ggaagacata catgtgaaat tcttcaccaa ggacatatta ttgtgtccag

10861 aggaaatcag acatgcttgc aagttcggga ttgctaagga taataataaa gacatgagct

10921 atcccccttg gggaagggaa tccagaggga caattacaac aatccctgtt tattatacga

10981 ccacccctta cccaaagatg ctagagatgc ctccaagaat ccaaaatccc ctgctgtccg

11041 gaatcaggtt gggccaatta ccaactggcg ctcattataa aattcggagt atattacatg

11101 gaatgggaat ccattacagg gacttcttga gttgtggaga cggctccgga gggatgactg

11161 ctgcattact acgagaaaat gtgcatagca gaggaatatt caatagtctg ttagaattat

11221 cagggtcagt catgcgaggc gcctctcctg agccccccag tgccctagaa actttaggag

11281 gagataaatc gagatgtgta aatggtgaaa catgttggga atatccatct gacttatgtg

11341 acccaaggac ttgggactat ttcctccgac tcaaagcagg cttggggctt caaattgatt

11401 taattgtaat ggatatggaa gttcgggatt cttctactag cctgaaaatt gagacgaatg

11461 ttagaaatta tgtgcaccgg attttggatg agcaaggagt tttaatctac aagacttatg

11521 gaacatatat ttgtgagagc gaaaagaatg cagtaacaat ccttggtccc atgttcaaga

11581 cggtcgactt agttcaaaca gaatttagta gttctcaaac gtctgaagta tatatggtat

11641 gtaaaggttt gaagaaatta atcgatgaac ccaatcccga ttggtcttcc atcaatgaat

11701 cctggaaaaa cctgtacgca ttccagtcat cagaacagga atttgccaga gcaaagaagg

11761 ttagtacata ctttaccttg acaggtattc cctcccaatt cattcctgat ccttttgtaa

11821 acattgagac tatgctacaa atattcggag tacccacggg tgtgtctcat gcggctgcct

11881 taaaatcatc tgatagacct gcagatttat tgaccattag ccttttttat atggcgatta

11941 tatcgtatta taacatcaat catatcagag taggaccgat acctccgaac cccccatcag

12001 atggaattgc acaaaatgtg gggatcgcta taactggtat aagcttttgg ctgagtttga

12061 tggagaaaga cattccacta tatcaacagt gtttagcagt tatccagcaa tcattcccga

12121 ttaggtggga ggctgtttca gtaaaaggag gatacaagca gaagtggagt actagaggtg

12181 atgggctccc aaaagatacc cgaatttcag actccttggc cccaatcggg aactggatca

12241 gatctctgga attggtccga aaccaagttc gtctaaatcc attcaatgag atcttgttca

12301 atcagctatg tcgtacagtg gataatcatt tgaaatggtc aaatttgcga agaaacacag

12361 gaatgattga atggatcaat agacgaattt caaaagaaga ccggtctata ctgatgttga

12421 agagtgacct acacgaggaa aactcttgga gagattaaaa aatcatgagg agactccaaa

12481 ctttaagtat gaaaaaaact ttgatcctta agaccctctt gtggttttta ttttttatct

12541 ggttttgtgg tcttcgtggg tcggcatggc atctccacct cctcgcggtc cgacctgggc

12601 atccgaagga ggacgtcgtc cactcggatg gctaagggag gggcccccgc ggggctgcta

12661 acaaagcccg aaaggaagct gagttggctg ctgccaccgc tgagcaataa ctagcataac

12721 cccttggggc ctctaaacgg gtcttgaggg gttttttgct gaaaggagga actatatccg

12781 gatcgagacc tcgatactag tgcggtggag ctccagcttt tgttcccttt agtgagggtt

12841 aatttcgagc ttggcgtaat catggtcata gctgtttcct gtgtgaaatt gttatccgct

12901 cacaattcca cacaacatac gagccggaag cataaagtgt aaagcctggg gtgcctaatg

12961 agtgagctaa ctcacattaa ttgcgttgcg ctcactgccc gctttccagt cgggaaacct

13021 gtcgtgccag ctgcattaat gaatcggcca acgcgcgggg agaggcggtt tgcgtattgg

13081 gcgctcttcc gcttcctcgc tcactgactc gctgcgctcg gtcgttcggc tgcggcgagc

13141 ggtatcagct cactcaaagg cggtaatacg gttatccaca gaatcagggg ataacgcagg

13201 aaagaacatg tgagcaaaag gccagcaaaa ggccaggaac cgtaaaaagg ccgcgttgct

13261 ggcgtttttc cataggctcc gcccccctga cgagcatcac aaaaatcgac gctcaagtca

13321 gaggtggcga aacccgacag gactataaag ataccaggcg tttccccctg gaagctccct

13381 cgtgcgctct cctgttccga ccctgccgct taccggatac ctgtccgcct ttctcccttc

13441 gggaagcgtg gcgctttctc atagctcacg ctgtaggtat ctcagttcgg tgtaggtcgt

13501 tcgctccaag ctgggctgtg tgcacgaacc ccccgttcag cccgaccgct gcgccttatc

13561 cggtaactat cgtcttgagt ccaacccggt aagacacgac ttatcgccac tggcagcagc

13621 cactggtaac aggattagca gagcgaggta tgtaggcggt gctacagagt tcttgaagtg

13681 gtggcctaac tacggctaca ctagaaggac agtatttggt atctgcgctc tgctgaagcc

13741 agttaccttc ggaaaaagag ttggtagctc ttgatccggc aaacaaacca ccgctggtag

13801 cggtggtttt tttgtttgca agcagcagat tacgcgcaga aaaaaaggat ctcaagaaga

13861 tcctttgatc ttttctacgg ggtctgacgc tcagtggaac gaaaactcac gttaagggat

13921 tttggtcatg agattatcaa aaaggatctt cacctagatc cttttaaatt aaaaatgaag

13981 ttttaaatca atctaaagta tatatgagta aacttggtct gacagttacc aatgcttaat

14041 cagtgaggca cctatctcag cgatctgtct atttcgttca tccatagttg cctgactccc

14101 cgtcgtgtag ataactacga tacgggaggg cttaccatct ggccccagtg ctgcaatgat

14161 accgcgagac ccacgctcac cggctccaga tttatcagca ataaaccagc cagccggaag

14221 ggccgagcgc agaagtggtc ctgcaacttt atccgcctcc atccagtcta ttaattgttg

14281 ccgggaagct agagtaagta gttcgccagt taatagtttg cgcaacgttg ttgccattgc

14341 tacaggcatc gtggtgtcac gctcgtcgtt tggtatggct tcattcagct ccggttccca

14401 acgatcaagg cgagttacat gatcccccat gttgtgcaaa aaagcggtta gctccttcgg

14461 tcctccgatc gttgtcagaa gtaagttggc cgcagtgtta tcactcatgg ttatggcagc

14521 actgcataat tctcttactg tcatgccatc cgtaagatgc ttttctgtga ctggtgagta

14581 ctcaaccaag tcattctgag aatagtgtat gcggcgaccg agttgctctt gcccggcgtc

14641 aatacgggat aataccgcgc cacatagcag aactttaaaa gtgctcatca ttggaaaacg

14701 ttcttcgggg cgaaaactct caaggatctt accgctgttg agatccagtt cgatgtaacc

14761 cactcgtgca cccaactgat cttcagcatc ttttactttc accagcgttt ctgggtgagc

14821 aaaaacagga aggcaaaatg ccgcaaaaaa gggaataagg gcgacacgga aatgttgaat

14881 actcatactc ttcctttttc aatattattg aagcatttat cagggttatt gtctcatgag

14941 cggatacata tttgaatgta tttagaaaaa taaacaaata ggggttccgc gcacatttcc

15001 ccgaaaagtg c

//
