## Supplementary material for "An efficient plasmid-based system for the recovery of recombinant vesicular stomatitis virus encoding foreign glycoproteins": Data_S2_recovery protocol

**Supplementary Data 2**

Protocol for plasmid-based recovery of recombinant VSV.

Day 1.

Plate 0.15 x10^6^ BHK-G43 or BHK21 cells in a **12-well plate** in DMEM 1x + **FBS 5%** (w/o antibiotics). Incubate at 37 ºC for 24 hours.

Day 2.

A. Prepare DNA mix according to the tables below (depending on the cell line used). Each calculation is for a single well to be transfected. Add the indicated amount of DNA to 50ul of Opti-MEM with 2ul of P3000 reagent from the Lipofectamine 3000 kit. Incubate for 5 minutes at room temperature.

If using BHK-G43 cells:

| Plasmid | kbp | fmol DNA | ng DNA |
| --- | --- | --- | --- |
| pCMV-P | 6.3 | 25 | 97 |
| pCMV-N | 6.8 | 75 | 313 |
| pCMV-L | 11.8 | 25 | 182 |
| pCAGGS-T7opt | 7.4 | 50 | 228 |
| pVSV antigenomic | Varies | 25 | Varies by size |

If using standard BHK-21 cells:

| Plasmid | kbp | fmol DNA | ng DNA |
| --- | --- | --- | --- |
| pCMV-P | 6.3 | 25 | 97 |
| pCMV-N | 6.8 | 50 | 208 |
| pCMV-L | 11.8 | 25 | 182 |
| pCAGGS-T7opt | 7.4 | 50 | 228 |
| pVSV-G (pMD2.G) | 5.8 | 25 | 90 |
| pVSV antigenomic | Varies | 25 | Varies by size |

B. For each sample, prepare the Lipofectamine 3000 mixture by gently mixing the Lipofectamine 3000 reagent and then adding 2 ul of Lipofectamine 3000 to 50ul Opti-MEM. Incubate for 5 minutes at room temperature.

C. Add the DNA mixture to the diluted Lipofectamine 3000 and mix gently. Incubate for 15 minutes at room temperature. Meanwhile, wash cells with PBS 1x, then add 200ul of Opti-MEM and return to the incubator.

D. Add 100ul of transfection mixture to each well dropwise. Incubate cells at 37°C for 3 hours in the incubator.

E. Finally, add 1mL of DMEM + 10% FBS. For BHK-G43 cells, include 10nM of mifepristone to induce G expression. Incubate cells at 33°C for 40 hours.

Day 4.

Move cells to 37°C and incubate for an additional 36-48 hours. Examine cells for GFP expression to monitor recovery efficiency.

Day 6.

Collect supernatant, centrifuge 500 x g for 5 minutes to pellet cell debris, aliquot, and freeze at -80°C.

Notes:

1. Virus production can be quantified by performing 10-fold serial dilutions, infecting cells in a 96-well plate, and counting the number of cells expressing the fluorescent protein encoded in the antigenome at 8-9 hours post-infection.
2. The recombinant virus will be coated with the VSV G protein if recovered in mifepristone-treated BHK-G43 and therefore will also enter cells using this glycoprotein. Therefore, it is recommended to grow the virus in cells that do not express the G protein. For this, infect cells at a multiplicity of infection of <0.1 for at least one cycle.
3. If there is a need to increase viral titer at any point, it is possible to infect BHK-G43 cells induced to express G with mifepristone.
4. Incubation at 30°C is not essential and the virus can be recovered efficiently at 37°C. However, we find that the lower temperature improves recovery in our hands.
